# NGS-RFA/FRA: A High Throughput Experimental and Computational Pipeline for Selective Mutational Scanning in Parallel

**DOI:** 10.64898/2026.09.01.748481

**Authors:** Sina Tureli, Theo Bestebroer, Sarah James, Rachel Scheuer, Randall Dahn, Shufang Fan, Sam Turner, Sam Wilks, Antonia Netzl, A. Mosterin Hopping, Terry C. Jones, Gabriele Neumann, Yoshihiro Kawaoka, Ron Fouchier, Mathilde Richard, Derek J. Smith

## Abstract

Influenza viruses evade vaccine and infection mediated immunity by accumulating mutations in their hemagglutinin (HA) protein. Predicting this evolution might be possible via selective mutational scanning (SMS) – the generation of many specific mutants of interest from currently circulating viruses and characterizing their escape potential and fitness with high accuracy (Mögling 2016). However, this task is challenging, even when focusing on a reduced set of key HA positions (Koel et al. 2013). Here we describe a high-throughput SMS method to address this challenge. Our approach consists of a three-stage pipeline: (1) a parallel optimized virus rescue process that generates balanced target mutant virus libraries (2) an assay to assess replicative fitness and neutralisation of these variants as a mixture, and (3) a bespoke statistical model to quantify statistically significant differences between these observables. We tested the pipeline on libraries of up to 134 variants finding excellent correlation to classical hemagglutination inhibition (HI) and plaque growth assays used to assess antigenic phenotype and replicative fitness respectively, as well as remarkable repeatability overall. Notably, the method reduces the timeline required to carry out such assessments with classical methods from about a year to several weeks. By enabling rapid and efficient characterization of influenza virus variants, this approach has the potential to greatly enhance surveillance efforts, transforming reactive monitoring into proactive forecasting.

## INTRODUCTION

Influenza is an acute viral respiratory disease caused by influenza A, B, C, or D viruses. Type A and B viruses cause annual seasonal epidemics in humans and type A viruses can also cause unpredictable pandemics such as those in 1918, 1957, 1968, 2009 (Krammer et al. 2018). With an annual infection rate of 5%-15% causing an estimated 1 billion cases and 290,000 - 650,000 respiratory-related deaths, influenza represents a significant global health burden (Iuliano et al. 2018).

The standard management strategy against influenza, especially for at-risk groups, is annually administered vaccines with updates made as needed. Current vaccines protect by eliciting an antibody-mediated response against the immunodominant parts of the constituent vaccine strains. However, in the face of evolving host immunity, influenza virus persists by employing a swarm optimization strategy: With a high propensity to evolve, it can diffuse through a large space of variants and escape existing immunity by accumulating mutations in its hemagglutinin (HA) and neuraminidase (NA) proteins, undergoing antigenic evolution (Han et al. 2023) and individuals are reinfected multiple times during their lifetime (Hay et al. 2024). Even with annual vaccine updates, antigenic evolution and reliance on vaccine production after emergence of variants can cause low vaccine efficacy due to strain mismatch (Tenforde et al. 2021; Krammer et al. 2018; Boni 2008)

Hemagglutination inhibition (HI) titers or virus neutralisation (VN) titers are considered the “gold standards” for antigenic characterization of HA of evolving viruses. For instance, HI titrations were used to make antigenic maps which showed that H3N2 viruses, a subtype of influenza A virus, have formed ten distinct antigenic clusters between 1968-2002, each of which had significantly lowered HI titers against sera from the predecessor clusters (Smith et al. 2004). It was later that only seven positions, 145, 155, 156, 158, 159, 189, 193 in HA (the ‘Koel seven”) determine antigenic characteristics of influenza up to first order (Fig.1) (Koel et al. 2013). Since antigenic evolution of influenza virus often impacts regions around the receptor binding site (RBS), it can alter receptor binding properties and introduces replicative fitness as another critical parameter to the tug-of-war between evolution of the virus and adaptation of the immune system. Indeed, it was shown in (Mögling 2016) that immune escape substitutions negatively alter replicative fitness. This was demonstrated using 134 single amino acid substitution variants of the Koel-7 positions generated on the two root viruses A/Beijing/353/1989 (BE89 cluster) and A/Beijing/32/1992 (BE92 cluster). HI assays using multiple ferret sera from nearby clusters were performed on the full set of variants to assess antigenic phenotype and plaque radius growth was measured on thirty variants as a proxy of replicative fitness.

**Fig. 1.**
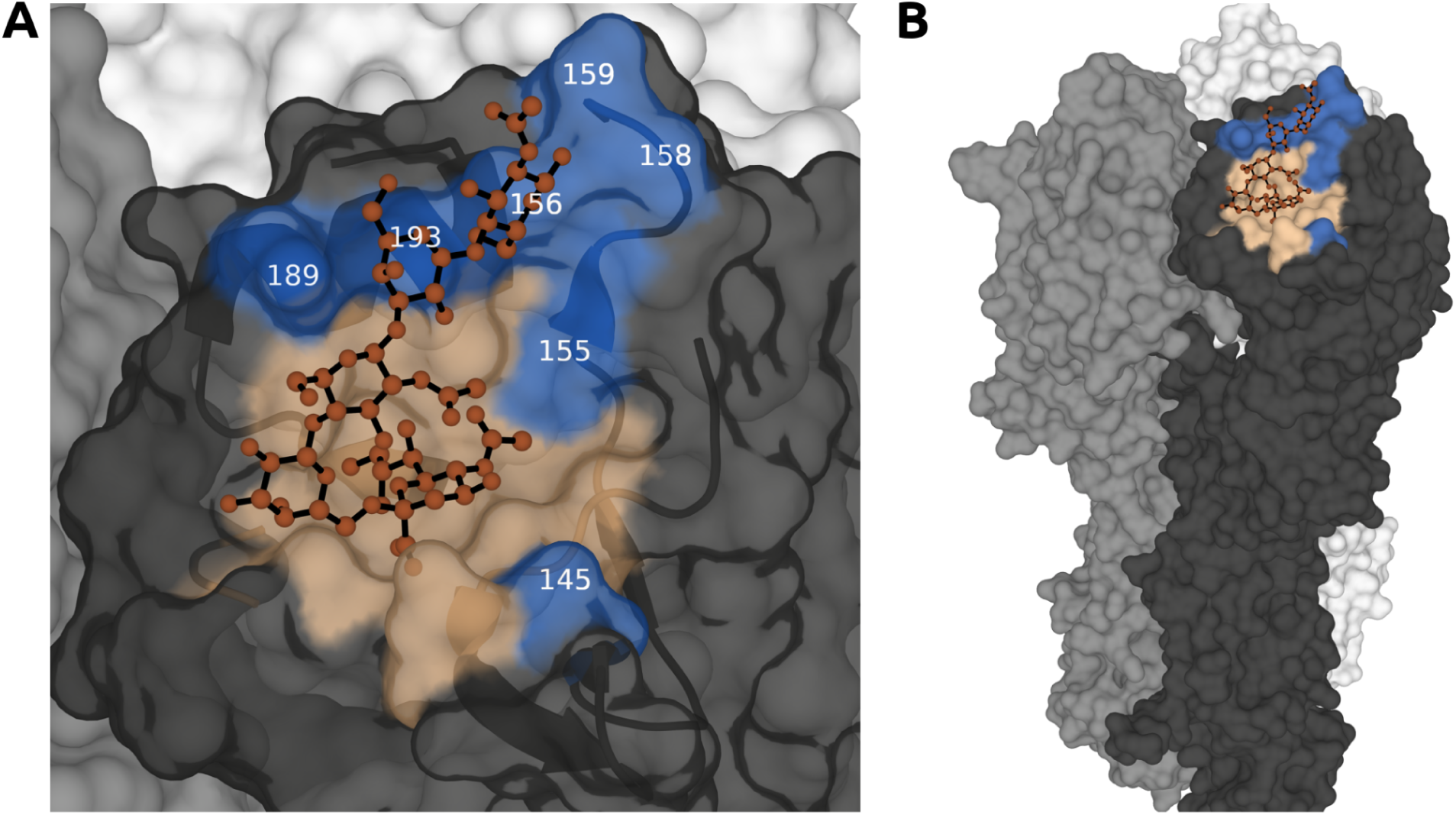
The Receptor Binding Site and Positions Mutated Presented on A/Hong Kong/1/1968 **(B)** The receptor binding site (wheat color) and Koel seven positions (marine) shown on HA structures (pdb id: 6TZB). Sticks show a human receptor analogue 6’-SLNLN bound to the receptor binding site. The secondary structure of the receptor binding site is shown beneath the transparent surface. **(B)** Zoomed out version of C, showing all three monomers.

Classical approaches to phenotypical characterization of viruses, albeit being very informative, scale up poorly in terms of time cost, both for the assay and the mutant generation component. Therefore they are more suitable for retrospective analysis rather than proactive surveillance. There have been several approaches to engineer high-throughput versions of these assays. One particular line of approach is to use Next Generation Sequencing (NGS) to replace the plaque counting / area estimation component of classical VN assays. Next Generation Sequencing (NGS) has seen widespread use in virology, including within the framework of Deep Mutational Scanning (DMS) (Li et al. 2016; Loes et al. 2024; Welsh et al. 2024; Dadonaite et al. 2024; Bloom and Neher 2023; Yu et al. 2024; Jian et al. 2024; Fowler and Fields 2014; Lei et al. 2023; Frank et al. 2022; Hom et al. 2019). In DMS experiments, viral populations are generated using random mutagenesis (RM), and these populations are then subjected to assays either in the absence of serum—to assess replicative fitness in vitro—or in the presence of serum to identify variants that escape neutralisation. A recent study (Welsh et al. 2024) utilized NGS in the context of DMS to construct analogues of virus neutralisation curves. Due to the presence of variants containing multiple substitutions, the study employed an epistatic model (with parameter regularization to prevent over-fitting due to existence of many interaction terms) to estimate the effect of individual mutations on an observable that functions as an NGS-based analogue of virus neutralisation titer. Building on this, other studies applied NGS to generate virus neutralisation curves outside the DMS context (Welsh et al. 2024; Loes et al. 2024). In these works, the authors individually constructed plasmids encoding viral variants known to circulate in nature, rescued them as a pooled mixture, and performed NGS-based neutralisation assays also as a mixture. Each viral variant was tagged with a unique barcode, enabling the reconstruction of individual neutralisation curves using classical curve fitting techniques (also with the help of an absolute standard to convert sequencing counts to proportions). The use of NGS in this way allows one to assay viruses as a mixture, rather than as individual variants, reducing the assay time requirements by orders of magnitude. RM on the other hand allows a very fast method to generate thousands of mutants in parallel. It, however, trades off scanning precision with depth and may rely on position-based mean inferences or epistatic models to decompose effects of multiple co-occurring substitutions into single ones–a non-trivial problem. It also does not guarantee generating every single mutant of interest due to its stochastic nature, although it can be optimized to favour single amino acid substitutions.

Selective mutational scanning (SMS) is a more controlled and precise alternative to DMS where the specific single (or sometimes multiple) amino-acid substitution mutants of interest are generated individually to ask more targeted questions about the evolution, as was done for instance in (Wilks et al. 2022; Mögling 2016). However even when just focusing on a restricted subset of positions such as the Koel seven, one potentially needs to generate hundreds of variants to be able to derive meaningful results about possible evolution routes before it occurs. Such directed mutagenesis approaches can become a crippling time bottleneck for pre-emptive surveillance. So, if the goal is a completely high-throughput SMS, a next generation sequencing (NGS) based high-throughput assay must be complemented with a similarly high-throughput but controlled method of producing a library of target mutants as a mixture, with as uniform proportions as possible for precision of results.

In this paper, we introduce the Next Generation Sequencing Replicative Fitness and Focus Reduction Assays (NGS RFA/FRA), which along with an optimized rescue and a statistical model for analysis of the results, establishes a high-throughput SMS pipeline that addresses these challenges (Fig.2) and extends upon the approach in (Mögling 2016). The variants in the assay are defined at the outset and assayed for replicative fitness and antigenic escape from selected sera. We demonstrate the utility of the assay by testing it on a mixture of 134 variants of A/Hong Kong/56/94 (A/HK/56/94, BE92 cluster) generated on the seven Koel positions and up to six monovalent ferret infection sera from nearby antigenic clusters. The bespoke statistical model allows rigorous comparisons of VN titers and replicative fitness at the level of individual variants. The procedure outlined here substantially reduces the amount of time required to explore a cluster’s future antigenic diversity based on a complete search of single amino acid substitutions in the Koel seven positions.

**Fig. 2.**
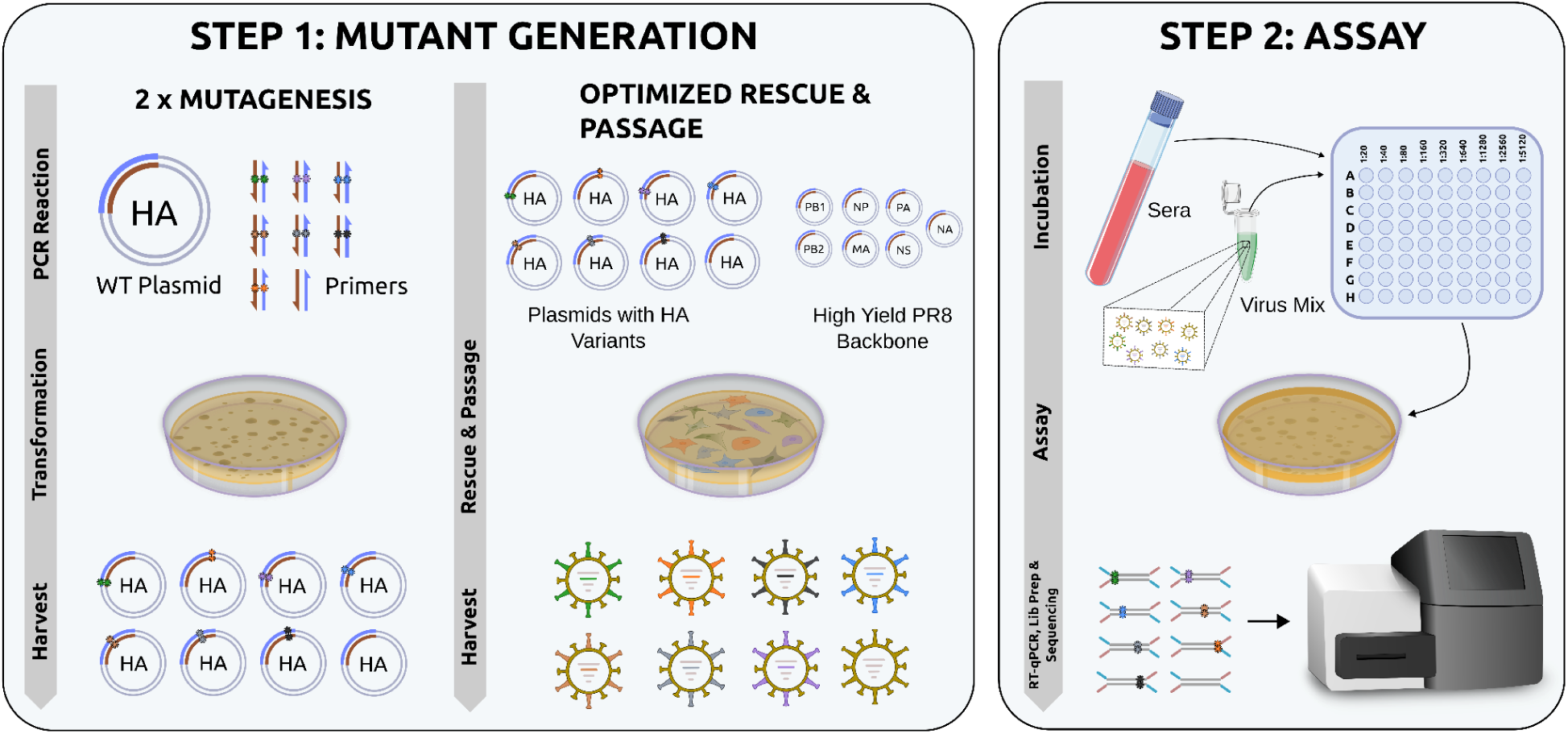
The NGS-FRA/RFA Experimental Pipeline. The panels outline the important steps of the experimental parts of the NGS-RFA/FRA pipeline in the context of this manuscript where we generate variants on Koel-seven positions. In Step 1, a mixture of variants is generated by an optimized mutagenesis, rescue and passage sub-pipeline. Mutagenesis part of Step 1 involves first generating wild-type plasmids with stop codons and then variant plasmids with stop codons by introducing substitution (this two step mutagenesis is therefore labelled as 2xmutagenesis). In Optimized Rescue & Passage, parameters such as Trypsin usage, incubation temperature and duration are optimized to yield a mixture of mutants with balanced proportions. Primers and plasmids with markers indicate variants of different positions and the one without any marker indicates the root virus A/Hong Kong/56/1994 (A/HK/56/94). The execution of step 2 is similar to classical VN assays except for two modifications: A mixture of viruses is used rather than a single variant and sequencing is used to quantify relative changes in proportions from non-neutralized to neutralized samples. Moreover RT-qPCR is used to quantify the total amount of viral genetic material in each sample. These data are then used to derive significant differences in fitness and titer (see <u>Fig. S1</u> for a schematic explaining the computational aspect). See “Experimental Method Details” and SI for details of each of the steps. The individual elements in this figure were prepared with Inkscape (Developers 2025) and have been deposited to open source repositories (“Bioicons - High Quality Science Illustrations,” n.d., “Open Science Art,” n.d.).

## RESULTS

The NGS-RFA/FRA SMS pipeline was designed with a specific quantitative target: to assay approximately 134 targeted single amino acid substitution variants in parallel, with sufficient virus per variant (approximately 50-100 PFUs/variant at 10,000 total PFUs) to achieve single-variant statistical resolution. The pipeline is divided into two main components: (1) generation of a balanced variant library, and (2) the NGS-based replicative fitness and focus reduction assays, including computational analysis and validation of results.

### **I.** Generation of a Balanced Variant Library

One of the technical difficulties in assaying 134 variants together is undersampling; a variant present at low initial frequency could lead to inflated noise in its measured phenotypes. Such an assay therefore benefits greatly from maintaining uniform proportions at each step of the variant generation stage — synthesis of the plasmid library, rescue on 293T cells, and passage on MDCK cells. This is one of the central technical challenges that motivates the optimisations of rescue and passage steps described here; since any systematic bias towards higher-fitness or root variants having higher proportions at earlier stages propagates into reduced precision for other variants at the assay stage.

### Producing Balanced HA Plasmid Libraries as a Mixture

The first step in the generation of a balanced virus library is the generation of an unbiased HA plasmid library. The experiments we describe here use the A/HongKong/56/1992 virus from the BE92 cluster as the root. As a template, we generated the unidirectional plasmid coding for hemagglutinin viral RNA of A/HK/56/94 under the control of the RNA polymerase I promoter (described previously in (Neumann et al. 1999)). We then sequenced the resulting libraries to verify variant proportions (see Experimental Method Details and SI Sections “Error Rate Analysis” and “Description of the in-house Code Packages”).

The optimized protocol employs two key design choices: pooling all primers for a given position into a single directed mutagenesis reaction, and using template plasmids bearing TAA stop codons at the target position rather than wild-type root plasmids (see Section “Mutagenesis to Create Plasmid Libraries of All Possible 20 Amino Acids at a Given Position” in” Experimental Method Details”). Pooled primers allow all amino acid variants at a position to be generated in a single reaction, substantially reducing the time required compared to individual variant generation. Stop-codon templates prevent overrepresentation of the wildtype root variant, arising from incomplete DpnI digestion of the template plasmid DNA, which was the dominant source of bias in earlier protocol variants. The proportions obtained with this earlier variant of the protocol where we did not use plasmid with stop codon revealed that apart from the root, the proportions of variants were quite uniform (Fig. 3A). This analysis also demonstrates that over-expression of the root plasmid carried over to the next stages of virus rescue in 293T cells and passage in MDCK cells lead to over-expression of root virus. The first two panels of (Fig. 3B) compare virus proportions after passage when respectively wild-type root plasmids and plasmids bearing stop-codons are used. The latter leads to a significant reduction in the proportion of the root variant as well as higher proportions for other variants.

**Fig. 3.**
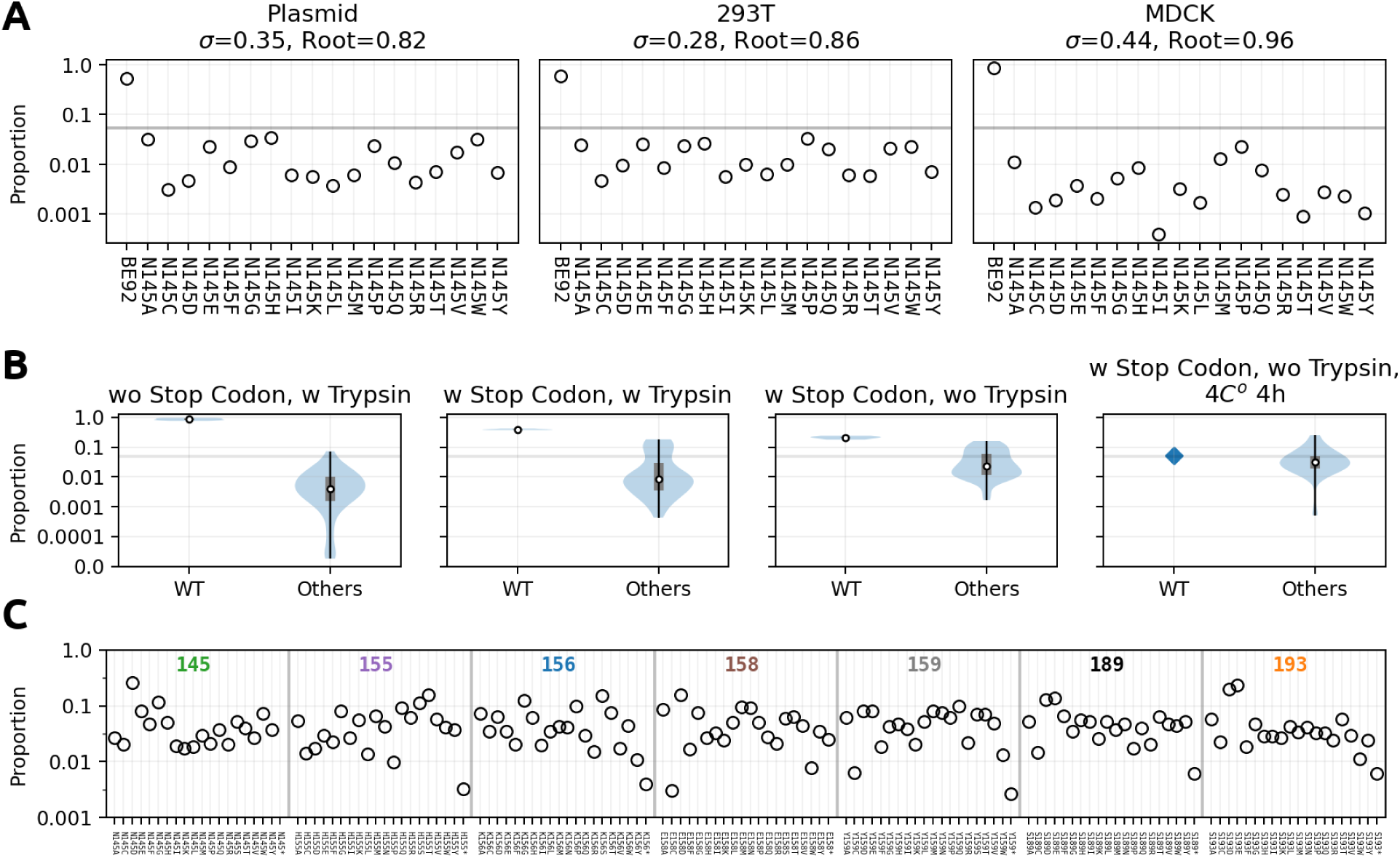
Several Stages of the Optimization for the Plasmid Generation, 293T Rescue and MDCK Passage. **(A)** Plasmid, post rescue (293T) and post passage (MDCK) proportions of variants of position 145 obtained with an earlier version of the rescue and passage protocol where “plasmids with stop codons” and “passage without Trypsin” optimizations are not used. The first subpanel is an average of six repeats and the following two are of twelve repeats. The title shows the standard deviation of the non root variants and the proportion of the root variant. The gray line shows the expected proportion; 1/20. N145S was excluded from the comparison because reverse primer was used for this variant whereas forward primer was used for all others. See panel C for an experiment where it was also included. **(B)** Each panel here shows the distribution of post MDCK passage proportions of variants of position 145, after further optimizations of the mutant generation step. The data in the first panel is identical to the one used in panel A. The second panel shows the case where templates with stop codon TAA were used instead of wild-type root plasmids (four repeats). Optimization in the third panel additionally carried out the final incubation of the 293T sup passage without Trypsin (four repeats). In the final panel, the time of attachment/inoculation was changed to 4°C for 4 hours (single repeat) and the root variant was generated separately from the rest of the mutants and combined based on PFU/ml measurements. For more details see Section “Virus Libraries Production” in “Experimental Method Details”. Within each panel, the data is separated into WT proportions and proportions of other variants. The white circle shows the median, the grey bar shows the quartile range and the vertical black line shows the full data range. The horizontal grey line shows the expected proportion; 1/20. **(C)** This shows the variant proportions within each of the seven Koel positions after the fully optimized rescue+passage (stop codons, no trypsin and 4°C for 4 hours) was utilized. The positions are grouped using vertical separators and indicated at the top of each group. The bars are centred around 1/20.

When generating libraries across multiple positions simultaneously — as required for the seven Koel positions — an additional source of root variant bias arises: if root primers are included in each position’s reaction, the combined mixture will contain the root variant at approximately seven-fold higher proportion than any individual variant. Consequently, the for the final version of the optimised protocol, we omit root primers from individual position reactions, and generate the root plasmid separately, combining it with position-specific libraries at the post-passage stage based on PFU/ml measurements (see Section “Virus Libraries Production” in “Experimental Method Details”). This is also implemented in anticipation of future applications possibly involving larger numbers of positions, where the root bias would become even more severe.

### Rescue and Optimized Passage of Virus Libraries

Once plasmid libraries are prepared, the seven position-specific mixtures are each rescued independently on 293T cells in the background of the attenuated vaccine high-yield strain A/Puerto Rico/8/1934 (PR/8 HY, PMID: 26334134), then passaged on MDCK cells, and finally combined into a mixture of 133 single-aa mutants plus the root virus. The optimised passage protocol uses two modifications designed to minimise fitness-dependent distortion of variant proportions (see Section “Virus libraries amplification in MDCK cells” in “Experimental Method Details”). First, the final incubation of 293T supernatant on MDCK cells is performed without trypsin, restricting infection to a single round of replication and thereby limiting the influence of replicative fitness differences on post-passage proportions (Fig. 3B, third panel). Second, this initial incubation is carried out at 4 degrees C for 4 hours rather than the more usual 37 degrees C for 2 hours, which increases the equilibrium fraction of virions bound to cells (Nunes-Correia et al. 1999) and thereby reduces the attachment disadvantage of low-fitness variants — including antigenically important escape mutants such as N145K — before the wash with PBS removes any unbound virus (Fig. 3B, fourth panel). All seven mixtures for the positions 145, 155, 156, 158, 159, 189, 193 rescued using the optimized protocol are shown in Fig. 3C.

### **II.** The NGS-based replicative fitness and focus reduction assay

Similar to the input mixture generation step, assaying 134 variants requires careful calibration of assay parameters to ensure unbiased access to resources for each virus. Once optimised in a no-serum context, the same conditions are then applied to VN experiments in which viruses are pre-incubated with sera before plating.

### The No-Serum Assay

In the no-serum assay, the virus mixture is grown on a plate with an Avicel overlay for 30 hours, and supernatants are collected in duplicate for RT-qPCR quantification and Illumina sequencing. CT values from RT-qPCR quantify total viral genomic matrix gene content, while hemagglutinin sequencing read proportions are compared to input read proportions to derive per-variant replicative fitnesses. This assay setup was adapted from the procedure outlined in (Mögling 2016) where Avicel was used to limit viral diffusion and obtain round plaques for radius plaque, (measured at T=30h). In the context of NGS-RFA/PFRA, the limitation on diffusion ensures that each variant replicates largely in parallel rather than competing with others for resources, consistent with the design goal of approximately uniform per-variant resource availability. The 30-hour growth duration was selected based on an initial experiment testing across different time points from 10-48h using an individually-rescued reference mixture of twenty variants (145N root plus 145K, 145R, 145Y, 155R, 156Q, 158G, 159S, 193W, 156E, 156F, 155Q, 155V, 145S, 155Y, 155T, 193F, 158D, 159N, 189K, which we call mix-20var), chosen by (Mögling 2016) to span a wide range of antigenic and fitness characteristics. Because these variants were rescued individually and combined based on plaque titration, this mixture is free of mixed-rescue variability and serves as a clean test of the assay step alone. The assay was performed using 1000 PFUs of the mixture and no serum. Similar to (Mögling 2016), we have observed that some viruses are completely missing from the supernatant at earlier time points (Fig. 4A, T=10h). Viruses display growth up to time point t=48h (Fig. 4B). Replicative fitness was estimated as the ratio of post-assay to input proportions at each time point (Fig. 4C). The correlation between NGS-based replicative fitness and plaque radii (Mögling 2016) stabilised after T=20h, with T=30h showing the best combination of complete variant recovery and consistent correlation with plaque radii for all variants including the 145Y and 156F outliers (Fig. 4C-D).

**Fig. 4.**
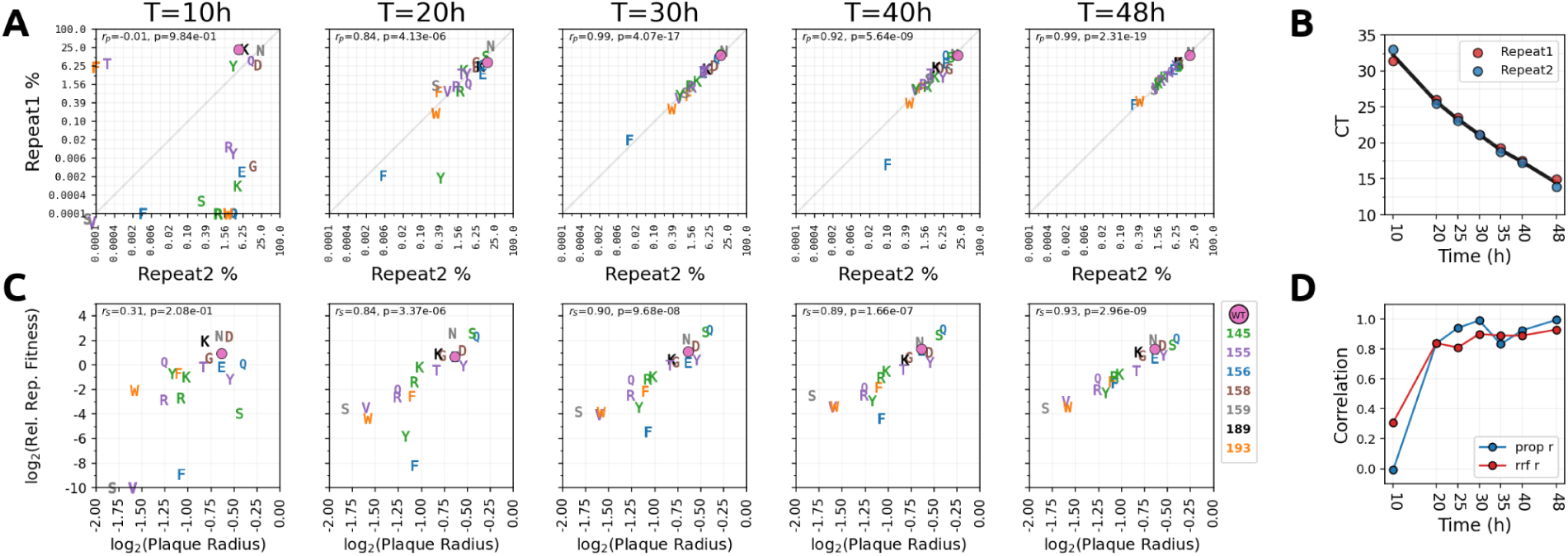
Testing Various Growth Times. **(A)** Shows comparison of variant % of repeats at various chosen time points (indicated on the titles). **(B)** Shows the CT values for the two repeats where the virus mix is grown on a plate with Avicel overlay up to the time point indicated at the x axis. **(C)** Shows comparison of log_10_ replicative fitnesses (y-axis, averaged over repeats) at vs plaque radii (x-axis). In panels A and C, colors indicate position as given in the legend and markers indicate variant amino acid. Round marker indicates the root variant. Variants which have zero proportion are plotted on top of the axis at which they have a non-zero value or at the bottom left corner of the plot if both proportions are zero. The inset text shows pearson correlation and p-value in the case of (A) and spearman correlation and p-value for (C). **(D)** Shows pearson (from A, in blue) and spearman correlations (from C, in red) across different time points.

### Measuring Antibody Escape: The FULL NGS-FRA/RFA

Antibody escape is measured by adding a serum incubation step before the virus replication assay. RDE-treated sera at two-fold serial dilutions from 1/20 to 1/2560 (or 1/5120 or 1/10240 for more reactive sera) are incubated with the input virus mixture for 1 hour at 37 °C before plating on MDCK cells with an Avicel overlay. Supernatant is collected at T=30 to measure qRT-PCR Ct values and sequencing counts. As in the no serum assay we tested multiple time points (T=30h, 40h) to verify that they gave comparable results (<u>Fig SI. 26</u>). Similar to a classical VN assay, the incubation at different dilutions are carried out so that per-variant VN titers can be calculated.

On the computational side, we developed a pipeline that estimates VN titers directly from the sequencing counts and qRT-PCR Ct values. The bulk of the model is built from a small set of parameters — a per-variant replicative fitness, per-serum/per-variant titers, and a per-serum slope — and generates the observed data through a chain of transformations. The titers and slopes first define a VN curve for each variant–serum combination; these curves, combined with the replicative fitnesses, are then converted into the expected proportion of each variant in the sequencing readout. The VN curves are therefore intermediate variables of the model and there is no observed remaining fraction of viruses that they are fitted to directly. Because the replicative fitness parameter enters the fit for every assay — with or without serum — it is estimated from the entire dataset, which increases their precision. The expected proportions thus constructed are compared against the observed sequencing counts using a per-variant BetaBinomial likelihood (batched over variants), whose concentration parameter models the overdispersion of the variant proportions. We compared a range of candidate likelihoods (SI); the BetaBinomial was selected because it best captures this overdispersion and therefore correctly reflects the uncertainty in low-proportion variants — an effect that is especially pronounced under strong neutralization, when many variants are driven to low abundance (<u>Fig</u> <u>SI. 2</u> and <u>Fig SI. 23</u>). Neither the VN curves nor the fraction of virus surviving neutralisation is directly observed, but both can be recovered from the fitted model. VN curves are sampled from the fitted titer and slope distributions, while the surviving fraction is "back-sampled" by inverting the final transformation above and applying it to the observed sequencing proportions. Full model details are given in SI Section “Description of the Model” (see <u>Fig SI. 1</u> for an overview). Construction of VN curves are detailed in the SI Section “Fitted Curve Examples”.

Fig. 5 A-E shows the results of the full pipeline tested on 134 variants (19 single substitution variants at each of the Koel seven positions 145, 155, 156, 158, 159, 189, 193 plus root variant) generated from the root virus A/HK/56/94 (we will refer to this mixture as mix1-ALL). We used two sera A/Beijing/32/1992 (BE92), A/Wuhan/359/1995 (WU95) first of which belongs to the same antigenic cluster as the root virus and the other one from the following future cluster. In generating mix1-ALL, we used the optimized rescue method which employs template plasmids with stop codons, the 4h at 4°C incubation of the 293T supernatant of rescued viruses,and the no-trypsin passage on MDCK cells (see section “Virus libraries amplification in MDCK cells” in “Experimental Methods Details”). Fig.5 C-E respectively show the predicted sequencing proportions, neutralisation curves and remaining fraction of variants obtained on this data.

**Fig. 5.**
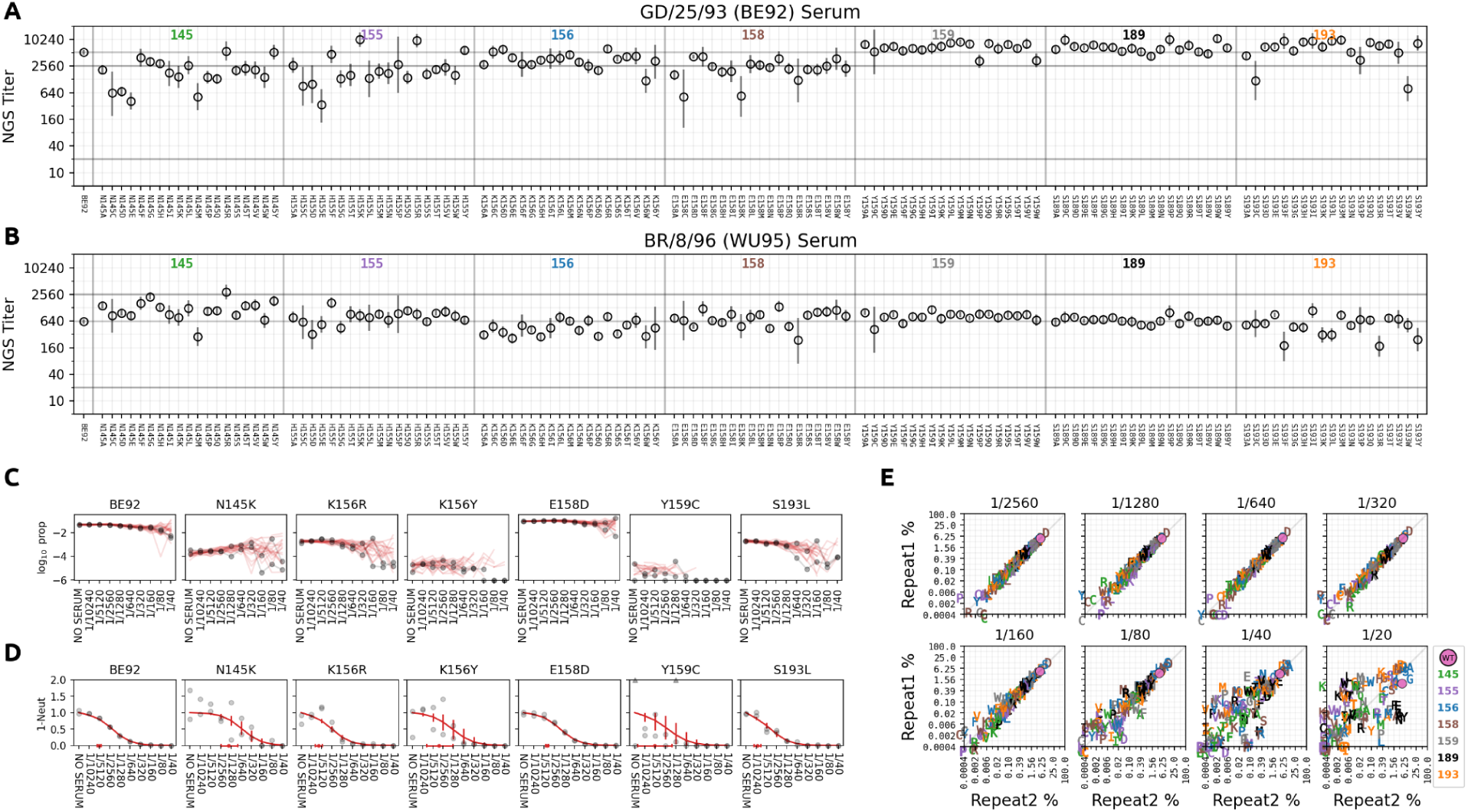
The NGS-FRA/RFA Full Pipeline Tested on Two Sera and 134 Variants. (**A-B)** y-axis shows titers obtained for the two sera indicated in the title using the NGS-PFRA/RFA pipeline. x-axis show the variants grouped according to position using vertical separators, with the colored numbers at the top indicating the position. The bars show the %95 High Density Intervals (HDI) bars. Heavy horizontal lines show experimental dilution limits and the light horizontal line shows the root variant titer level. The model titer range is +-1.5 log_2_ units of respectively upper limit and lower limit of detections for the experiment. **(C)** Comparison of observed repeat variant proportions (grey markers) vs twenty posterior predictions sampled from the Bayesian model (red lines) for six exemplary viruses with variable behaviour (in log_10_ scale). Zero observed proportions are plotted on the x-axis whereas zero sampled proportions are plotted as broken lines. x-axis shows dilutions on log_2_ scale, and NO SERUM at one minus the maximum dilution. **(D)** Comparison of latent samples for remaining fraction of six viruses (grey markers) to neutralisation curves (red) constructed from mean estimates for titer curve parameters. Markers outside the bound of the plot limits are shown with triangle markers (e.g. Y159C) at the top. x-axis is the same as the x-axis of panel C. Red markers and bars on the x-axis indicate mean estimate and %95 HDI for the titers. See SI section “Fitted Curve Examples” for details and more examples on how to construct neutralisation curves and latent remaining fractions. **(E)** Comparison of repeat variant %s across different dilutions (given in the title). Marker color indicates position as given by the legend and letter indicates variant. The round marker is the root variant.

Fig. 5E reveals the two main sources of noise in the system: 1-Highly reactive sera with low dilutions produce noisy patterns, most likely due to stochasticity associated with low number of PFUs left after neutralisation, 2-Variants that have low proportions (generally due to low replicative fitness or being more highly neutralized than others). The first type of noise affects all the variants en masse and usually only for lowest dilutions. The second type of noise can affect a variant throughout a longer dilution series (compare BE92 vs S193L in Fig. 5C). The computational pipeline encapsulates both of these noises by a suitable choice of likelihood and uses it to inform uncertainty of titers on a per variant basis.

### Repeatability and Cross-validation

We tested the assay using various repeatability and cross-validation tests by generating several other mixtures of differing complexity in their rescue methodology and population context and titrated them against six sera. To extend the number of sera, we focused on antigenic clusters A/Beijing/353/1989 (BE89), A/Beijing/32/1992 (BE92), A/Wuhan/359/1995 (WU95) and A/Sydney/5/1997 (SY97) near the root virus A/HK/56/94 (Smith et al. 2004) and first choose a subset of ten sera that can reflect the antigenic character of these clusters faithfully when one remakes the map subsetted to these clusters and sera (see section “Antigenic Cartography” in Supporting Information). From this subset of ten sera, we carry forward a further six sera (Fig. 6) chosen such that we have at least one serum from each of the clusters and that exhibit a wide range of titer values against the root variant.

**Fig. 6.**
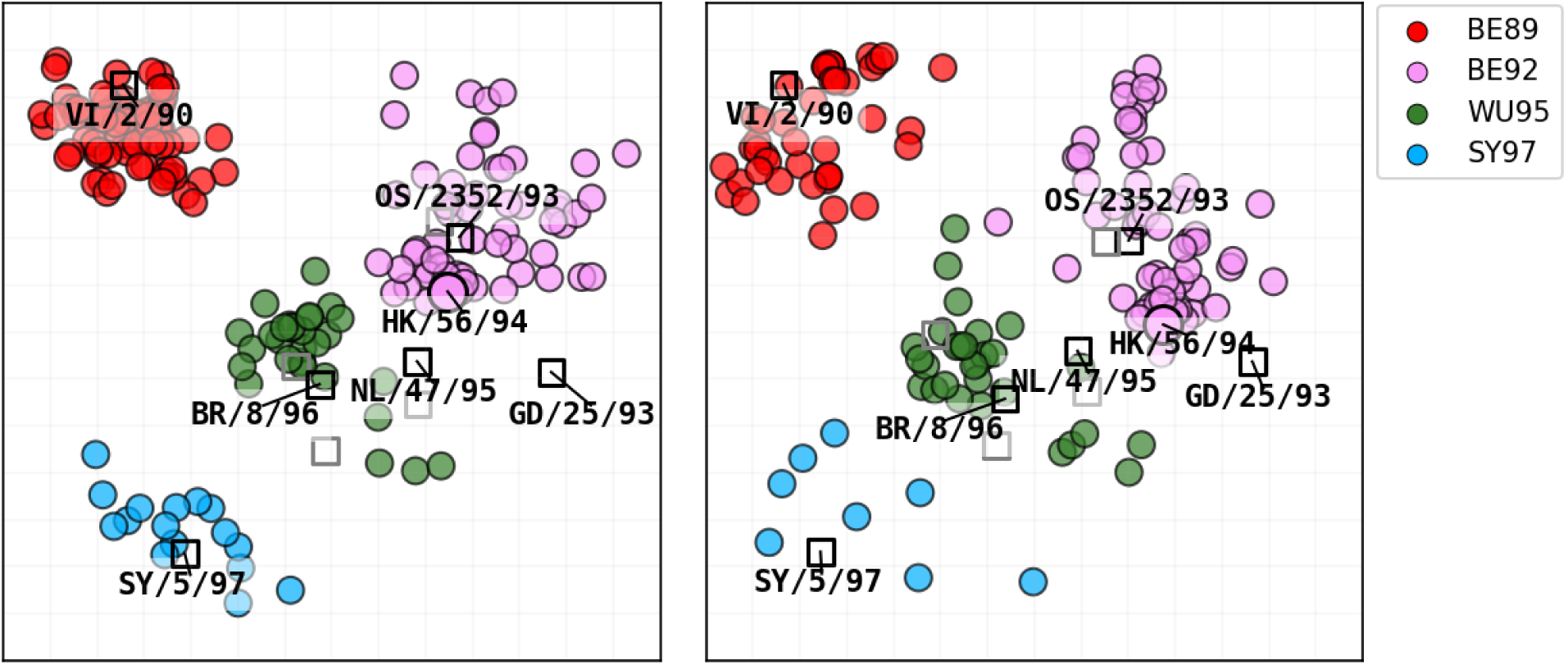
Selection of the Sera to Test and Positions to Mutate. **(A)** The antigenic map in (Smith et al. 2004) subsetted to clusters BE89, BE92, WU95 and SY97. Circles show antigens whereas squares show sera. The six sera tested in NGS-RFA/FRA validation are labelled and shown with darker squares and are labelled. The other lighter squares are the four sera that were selected as a part of the subsetting process. **(B)** Shows the map in (A) re-optimized with a subset of ten sera indicated by the squares. Legend indicates cluster of the antigens. Antigens for which there are less than three measured titrations (when subsetted as indicated) are not included in the optimization.

We generated mix2-ALL, consisting of the same variants and methodology as mix1-ALL, but without trypsin at the final incubation of the 293T sup and at 37C° for 2 hours instead of 4C° for 4 hours in the rescue. These two mixes provide a direct comparison between two independently rescued input mixtures in similar population contexts. mix1-ALL was assayed against two sera (GD/25/93, BR/8/96) and mix2-ALL was assayed against four sera (VI/2/90, GD/25/93, BR/8/96, SY/5/97) in duplicate for both. For each of mix1-ALL and mix2-ALL, we have also generated their counterparts only including the variants with substitutions at position 145, which we will refer to as mix1-posn145, mix2-posn145 (the former without root virus the latter with). These single position mixtures were used for testing repeatability of observables for the same variants in different population contexts. The smaller reference mixture containing twenty variants, mix-20var, was tested against all the six sera in quadruplicate. Because these viruses were rescued individually, mix-20var is free of mixed-rescue variability and serves as a control. The variants for the mix-20var were obtained from the same source as (Mögling 2016) and were also used for the comparator HI titrations. Therefore mix-20var will display the least amount of phenotypic variability when making comparisons to plaque radii or HI titers. Input PFUs were 1000 for mix1-posn145 and mix-20var, and 10,000 for mix1-ALL, mix2-posn145, and mix2-ALL; all assays used a 30-hour incubation time with Avicel overlay.

The NGS-RFA/FRA assay shows strong repeatability across a range of comparison types (Fig. 7). For replicative fitness, comparisons within the same rescue method but different variant contexts (mix1-posn145 vs mix1-ALL, Fig. 7A) and between different rescue methods in the same context (mix1-ALL vs mix2-ALL, Fig. 7B) both yielded orthogonal regression slopes above 0.88. Standard deviations were 0.12 (both x/y) for the same-rescue cross-context comparison and 0.40/0.46 (x/y) for the cross-rescue comparison, reflecting the additional variance expected when rescue methodology differs.

**Fig. 7.**
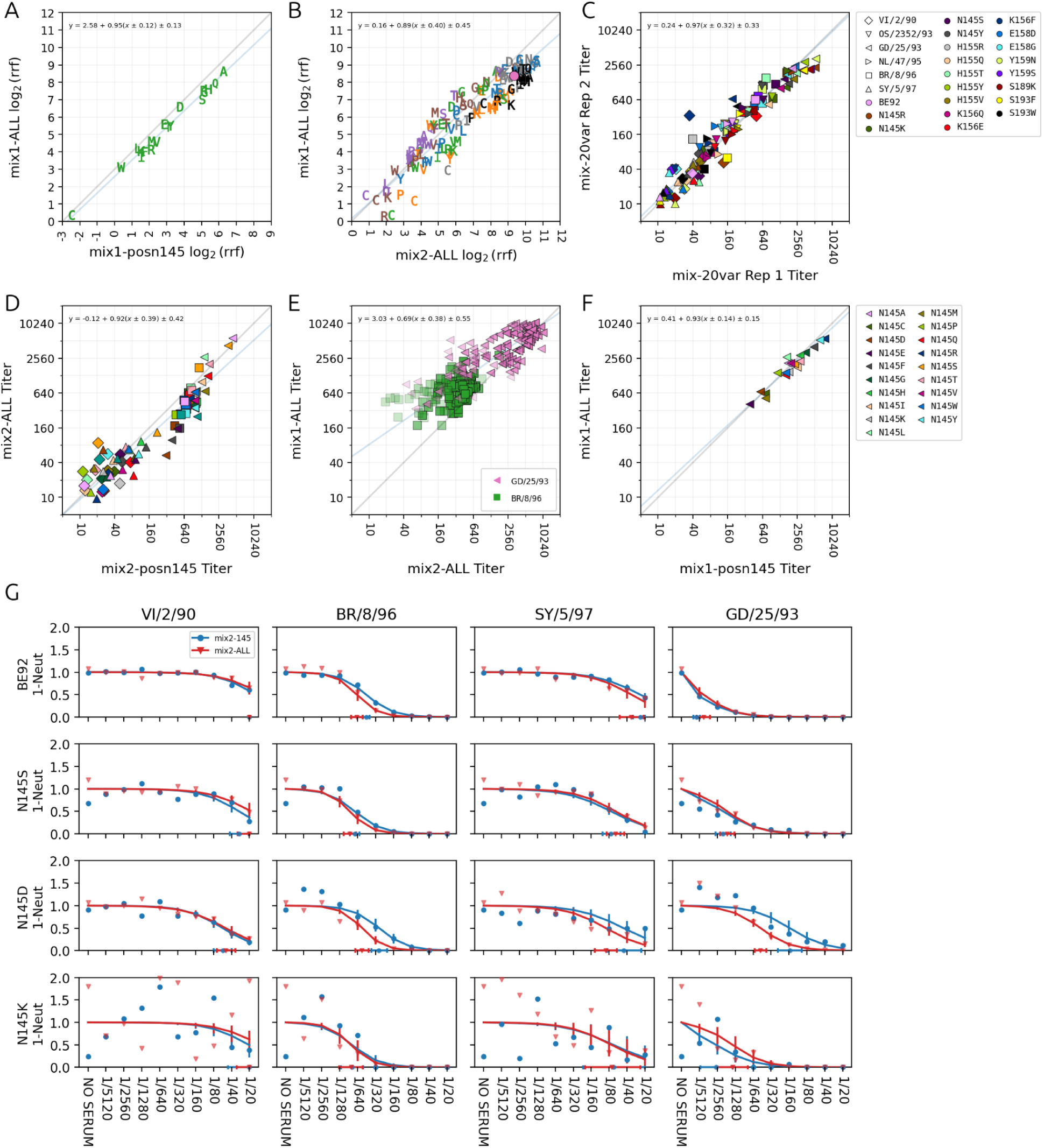
Reproducibility of NGS-RFA/FRA. **(A)** Comparison of relative replicative fitness for samples mix2-145 and mix2-ALL. **(B)** Comparison of relative replicative fitness for samples mix1-ALL and mix2-ALL. Letters indicate variant amino-acid whereas color indicates position with the same coloring as Fig. 5E **(C)** Comparison of titers obtained from training the model separately on two pairs of independent repeats for mix-20var. Markers indicate sera, color variant as given in the legend. **(D)** Comparison of titers (four sera) for samples mix1-ALL and mix1-145. Coloring and markers are the same as D. **(E)** Comparison of titers (two sera) for samples mix1-ALL and mix2-ALL. Markers indicate sera as given in the legend. The markers have transparency values that depend on their HDI values (maximum between mix1 and mix2) with those bigger than 3 in %95 HDI having alpha value of 0.25 and then linearly increasing to 1 as HDI decreases. **(F)** Comparison of titers (single serum GD/25/93) for samples mix2-ALL and mix2-145. Coloring is according to variant and is given in the legend. In E-F, the blue line is the orthogonal regression line and the grey line is the diagonal. **(G)** Exemplary neutralisation curves with increasing amounts of noise for four sera. Titer %95 HDI intervals for both cases are plotted on the x-axis with square markers.

Titer repeatability was equally strong. A technical repeat within mix-20var (Fig. 7C) had slope 1.00 with standard deviation 0.30 (both x/y). The same-rescue cross-context comparison (mix1-ALL vs mix1-pos145, Fig. 7D) had slope 0.91 with standard deviations 0.38/0.42. The cross-rescue comparison (mix1-ALL vs mix2-ALL, Fig. 7E) had slope 0.69 with standard deviations 0.39/0.50. There is a tendency for high HDI (i.e more noisy data) titers to have lower values (HDI is generally higher in the case of mix2-ALL i.e x-axis, not separately shown here). This is to be expected since in the limiting case where proportions are pure noise across dilutions would be best approximated by a straight line which would fit a low titer. Exemplary neutralisation curves for four variants and four sera with increasing levels of count repeat variability are shown in Fig. 7G, constructed as described in the SI section “Fitted Curve Examples”, using data from mix2-ALL and mix2-145.

Validation against classical assays also showed strong concordance across all mixture types. For replicative fitness compared to plaque radii, Spearman correlation was r=0.94 (p=6.34x10^-10^) in mix-20var where viruses from the same stock (Mögling 2016) were used in both comparisons, and was r=0.74 (p=1.68x10^-6^), and r=0.69 (p=1.45x10^-5^) respectively for mix1-ALL, and mix2-ALL where original stock was compared to viruses from the mixed rescue. Variant 159S’s replicative fitness did not agree with the plaque radii for both mix1-ALL and mix2-ALL but agreed with mix-20var suggesting a genuine biological rescue variability and not an assay error. An extended replicative fitness comparison using visually categorised plaque sizes across all 134 variants yielded Kendall tau_B_=0.49 (p=4.69x10^-14^, Fig. 8 A). The best fit regression slopes against HI titers were 0.89, 0.89, and 0.78 for the same three mixtures (Fig. 8D). The 155R variant was an outlier in the HI vs NGS-FRA titers for the GD/24/93 serum and was found to be an assay type difference (see <u>Fig SI. 23</u>C where 155R shows a similar difference when comparing FRA to HI).

**Fig. 8.**
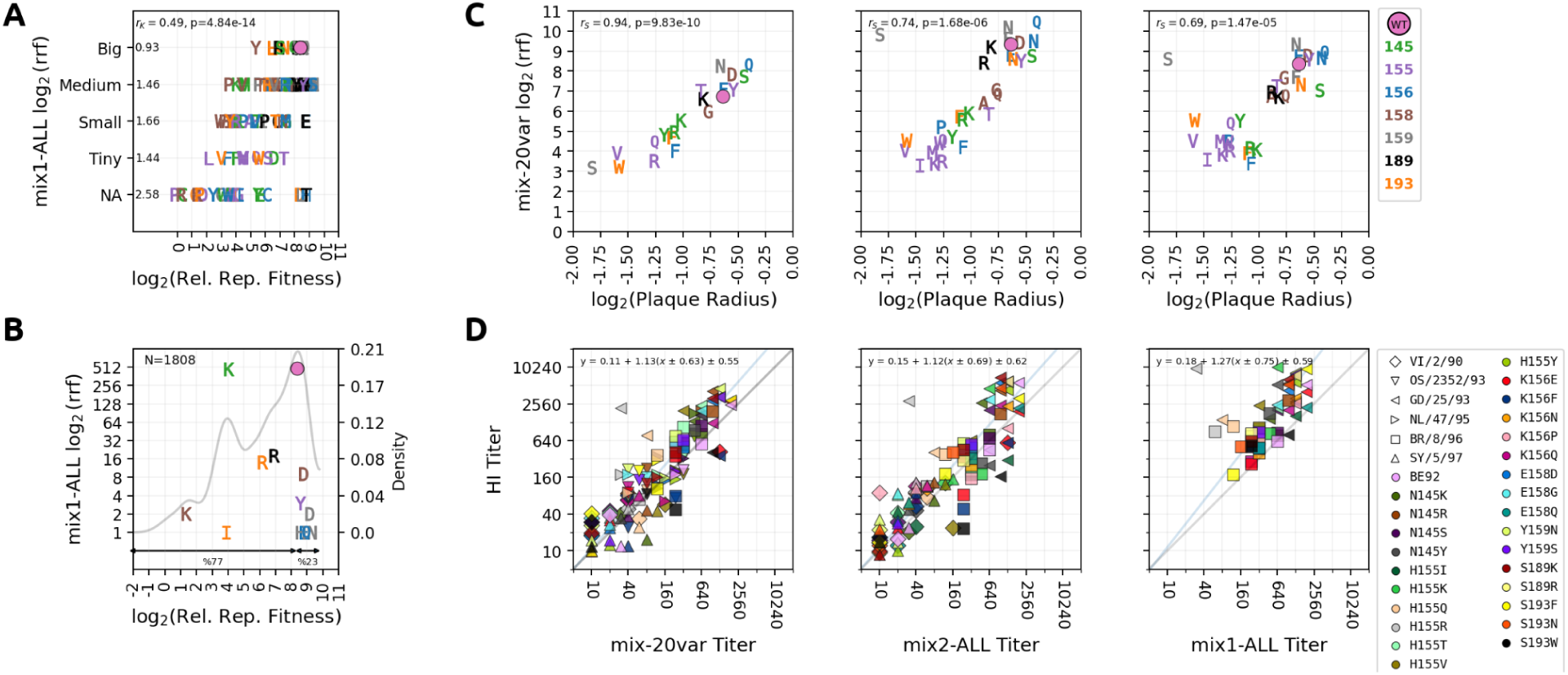
Validation of NGS-RFA/FRA Assay By Comparison to HI Assay and Plaque Radii. **(A)** Comparison of relative replicative fitness measured in mix2-ALL to plaque sizes of individually rescued variants as judged by eye [REF??, TIME POINT]. NA value on the y-axis indicates that the virus could not be individually rescued or no plaque visible. Inset value at the top shows Kendall τ_B_ correlation coefficient and its p-value. The other values along the y-axis show the standard deviation of the x-values within each group. **(B)** Comparison of relative replicative fitness measured in mix2-ALL to historical counts (within the time range 1979-1997) of these variants in GISAID (Elbe and Buckland-Merrett 2017). The grey line indicates the kernel density estimate of the log_2_(rrf) values with a bandwidth of 0.5 (with density values shown on the right y-axis). Ranges indicated beneath the double arrows at the bottom of the axis indicate the inverse percentiles from minimum to 8.3 and from 8.3 to maximum of the log_r_(rrf) values. **(C)** Comparison of plaque radii (Mögling 2016) to replicative fitness measured respectively in mix-20var, mix1 and mix2-ALL. Inset shows spearman correlation value and its p-value. Color indicates the position given in the legend and letter indicates variant. There are twenty variants being compared for mix-20var and 30 for mix1-ALL and mix2-ALL. **(D)** Comparison of HI titers to titers measured respectively in mix-20var (six sera given in the legend) mix1-ALL (four sera: VI/2/90, GD/25/93, BR/8/96, SY/5/97) and mix2-ALL (two sera: GD/25/93, BR/8/96). Marker shapes indicate the serum and color the variant as shown in the legend. Blue lines show the orthogonal regression line whose equation appears as inset.

To explore how NGS-RFA replicative fitness compares in vivo fitness, we looked at how often high replicative fitness viruses tend to appear in nature by computing the frequency of these variants in the GISAID database (Khare et al. 2021). We classified sequences within the time range 1979-1997 by their substitutions at the Koel seven positions relative to the root A/HK/56/94 (see “Counting GISAID Sequences” in “Supporting Information”) and single-substitution variant counts were compared to replicative fitness (Fig. 8 B). We make two observations: first, seven of the eleven variants (63%) with known natural occurrence (excluding the root and N145K) fall within the top 22% of the base replicative fitness distribution (indicated by the arrow spanning from rf=8.3 and to the end of the x-axis) of generated mutants. This demonstrates that naturally occurring viruses are biased towards the high-fitness tail and are not randomly sampled from this distribution. Since replicative fitness in MDCK cells is a proxy for one of the components (in situ replicative fitness) of overall virus fitness, these observations are consistent with the expectation that it should play a role in natural selection. Second, N145K is an expected outlier — despite relatively low replicative fitness it appears at high natural frequency, consistent with its role as the BE92-to-WU95 cluster-transition mutant whose antigenic escape compensates for its fitness cost.

To confirm that the NGS-RFA/FRA data captures known antigenic relationships, we constructed an antigenic map from mix-20var, which provides the most complete sera coverage: six sera including two from the BE92 cluster from which escape is being measured (Fig. 9). The relative positions of cluster sera in the map are consistent with the known antigenic topology from (Smith et al. 2004) (compare Fig. 6A). The behaviour of forward and reverse cluster transition mutants matches the observations in (Mögling 2016). Of all position 145 variants, N145K and N145R show the greatest movement towards the WU95 cluster, consistent with N145K being the known BE92-to-WU95 cluster transition substitution (Koel et al. 2013). N145K shows greater escape from the BE92 serum GD/25/93 compared to the root (Fig. 5A, Fig. 9A). Of all position 156 variants, K156E is the only one that moves in the direction of clusters SI87 and BE89, consistent with its role in the SI87-to-BE92 transition and with E being the residue at position 156 in both those clusters ((Smith et al. 2004)). The angular position of 156F superficially resembles 156E in the map, but its titer plot (Fig. 9A) reveals uniformly low titers against all sera, suggesting it is simply moving away from BE92 in any available direction, not necessarily towards BE89 — analogous to the low-titer position 155 variants described in (Mögling 2016). A comparison of replicative fitness against antigenic escape — measured as map distance for mix-20var and fold-change from BE92 for mix2-ALL (Fig. 9C) — reveals a pattern consistent with the analogous Figure 3B in (Mögling 2016). This pattern suggests that among well-adapted viruses, large antigenic escape from BE92 may be associated with a replicative fitness cost, while high fitness is compatible with a range of escape levels including near-zero.

**Fig. 9.**
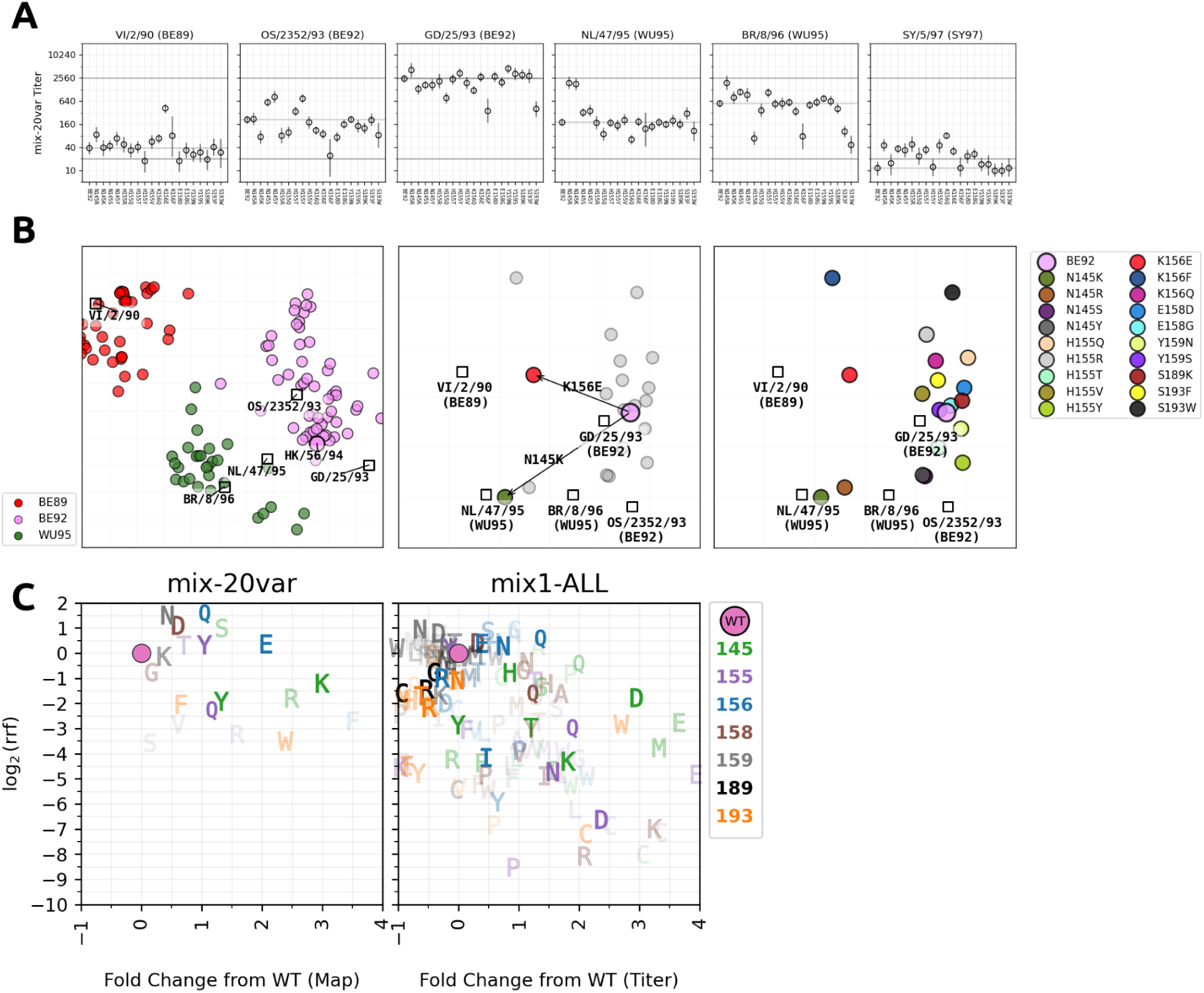
Checking the Validity of Antigenic Relations in NGS-RFA/FRA Data. **A)** Titer plots for the six sera titrated against mix-20var **(B)** The first panel shows the antigenic map from Fig.6B but zoomed on clusters BE89, BE92 and WU95. The second panel is an antigenic map made with data obtained from mix-20var. Arrows show important substitutions for forward (N145K) and reverse (K156E) cluster transitions. The root virus and the important transition mutants are colored according to the cluster colors in the first panel. The other variants are shown in grey. SY/5/97 serum is not used in the map due to its overall very low and non-discriminatory titers (see panel A and SI Section “Antigenic Cartography“ in "Supporting Information"). The map in the third panel is identical to the one in the second but variants are colored distinctly. **(C)** First panel shows antigenic distance (x-axis) vs relative replicative fitness (y-axis) obtained from mix-20var. The second panel is similar but the data is from mix2-ALL. Also instead of antigenic distance, fold-change of GD/25/92 titers from WT is used for the values at x-axis.

## DISCUSSION

Because of its mutation-driven antigenic evolution, influenza remains a global health challenge to this day, underlining the importance of tools for proactive surveillance efforts. This study presents a high-throughput SMS pipeline; NGS-RFA/FRA, which enables the rapid and efficient characterization of antigenic and replicative characteristics of influenza variants of interest, addressing a critical challenge in the field. The pipeline combines a mixture virus neutralisation assay with a targeted and controlled approach for generating mutants in uniform proportions.

We validated the pipeline with a retrospective analysis of the H3N2 cluster A/Beijing/32/1992 (BE92). It demonstrated excellent repeatability within the assay and strong agreement with classical HI titers and plaque radii. Although we used naive ferret sera against influenza viruses in this study, NGS-RFA/FRA can be naturally extended to human sera and to other viruses that can be effectively propagated in in vitro cell cultures which is currently under investigation.

Moreover, employing a Bayesian model customized for this experimental procedure enabled a relatively straightforward simulator construction for the system and allowed analysis of the relation between average number of PFUs per variant, neutralisation and the uncertainty of estimates (see SI section “Relation Between Noise, PFU and Average Neutralisation” for details). These simulations for instance quantified how increasing the number of variants impacted titer estimate uncertainty, leading us to conclude that a PFU/variant ratio corresponding to approximately 140 variants with 10,000 PFUs represents a reasonable stopping point in the context of this experiment (width of %95 HDI interval for pairwise differences peak at a difference of 2-fold, Fig SI. 22). It also allows one to study other effects such as the impact of reducing the number of repeats from two to one on the titer certainty.

The complementarity between DMS and SMS methodologies is note-worthy. The DMS approach can serve as a broad-brush technique to survey positions of antigenic interest agnostically, whereas SMS can act as a focused lens, delving into the regions of known antigenic impact with high resolution to tease apart the substitutions relevant to changing antigenic properties. By mapping potential immune escape routes for circulating strains, SMS can inform the design of more effective vaccines and therapeutic strategies, in a much shorter time span than classical methods of generating and assaying viruses individually.

### Limitations of the study

The SMS method of generating and assaying variants as presented here allows for less mutants than a typical DMS based study however it combines a more targeted mutational scanning with relatively uniform input proportions for each mutant of interest, leading to increased precision. DMS based mixture neutralisation studies can aggregate substitution escape scores into positions-wise estimates whereas our aim is to get as precise estimates as possible on a single variant level and therefore the need to engineer a more controlled scanning methodology.

An inherent requirement of this SMS pipeline is a prior selection of positions over which mutants can be generated. Whereas this might seem like a cyclic dependency, it is for instance already known that seven positions on the H3 HA majorly determine antigenic evolution from 1968 to 2003 (Koel et al. 2013) and doing an SMS experiment on seven positions is fully within the capacity of this pipeline even with all amino acids considered. More positions can be considered within a single experiment by removing historically unlikely substitutions. Moreover methods as presented in (Turner et al. 2026) or even a DMS based position scanning can be integrated into the pipeline to determine positions of interest. This method also does not provide any high-throughput means to generate a mixture of targeted double or triple mutants in balanced proportions. If such mutants are required, one would either have to rescue them individually, accept the extra mutants that arise as a side product of having primers in the mixture which target different positions or generate variants on a subset of positions using multiple roots.

As with every neutralisation type assay, this assay measures neutralisation in vitro using a particular cell-type and therefore is subject to identical caveats of comparing in vitro fitness to in vivo fitness. Although we have focused on the HA protein in this paper, in principle mutations on other proteins that might impact replication in ways other than binding can also be measured with this type of assay.

Time-wise, there are two main bottlenecks in this pipeline, one is the requirement of two mutagenesis experiments for generating the variants of interest and the second one is propagating viruses in cells. For the former, the first mutagenesis is the generation of template plasmids containing stop codons. This is done to prevent over-expression of the root variant due to possible incomplete DpnI digestion. In principle, our tests revealed that removing the root variant primers from the second mutagenesis step can be sufficient to remedy this bias without the need to generate plasmids with stop codons. For the latter, one could in principle incubate the viruses in suspension in the 96 well plates, for about the time it takes for a single round of replication, as is done in some of the DMS studies. This might however impact the replicative fitness readouts as the washing step involved in plate based assays likely affects the virus proportion readouts in a way correlated to fitness by removing from the system the viruses which can not bind as strongly. Moreover, less cycles of replication might increase subsampling error for lower replicative fitness viruses during supernatant collection. Experiments comparing both methodologies would be required to see each’s effect on replicative fitness/titers and how well both correlate with what is observed in nature.

## ACKNOWLEDGEMENTS

The authors would like to thank Mathis Funk, Willemijn Rijnink, Dennis de Meulder, Mark Pronk, Adinda Kok from EMC and Sam Wilks from University of Melbourne for useful discussions and their help with the experiments.

## AUTHOR CONTRIBUTIONS

**Conceptualization:** Si.T., S.J, Sa.T., A.N., A.M.H, T.C.J., D.J.S., M.R., T.M.B., R.F., G.N., Y.K.

**Methodology:** Si.T., S.J., D.J.S., M.R., T.M.B., R.S., R.F., G.N., Y.K

**Investigation:** Si.T., S.J., D.J.S., T.M.B., M.R., R.S., R.F., S.F., R.D., G.N., Y.K.

**Validation:** Si.T., D.J.S., M.R., T.M.B, R.S., R.F.

**Software:** Si.T.

**Formal Analysis:** Si.T.

**Visualization:** Si.T.

**Writing-original draft:** Si.T., D.J.S

**Writing-review&editing:** Si.T., D.J.S, M.R, G.N., Y.K. R.F.

**Resources:**M.R R.F. D.J.S.

**Funding Acquisition:**M.R R.F. D.J.S

## REFERENCES

1. Ayer, Miriam, H. D. Brunk, G. M. Ewing, W. T. Reid, and Edward Silverman. 1955. “An Empirical Distribution Function for Sampling with Incomplete Information.” The Annals of Mathematical Statistics 26 (4): 641–647. “Bioicons - High Quality Science Illustrations.” n.d. Accessed July 14, 2025. https://bioicons.com/.

2. Bloom, Jesse D., and Richard A. Neher. 2023. “Fitness Effects of Mutations to SARS-CoV-2 Proteins.” Virus Evolution 9 (2): vead055.

3. Boni, Maciej F. 2008. “Vaccination and Antigenic Drift in Influenza.” Vaccine 26 Suppl 3 (Suppl 3): C8–14.

4. Czado, Claudia, Tilmann Gneiting, and Leonhard Held. 2009. “Predictive Model Assessment for Count Data.” Biometrics 65 (4): 1254–1261.

5. Dadonaite, Bernadeta, Jenny J. Ahn, Jordan T. Ort, et al. 2024. “Deep Mutational Scanning of H5 Hemagglutinin to Inform Influenza Virus Surveillance.” PLoS Biology 22 (11): e3002916.

6. Developers, Inkscape Website. 2025. “Inkscape - Draw Freely.” May 12. https://inkscape.org.

7. Edgar, Robert C., and Henrik Flyvbjerg. 2015. “Error Filtering, Pair Assembly and Error Correction for next-Generation Sequencing Reads.” Bioinformatics (Oxford, England) 31 (21): 3476–3482.

8. Elbe, Stefan, and Gemma Buckland-Merrett. 2017. “Data, Disease and Diplomacy: GISAID’s Innovative Contribution to Global Health.” *Global Challenges (Hoboken*, NJ) 1 (1): 33–46.

9. Fowler, D. M., and S. Fields. 2014. “Deep Mutational Scanning: A New Style of Protein Science.” Nature Methods 11 (8). 10.1038/nmeth.3027.

10. Frank, Filipp, Meredith M. Keen, Anuradha Rao, et al. 2022. “Deep Mutational Scanning Identifies SARS-CoV-2 Nucleocapsid Escape Mutations of Currently Available Rapid Antigen Tests.” Cell 185 (19): 3603.

11. GitHub. n.d. “Prior Choice Recommendations.” Accessed July 11, 2025. https://github.com/stan-dev/stan/wiki/Prior-Choice-Recommendations.

12. Han, Alvin X., Simon P. J. de Jong, and Colin A. Russell. 2023. “Co-Evolution of Immunity and Seasonal Influenza Viruses.” Nature Reviews. Microbiology 21 (12): 805–817.

13. Hay, James A., Huachen Zhu, Chao Qiang Jiang, et al. 2024. “Reconstructed Influenza A/H3N2 Infection Histories Reveal Variation in Incidence and Antibody Dynamics over the Life Course.” PLoS Biology 22 (11): e3002864.

14. Hom, Nancy, Lauren Gentles, Jesse D. Bloom, and Kelly K. Lee. 2019. “Deep Mutational Scan of the Highly Conserved Influenza A Virus M1 Matrix Protein Reveals Substantial Intrinsic Mutational Tolerance.” Journal of Virology 93 (13). 10.1128/JVI.00161-19.

15. Iuliano, A. Danielle, Katherine M. Roguski, Howard H. Chang, et al. 2018. “Estimates of Global Seasonal Influenza-Associated Respiratory Mortality: A Modelling Study.” *Lancet (London*, England) 391 (10127): 1285–1300.

16. Jian, Fanchong, Jing Wang, Ayijiang Yisimayi, et al. 2024. “Evolving Antibody Response to SARS-CoV-2 Antigenic Shift from XBB to JN.1.” Nature 637 (8047): 921.

17. Kärber, G. 1931. “Beitrag zur kollektiven Behandlung pharmakologischer Reihenversuche.” Naunyn-Schmiedeberg’s archives of pharmacology 162 (4): 480–483.

18. Katoh, Kazutaka, John Rozewicki, and Kazunori D. Yamada. 2019. “MAFFT Online Service: Multiple Sequence Alignment, Interactive Sequence Choice and Visualization.” Briefings in Bioinformatics 20 (4): 1160–1166.

19. Khare, Shruti, Céline Gurry, Lucas Freitas, et al. 2021. “GISAID’s Role in Pandemic Response.” China CDC Weekly 3 (49): 1049–1051.

20. Koel, Björn F., David F. Burke, Theo M. Bestebroer, et al. 2013. “Substitutions near the Receptor Binding Site Determine Major Antigenic Change during Influenza Virus Evolution.” *Science (New York*, N.Y.) 342 (6161): 976–979.

21. Krammer, Florian, Gavin J. D. Smith, Ron A. M. Fouchier, et al. 2018. “Influenza.” Nature Reviews. Disease Primers 4 (1): 3.

22. Kumar, Ravin, Colin Carroll, Ari Hartikainen, and Osvaldo Martin. 2019. “ArviZ a Unified Library for Exploratory Analysis of Bayesian Models in Python.” Journal of Open Source Software 4 (33): 1143.

23. Lei, Ruipeng, Andrea Hernandez Garcia, Timothy J. C. Tan, et al. 2023. “Mutational Fitness Landscape of Human Influenza H3N2 Neuraminidase.” Cell Reports 42 (1): 111951.

24. Li, Chengjun, Masato Hatta, David F. Burke, et al. 2016. “Selection of Antigenically Advanced Variants of Seasonal Influenza Viruses.” Nature Microbiology 1 (6): 16058.

25. Loes, Andrea N., Rosario Araceli L. Tarabi, John Huddleston, et al. 2024. “High-Throughput Sequencing-Based Neutralization Assay Reveals How Repeated Vaccinations Impact Titers to Recent Human H1N1 Influenza Strains.” Journal of Virology 98 (10): e0068924.

26. Marco-Sola, Santiago, Jordan M. Eizenga, Andrea Guarracino, Benedict Paten, Erik Garrison, and Miquel Moreto. 2023. “Optimal Gap-Affine Alignment in O(s) Space.” *Bioinformatics (Oxford*, England) 39 (2). 10.1093/bioinformatics/btad074.

27. Marco-Sola, Santiago, Juan Carlos Moure, Miquel Moreto, and Antonio Espinosa. 2021. “Fast Gap-Affine Pairwise Alignment Using the Wavefront Algorithm.” *Bioinformatics (Oxford*, England) 37 (4): 456–463.

28. Miller, Rupert G. 1973. “Nonparametric Estimators of the Mean Tolerance in Bioassay.” Biometrika 60 (3): 535.

29. Mögling, Ramona. 2016. “Evolution of Human Seasonal Influenza Viruses : Intrinsic and Extrinsic Fitness / Ramona Mögling.” 2016.

30. Neumann, G., T. Watanabe, H. Ito, et al. 1999. “Generation of Influenza A Viruses Entirely from Cloned cDNAs.” Proceedings of the National Academy of Sciences of the United States of America 96 (16): 9345–9350.

31. Nguyen, Alan B. H., Marco Bonici, Glen McGee, and Will J. Percival. 2025. “LOO-PIT: A Sensitive Posterior Test.” Journal of Cosmology and Astroparticle Physics 2025 (01): 008.

32. Nunes-Correia, I., J. Ramalho-Santos, S. Nir, and M. C. Pedroso de Lima. 1999. “Interactions of Influenza Virus with Cultured Cells: Detailed Kinetic Modeling of Binding and Endocytosis.” Biochemistry 38 (3): 1095–1101. “Open Science Art.” n.d. Accessed July 14, 2025. https://openscienceart.com/.

33. Pauly, Matthew D., Megan C. Procario, and Adam S. Lauring. 2017. A Novel Twelve Class Fluctuation Test Reveals Higher than Expected Mutation Rates for Influenza A Viruses. June 9. 10.7554/eLife.26437.

34. “Racmacs.” n.d. Accessed December 1, 2025. https://acorg.github.io/Racmacs/index.html.

35. Smith, Derek J., Alan S. Lapedes, Jan C. de Jong, et al. 2004. “Mapping the Antigenic and Genetic Evolution of Influenza Virus.” Science 305 (5682): 371–376.

36. Šošić, Martin, and Mile Šikić. 2017. “Edlib: A C/C ++ Library for Fast, Exact Sequence Alignment Using Edit Distance.” Bioinformatics 33 (9): 1394–1395.

37. Spearman, C. 1908. “The Method of Right and Wrong Cases (Constant Stimuli) without Gauss’s Formulae.” British Journal of Psychology 2 (3): 227.

38. Spearman, C. 1909. “Review of the Method of ’right and Wrong Cases’ (’constant Stimuli’) without Gauss’s Formula.” Psychological Bulletin 6 (1): 27–28.

39. Tenforde, Mark W., Rebecca J. Garten Kondor, Jessie R. Chung, et al. 2021. “Effect of Antigenic Drift on Influenza Vaccine Effectiveness in the United States-2019-2020.” Clinical Infectious Diseases : An Official Publication of the Infectious Diseases Society of America 73 (11): e4244–e4250.

40. Tureli, Sina. 2025a. “Merlign.” GitHub. https://github.com/iAvicenna/Merlign. Tureli, Sina. 2025b. “Mitril.” GitHub. https://github.com/iAvicenna/Mitril.

41. Tureli, Sina. 2025c. “NGS-RFA/FRA Paper Repository.” GitHub. https://github.com/iAvicenna/ngs_rfa_fra_paper_repo.

42. Tureli, Sina. 2025d. “Thor.” GitHub. https://github.com/iAvicenna/Thor.

43. Turner, Samuel A., David J. Pattinson, Ron A. M. Fouchier, and Derek J. Smith. 2026. “Predicting the Antigenic Evolution of Seasonal Influenza Viruses Using Phylogenetic Convergence.” In bioRxiv. April 10. 10.64898/2026.04.10.717627.

44. Vehtari, Aki, Andrew Gelman, and Jonah Gabry. 2015. Practical Bayesian Model Evaluation Using Leave-One-out Cross-Validation and WAIC. July 16. 10.1007/s11222-016-9696-4.

45. Vehtari, Aki, Andrew Gelman, Daniel Simpson, Bob Carpenter, and Paul-Christian Bürkner. 2021. “Rank-Normalization, Folding, and Localization: An Improved R^ for Assessing Convergence of MCMC (with Discussion).” Bayesian Analysis 16 (2). 10.1214/20-ba1221.

46. Welsh, Frances C., Rachel T. Eguia, Juhye M. Lee, et al. 2024. “Age-Dependent Heterogeneity in the Antigenic Effects of Mutations to Influenza Hemagglutinin.” Cell Host & Microbe 32 (8): 1397–1411.e11.

47. Wilks, Samuel H., Barbara Mühlemann, Xiaoying Shen, et al. 2022. “Mapping SARS-CoV-2 Antigenic Relationships and Serological Responses.” In bioRxiv. January 28. 10.1101/2022.01.28.477987.

48. Yu, Yingpu, Maximilian A. Kass, Mengyin Zhang, et al. 2024. “Deep Mutational Scanning of Hepatitis B Virus Reveals a Mechanism for Cis-Preferential Reverse Transcription.” Cell 187 (11): 2735–2745.e12.

