## Supporting Information for "NGS-RFA/FRA: A High Throughput Experimental and Computational Pipeline for Selective Mutational Scanning in Parallel"

**Supporting Information for the Article “NGS-RFA/FRA: A High Throughput Experimental and Computational Pipeline for Selective Mutational Scanning in Parallel”**

Sina Tureli*[1], Theo Bestebroer*[2], Sarah James[1], Rachel Scheuer[2], Randall Dahn[3], Shufang Fan[3], Sam Turner[1], Sam Wilks[1,‡], Antonia Netzl[1], A. Mosterin Hopping[1], Terry C. Jones[1,4], Gabriele Neumann[3], Yoshihiro Kawaoka[3], Ron Fouchier[2], Mathilde Richard[2]**, Derek J. Smith[1]**

**Affilitiations:**
[1] Center for Pathogen Evolution, Department of Zoology, University of Cambridge; Cambridge, CB25 3EJ, UK.
[2] Department of Viroscience, Erasmus Medical Centre, Rotterdam 3015 CE, the Netherlands
[3] Department of Pathobiological Sciences, Influenza Research Institute, University of Wisconsin-Madison, Madison, Wisconsin, USA
[4] Institute of Virology, Charité - Universitätsmedizin Berlin, Corporate Member of Freie Universität Berlin, Humboldt-Universität zu Berlin and Berlin Institute of Health, Berlin, Germany
[‡]Present address: Institute for Electrical and Electronic Engineering, Faculty of Engineering and Information Technology, University of Melbourne; Melbourne, Australia

***:** Co-first authors
**:Co-corresponding authors

Co-corresponding author emails:
Mathilde Richard:
Derek J. Smith:

### SUPPLEMENTAL INFORMATION

The supplementary information contains details on the model construction (SI Section 1), model convergence, goodness of fit and cross-validation diagnostics (SI Section 2), extra analysis not presented in the main manuscript (SI Section 3) and description of the in-house packages used for the analysis (SI Section 4).

### 1. DESCRIPTION OF THE STATISTICAL MODEL

#### 1.1 The Data Structure Overview

A typical data table after processing the reads looks like

**Table SI. 1 A Representative Data Table**

The first three columns give information about the sample, the fourth column is the CT value measured after the assay is complete and the remaining columns give counts of variants obtained from analysing the sequences. Every row is used as an observed data in the likelihood. The dilutions values appear in the model as covariates after appropriate log transformations.

| SERUM | REPEAT | DILUTION | CT | BE92 | N145A | N145C | N145D | N145E | N145F | N145G | N145H | N145I | N145K | N145L | N145M | N145P | N145Q | N145R | N145S | N145T | N145V | N145W | N145Y |
|---|---|---|---|---|---|---|---|---|---|---|---|---|---|---|---|---|---|---|---|---|---|---|---|
| **F09016** | **A** | **1/20** | 20 | 17895 | 224 | 0 | 1622 | 58 | 4 | 148 | 733 | 1 | 1 | 8 | 1334 | 64 | 110 | 0 | 93 | 17 | 3 | 59 | 7 |
| **F09016** | **A** | **1/40** | 18 | 20068 | 1193 | 1 | 6109 | 447 | 41 | 197 | 117 | 6 | 4 | 1 | 1082 | 58 | 368 | 2 | 195 | 19 | 6 | 211 | 13 |
| **F09016** | **A** | **1/80** | 17 | 16985 | 1630 | 1 | 3460 | 342 | 92 | 508 | 706 | 74 | 6 | 68 | 509 | 707 | 1467 | 6 | 1088 | 64 | 127 | 28 | 21 |
| **F09016** | **A** | **1/160** | 14 | 22285 | 513 | 5 | 4065 | 790 | 77 | 927 | 1320 | 121 | 79 | 39 | 390 | 649 | 1515 | 7 | 2085 | 127 | 29 | 131 | 78 |
| **F09016** | **A** | **1/320** | 13 | 26130 | 1394 | 1 | 5876 | 1005 | 126 | 1506 | 1659 | 55 | 49 | 87 | 410 | 1208 | 1448 | 40 | 1834 | 327 | 145 | 225 | 163 |
| **F09016** | **A** | **1/640** | 13 | 23509 | 2050 | 6 | 5232 | 987 | 80 | 2587 | 2640 | 28 | 96 | 50 | 172 | 1003 | 1777 | 35 | 2066 | 348 | 112 | 181 | 283 |
| **F09016** | **A** | **1/1280** | 13 | 19268 | 2990 | 5 | 6549 | 933 | 291 | 3418 | 2548 | 74 | 92 | 46 | 161 | 673 | 1823 | 16 | 2103 | 311 | 179 | 202 | 348 |
| **F09016** | **A** | **1/2560** | 13 | 20284 | 2604 | 20 | 7212 | 1147 | 170 | 5222 | 3222 | 113 | 113 | 49 | 167 | 1215 | 2700 | 122 | 2461 | 462 | 191 | 287 | 468 |
| **F09016** | **A** | **1/5120** | 13 | 36045 | 7509 | 6 | 10470 | 1600 | 377 | 11401 | 5431 | 187 | 68 | 278 | 303 | 3736 | 5420 | 168 | 4753 | 920 | 447 | 267 | 762 |
| **NO SERUM** | **A** | | 13 | 18073 | 2681 | 2 | 4268 | 600 | 161 | 4691 | 2524 | 64 | 59 | 133 | 246 | 1125 | 1903 | 184 | 1943 | 312 | 115 | 137 | 344 |
| **F09016** | **B** | **1/20** | 21 | 3975 | 148 | 0 | 639 | 95 | 3 | 181 | 94 | 2 | 2700 | 3 | 23 | 78 | 787 | 0 | 102 | 50 | 9 | 2933 | 12 |
| **F09016** | **B** | **1/40** | 18 | 17820 | 125 | 1 | 1745 | 1982 | 7 | 125 | 127 | 0 | 12 | 1 | 1275 | 55 | 140 | 1 | 83 | 26 | 7 | 41 | 14 |
| **F09016** | **B** | **1/80** | 16 | 17814 | 112 | 1 | 2190 | 338 | 51 | 398 | 835 | 4 | 36 | 33 | 355 | 949 | 2246 | 22 | 827 | 25 | 7 | 113 | 24 |
| **F09016** | **B** | **1/160** | 15 | 35340 | 1586 | 2 | 5082 | 945 | 26 | 1593 | 1227 | 48 | 17 | 45 | 607 | 779 | 3033 | 5 | 2317 | 197 | 9 | 102 | 53 |
| **F09016** | **B** | **1/320** | 14 | 25336 | 2355 | 2 | 4595 | 897 | 45 | 1187 | 1914 | 30 | 32 | 51 | 279 | 963 | 1690 | 5 | 3024 | 220 | 98 | 82 | 143 |
| **F09016** | **B** | **1/640** | 12 | 28960 | 2469 | 2 | 5892 | 942 | 109 | 2382 | 2422 | 79 | 71 | 84 | 252 | 1315 | 2047 | 27 | 1703 | 302 | 103 | 105 | 191 |
| **F09016** | **B** | **1/1280** | 13 | 26662 | 3974 | 14 | 6023 | 978 | 131 | 3917 | 3105 | 49 | 59 | 75 | 281 | 982 | 2880 | 119 | 2207 | 374 | 402 | 132 | 322 |
| **F09016** | **B** | **1/2560** | 12 | 20291 | 2847 | 1 | 5413 | 839 | 160 | 4139 | 2919 | 66 | 68 | 113 | 191 | 1337 | 2693 | 131 | 2162 | 465 | 91 | 181 | 321 |
| **F09016** | **B** | **1/5120** | 13 | 19576 | 3355 | 2 | 6781 | 1040 | 220 | 5054 | 2826 | 44 | 111 | 91 | 227 | 1256 | 2599 | 147 | 2230 | 358 | 116 | 151 | 399 |
| **NO SERUM** | **B** | | 12 | 20146 | 2545 | 6 | 5454 | 836 | 248 | 5300 | 3549 | 47 | 145 | 163 | 214 | 1529 | 2601 | 136 | 2408 | 587 | 202 | 153 | 451 |
| **INPUT** | | | 21 | 3473 | 513 | 348 | 4174 | 1332 | 808 | 2027 | 1021 | 328 | 339 | 371 | 528 | 454 | 694 | 438 | 1097 | 686 | 513 | 1179 | 574 |

The example table in Table SI. 1 only shows a subset of variants due to space limitations. In our experiment we can have as many as 136 variants. Each row in the table corresponds to a unique combination of SERUM, DILUTION and REPEAT. Pairs of serum and dilution (such as F09016, 1/20 or NO SERUM, ‘’ ) are called **experiments**. In this case, each experiment (except INPUT, ‘’ ) has two **repeats** A, B. Each repeat is a full experimental repeat, that is, two independent inputs diverging at the point of sampling from an aliquot (so independent serum incubation, plate growth, amplification and sequencing). Finally each row is called a **sample.** We will use upper scripts to denote indices for experiments, repeats and samples where each index runs from 0 to the total number of its unique elements. There are some other terminologies, not used in this SI but inside the model code packages. Samples which are not INPUT are called **assay samples.** Samples which are not INPUT or NO SERUM are called **serum samples**. We define **assay experiments** and **serum experiments** in a similar manner as samples. When there is any risk of confusion with any other usage of the word sample (see section “Model Overview” for one such source of confusion), we will use **table sample**.

When we talk about proportions of variants we talk about row-wise proportions, so dividing the numbers above by the sum of counts in its row (that is the total number of sequences in that sample).

For any quantity with a lower script, such as $f_i$ , the version without the lower script, f, denotes the vector f=($f_1$, $f_2$, …). Similarly if one has a quantity $C^j_i$ then $C^j$=($C^j_1$ , $C^j_2$ , …). Capital letters are generally reserved for experimental observables whereas lowercase letters are used for model parameters, variables and predictions.

#### 1.1 Model Overview

In the Bayesian context, given a set of experimental observables and a prior probability distribution for the parameters of interest, a trained model outputs a posterior distribution for the parameters which are updated (or “fitted”) versions of the prior distributions in the light of the added data. In analogy to classical methods it can be thought of as taking a set of initial parameters and finding optimum parameters. Instead of doing it in a pointwise manner, Bayesian models do this on the level distributions. Such a model can be used to “sample” simulated observables by sampling likely parameters from the parameter posterior distributions and plugging them into the model likelihood for observables. The verb **sample** therefore will be sometimes used for the action of sampling a parameter or a prediction from a Bayesian model. Simulated observables from the model are also called **posterior predictive samples**. A schematic for this process is shown in Fig SI. 1.

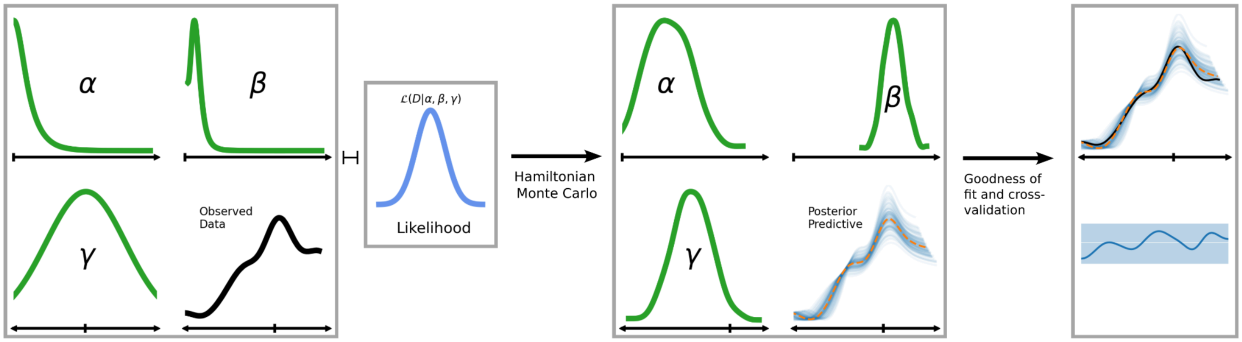


**Fig SI. 1 Overview of the Bayesian model**

A generic Bayesian model starts with initial prior distributions for some parameters, **α, □, ɣ**, … (green, first panel) and distribution of some observed data **D** (black, first panel). These prior distributions are then updated using the data and the chosen likelihood $\mathcal{L}$ (cornflower blue, second panel) to obtain posterior distributions which give the marginal likelihood of parameter values conditional on the data (green, third panel). Updating the priors is usually done through Monte Carlo sampling algorithms such as Metropolis Monte Carlo or Hamiltonian Monte Carlo. Moreover, using posterior distributions for the parameters, a Bayesian model can sample new data **d** (tableau blue for individual samples and orange for the mean, third panel) which are called posterior predictive samples. The posterior predictive samples can be used for goodness of fit (fourth panel first plot) and cross-validation (fourth panel second plot) checks by comparing them to the observed data. These are respectively called posterior predictive checks and leave-one-out probability integral transform.

In this context, the trained Bayesian model becomes a simulator for the experiment (unlike the case of fitting a separate curve to each of the repeats which does not model the in-between repeat noise). Posterior predictive samples can be compared to actual observations in what is called a posterior predictive check, to see how well the model fits the actual data. In such a case if $D^j = (D^j_1, \ldots)$ is a set of observables where j denotes table sample number and $d^{s,j} = (d^{s,j}_1, \ldots)$ are simulated observables from the trained model where s denotes the posterior predictive sample number, then one compares the distribution of the former to a selection (with respect to index s) of densities from the latter (see Fig. SI 1, last panel, first plot where black line corresponds to distribution D, individual blue lines correspond to samples $d^{s,j}$ for s=1,...100 and orange dashed line corresponds to the mean distribution of posterior predictive samples).

Note that in essence the goodness of fit check is the process of creating s=1,... simulations of the observables and looking at how far each distribution is to the distribution of observables. These goodness of fit checks are called **posterior predictive checks** (PPC). This is a visual test which can be complemented with plotting the density of residuals. A more qualitative cross-validation test known as leave-one-out probability integral transform **LOO-PIT** can be performed which is known to be very sensitive to the model misspecification (Nguyen et al. 2025). It essentially approximates the CDF of the posterior predictive samples when one observable is left out and then applies a probability integral transform to the experimental observables using this CDF for each leave-one-out case. If the model is suitable for the data at hand then the transformed densities should look uniform (i.e straight). Confidence intervals showing an acceptable level of variation from a straight line can also be used to make the comparison more quantitative rather than visual. As in the classical case, when the number of observables increases, the confidence interval gets narrower and therefore even minute variations of the data from the model can be detected. If there is a bias in the model it generally reflects itself in the form of sloped lines rather than straight-like (see Fig SI. 1, last panel, dark blue line). Underdispersed and overdispersed models result in lines that respectively look like smiling or frowning. Since we have repeat data, repeat errors can be used to see if model predictions match experimental noise. In particular histogram of model prediction residuals can be compared to histogram of experimental repeat differences. If the former histogram has a mean significantly higher than 0, it indicates a bias whereas if it is more or less dispersed than the latter distribution, it indicates that the model is respectively under or overfitting. One should also note that vanilla LOO-PIT is only suitable for continuous distributions whereas we use count distributions. Arviz allows handling of discrete data via a randomized tie-breaking procedure, which is what we employ here.

Apart from these, one also needs to check model convergence to make sure that the estimates we are getting are reliable. Steps involved in convergence checks are described in SI Section "Diagnostics"

#### 1.2 Some examples of repeats

Before we start describing the model, we look at some repeats from the data. Fig SI. 2 shows scatter plots of repeats of sequencing variant percentages obtained from several different experiments. Note that the examples used here won't always be full experimental repeats as shown in the table Table SI. 1. These repeats were done for the purpose of studying the sources of noise in the experiment and for instance include sequencing repeats.

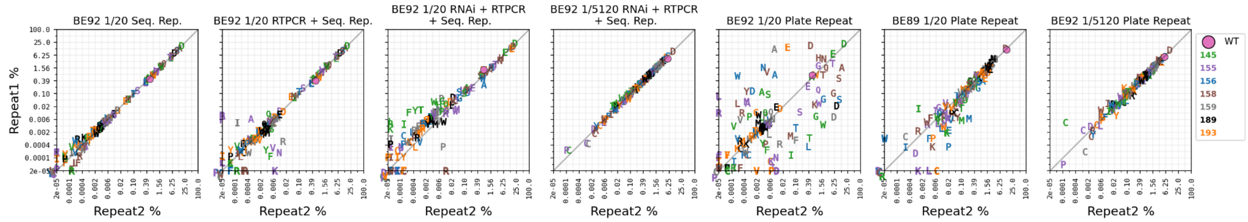


**Fig SI. 2 Repeat virus %s for various samples**

The plots show the scatter plot of virus % (in $\log_2$ scale) for two repeats of various experiments as indicated in the title. Each repeat contains 134 viruses. Seq. Rep. means sequencing repeat and includes repeating index ligation and 8-15 rounds of PCR + sequencing + read processing from a common sample. RTPCR is the reverse transcription which is for amplification of common isolated RNA samples. RNAi is the RNA isolation from independent samples collected from common supernatants. Plate Repeat means a fully independent repeat starting from the initial sampling of viruses. BE89/BE92 in the titles are the sera used and 1/20, 1/5120 indicate dilutions. %s which are either zero on the x or y axis (but not both) are shown on the axis. %s which are zero in both axes are shown at the bottom corner, with a jitter to place them slightly out of the plot.

It is clear that the experiment accommodates various sources of noise in a quite complex manner. It is possible to deduce that the “measurement noise” (those coming from RNAi, PCR, RNAi) is generally more prominent on variants whose proportions are on the order of $10^{-4}$ or less regardless of sample. The largest contributor to noise are plate repeats in the case of low dilutions of highly reactive sera (such as BE92). Indeed, we see that plate repeat noise for BE89 1/20 or BE92 1/5120 is nowhere near what is seen for BE92 1/20. This is the stochasticity of high neutralisation in which nearly all of the PFUs are neutralized and therefore proportions of variants are highly unpredictable (considering that neutralisation equilibrium is not a static state but a dynamic one where antibodies continuously associate and disassociate as well as other sources of noise such as replication and transcription errors). It would be a fruitless endeavor to try to capture the experiment in its full glory with a model that individually addresses each type of noise present (such as a mixture model). Instead we will try to aim for a relatively simple model by showing that a certain type of likelihood can capture the combined noise present in the system sufficiently well. To describe this likelihood, we study in the next section, the simple case of no serum (so no neutralisation) and where only replication fitness matters.

#### 1.3 Choice of Likelihood

In this section we establish why the choice of a BetaBinomial likelihood is more appropriate for the problem at hand, while also presenting a gentle introduction to the full model by considering a simpler case where no neutralisation is involved and we only consider replicative fitness. So in this case we only have two rows of data; table samples corresponding to rows where SERUM value is equal to NO SERUM in table SI. 1. They will be indicated by the upper index $^r$ which can also in this case be thought of as the repeat index for repeats A and B.

In modelling any experiment involving count data, binomial, poisson or multinomial likelihoods are commonly used. Whereas these models can describe the sampling noise associated to different levels of proportions (Fig SI. 2), they are insufficient to describe other experimental effects that might lead to overdispersion. In such cases BetaBinomial and Dirichlet Multinomial likelihoods can be used. Even though Dirichlet Multinomial likelihoods allows modulating the overall noise that acts on all the categories of a counting process, if more flexibility is needed collections of univariate BetaBinomials (same number as the number of variants in the model) can be used to modulate the noise further on a per category basis. Such collections of univariate distributions are sometimes called "batched" distributions. This would be akin to comparing a Multinomial with n categories vs a batched collection of n Binomials. BetaBinomial is a Binomial distribution where p is not a free parameter but is instead drawn from a Beta distribution with parameters α, β. Therefore, most commonly used parametrization for BetaBinomial is an **n, α, β** parametrization (where n denotes total count and α, β are the parameters of the Beta distribution). For our purposes we employ a **n, p, c** parametrization in which **α = c x p, β = c x (1-p).** With this parametrization, the mean of the BetaBinomial simply becomes n x p (as in the Binomial) and variance becomes p x (1-p) x n x (c+n)/(c+1) (where as Binomial would be p x (1-p)). In this way, the relation between BetaBinomial and Binomial becomes very clear. BetaBinomial with parameters n,p,c represents a distribution which has the same mean as a Binomial with parameters n,p but larger variance depending on c. As c goes to infinity, the variance approaches the regular Binomial variance. In the following we generally call c the **concentration** parameter (inverse of dispersion).

In our context, the experiment we want to model is a mixture of $\mathbf{N_v}$ viruses ($_v$ is not an index but a label in this case) with **input** proportions $\mathbf{p^0_i}$ (i=1,..,$N_v$) which are then grown on a plate. To obtain the final proportions, the output counts of each virus are measured in the form of sequencing counts and divided by total sequence counts. Two full repeats are carried out leading to data $\mathbf{C^r_i}$ (r=1,2 and i=1,...$N_v$). Let $\mathbf{T^r = \Sigma_i C^r_i}$ be the total number of counts for each repeat and $\mathbf{f_i}$ be the parameter indicating the fitness of the variant i.

To see if the flexibility of modulating the noise in BetaBinomial is required, we compare different versions of the Binomial models against each other. In particular we compare BetaBinomial with variable concentration (that depends on proportion as described below) vs a constant concentration BetaBinomial and the regular Binomial distribution. The three likelihoods are given as below:

$$p_i = \frac{p^0_i \times f_i}{\sum_{k=1}^{N_v} p^0_k \times f_k}$$

$$C^r_i \sim Binomial\left(n = T^r,\ p = p_i\right)$$

$$C^r_i \sim BetaBinomial\left(n = T^r,\ c,\ p_i\right)$$

$$C^r_i \sim BetaBinomial\left(n = T^r,\ c\left(p_i\right),\ p_i\right) \quad (0)$$

Here c without brackets denotes a free concentration parameter used to modulate the amount of noise whereas $c(p_i)$ denotes a concentration parameter that also depends on proportion (such as a linear dependance). Note that in the context of Bayesian modeling one needs to specify priors for parameters such as $\mathbf{f_i}$ and $\mathbf{c}$ or $\mathbf{c(p_i)}$. We leave the definition of these priors to section **1.4 The Full Model** where we describe the full model. However our general approach to specifying priors is using weakly informative priors as much as possible. The attentive reader will have noticed that the proportions $p_i$ are invariant under the scaling of the replicative fitnesses: $f_i \rightarrow \alpha f_i$ . This makes sense because proportion differences only define relative replicative fitness. This results in a non-uniqueness in the model which can be remedied by adding another likelihood which controls for the total amount of change of virus content in these assays where observables in our case will be qRT-PCR ct values. This will be explained in detail when the full model is presented.

We evaluate these models by looking at PPC, residual and LOO-PIT plots (see section 1.1 Model Overview). Of particular interest are the following questions: 1- How well each model captures the proportion variation in repeats, 2- how well each model fits the data, 3- how well each model cross-validates. We answer these three questions by looking at four different data sets in Fig. SI 3.

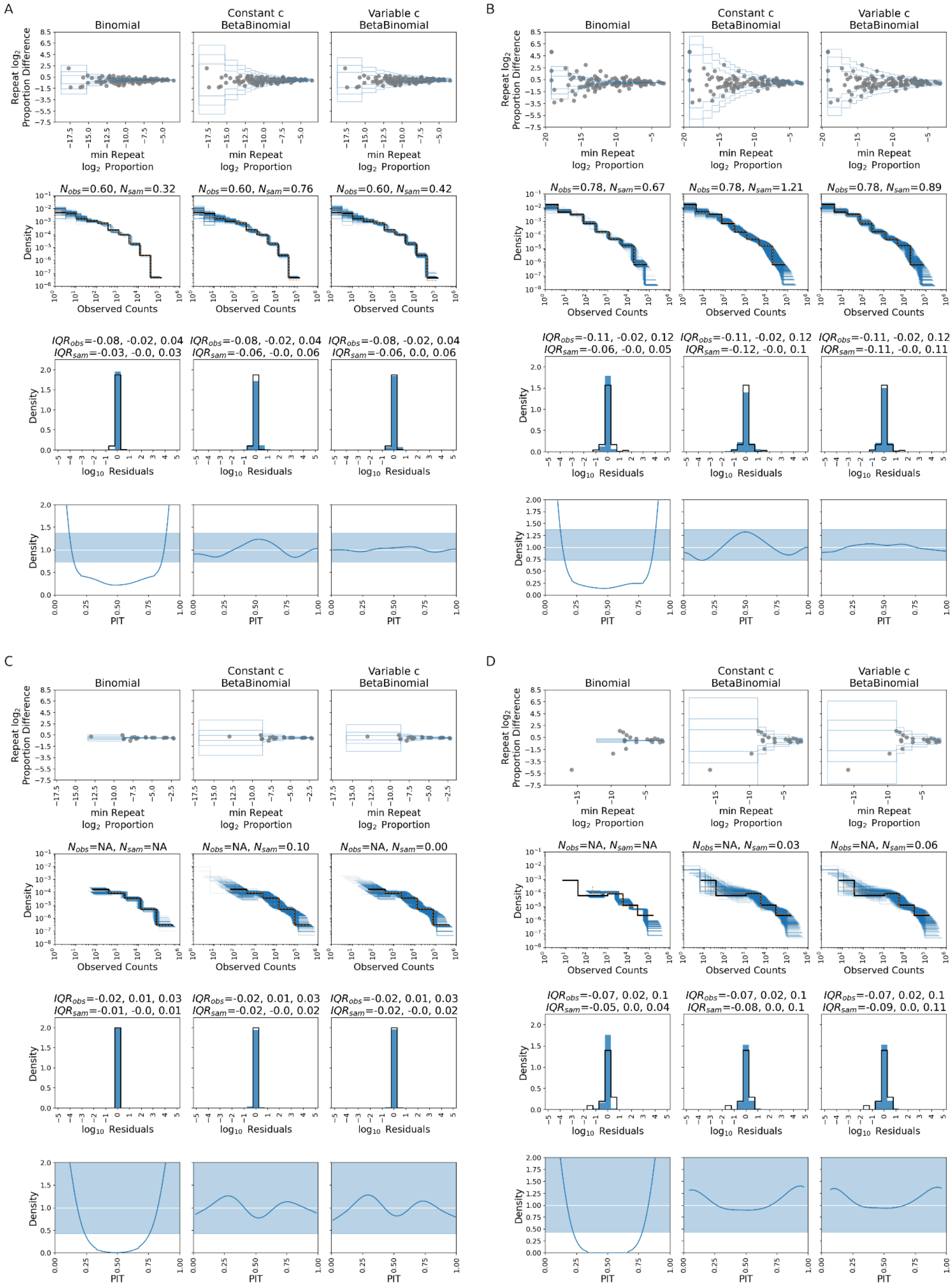
A
Binomial
Constant c BetaBinomial
Variable c BetaBinomial
Repeat log2 Proportion Difference
min Repeat log2 Proportion
Nobs=0.60, Nsam=0.32
Nobs=0.60, Nsam=0.76
Nobs=0.60, Nsam=0.42
Density
Observed Counts
IQRobs=-0.08, -0.02, 0.04 IQRsam=-0.03, -0.0, 0.03
IQRobs=-0.08, -0.02, 0.04 IQRsam=-0.06, -0.0, 0.06
IQRobs=-0.08, -0.02, 0.04 IQRsam=-0.06, 0.0, 0.06
log10 Residuals
PIT
B
Binomial
Constant c BetaBinomial
Variable c BetaBinomial
Nobs=0.78, Nsam=0.67
Nobs=0.78, Nsam=1.21
Nobs=0.78, Nsam=0.89
IQRobs=-0.11, -0.02, 0.12 IQRsam=-0.06, -0.0, 0.05
IQRobs=-0.11, -0.02, 0.12 IQRsam=-0.12, -0.0, 0.1
IQRobs=-0.11, -0.02, 0.12 IQRsam=-0.11, -0.0, 0.11
C
Binomial
Constant c BetaBinomial
Variable c BetaBinomial
Nobs=NA, Nsam=NA
Nobs=NA, Nsam=0.10
Nobs=NA, Nsam=0.00
IQRobs=-0.02, 0.01, 0.03 IQRsam=-0.01, -0.0, 0.01
IQRobs=-0.02, 0.01, 0.03 IQRsam=-0.02, -0.0, 0.02
IQRobs=-0.02, 0.01, 0.03 IQRsam=-0.02, -0.0, 0.02
D
Binomial
Constant c BetaBinomial
Variable c BetaBinomial
Nobs=NA, Nsam=NA
Nobs=NA, Nsam=0.03
Nobs=NA, Nsam=0.06
IQRobs=-0.07, 0.02, 0.1 IQRsam=-0.05, 0.0, 0.04
IQRobs=-0.07, 0.02, 0.1 IQRsam=-0.08, 0.0, 0.1
IQRobs=-0.07, 0.02, 0.1 IQRsam=-0.09, 0.0, 0.11

**Fig SI. 3 Goodness of fit and cross-validation checks on three different likelihoods across four experiments.**

In the figures above, each panel A, B, C, D indicate different datasets. The datasets A, B both have 134 variants and 10000 PFUs whereas C, D have 19 or 20 variants and 10000 or 1000 PFUs respectively. Datasets A and C are what are called mix1-ALL and mix1-posn145 in the main manuscript whereas B and D are mix2-ALL and mix2-posn145. Within each panel, each column indicates one of the three different likelihood models (indicated in the title of the first row) and each row within a panel shows a different type of diagnostic. Let $P^r$ be the vector of observed proportions where r=1,2 is the repeat index and $p^{s,r}$ the vector of posterior predictive proportions where s is the model posterior predictive sample number. Within each of the subpanels, the grey disks in the first row show the scatter of vectors = $\min_r \log P^r$ vs $Y = \log P^1 - \log P^2$ i.e scatter of minimum log repeat proportion vs their difference for each variant. One can think of blue lines as level sets of a 2D density fitted to a collection of 2D sampled $(X^s,Y^s)$ data from the model. They show the conditional lower and upper quantiles of the bivariate distribution $(X^s,Y^s) = (\min_r \log p^{s,r}, \log p^{s,1} - \log p^{s,2})$. Lower and upper quantiles given by $q_{low}$ = (0.01, 0.05, 0.2) and $q_{up}$ = (0.8, 0.95, 0.99) (so it represents how the model posterior predictive samples are spread when transformed as above). Conditioning is done by dividing the X into 20 (for A, B) or 7 (for C, D bins and then computing various quantiles of the Y values for data within these X ranges. The second row in each panel shows posterior predictive check (PPC) plots. In the PPC plots, the black line shows the distribution of observed counts $C^r$ (r=1,2 combined), the orange line shows the distribution of a combination of 200 randomly selected posterior predictive samples for counts ($c^{s,r}$) and the blue lines show the distribution of each individual posterior sample from the same set. The bins for the orange and black lines are identical and are obtained by using the get_bins functionality from the ArviZ (Kumar et al. 2019) package on the $\log_{10}$ transformation of the observed counts. The bins for each blue line are determined individually using the same method. Histograms are plotted using histogram function from the ArviZ package and plotted in logarithmic scale for both axes to be able to visualize finer discrepancies better. Titles show the $\log_{10}$ number of zeros in the observed and sampled counts (mean over the different samples in the latter case) and NA if no zeros. The third row shows the histogram of $\log_{10}$ residuals $r^{s,r} = \log_{10}(P^r) - \log_{10}(p^{s,r})$ as the black line and distribution of repeat errors $\log_{10}(P^1) - \log_{10}(P^2)$ as the blue bars (when proportions are non-zero). A uniformly spaced set of bins from -5 to 5 (inclusive) with steps of 0.5 is used. Titles show the interquantile range of the distribution in both cases. The final plot shows the LOO-PIT using the loo_pit function from the ArviZ package's arviz_stats module and random tie-breaking of discrete data as implemented there. The blue band shows the %99 HDI for PIT of uniform distributions with the same number of observations. The dark blue line shows the probability integral transform of leave-one-out observables (Nguyen et al. 2025). For a good model the dark blue line is expected to be as straight as possible and within the blue band as much as possible.

Fig. SI 3 suggests multiple implications. By looking at the first column of each panel, we see that the Binomial model underestimates the amount of repeat noise. This is implied by the first, third and fourth plots in those columns (the smile pattern in LOO-PIT is indicative of model under-dispersion, whereas frown over-dispersion). If we compare constant and variable BetaBinomial models in terms of cross-validation (last two rows in each panel), where as they perform similar for the models with low number of strains (C,D) the constant BetaBinomial model overestimates the noise in the case of high number of variants (A,B). Even though the LOO-PIT plots in the last row suggest that this over-estimation is within acceptable limits, the discrepancy will become more severe when we consider more complicated scenarios where serum is involved (see section Diagnostics). All of the models generally perform well for the goodness of fit checks (with possibly the exception of Binomial model for dataset D). Therefore the correct choice of model in this case is more about correct estimation of uncertainty intervals rather than point estimates of parameters involved. In summary, the binomial model underestimates the noise and constant concentration BetaBinomial model overestimates it (more for the lower proportions as can be seen by the first row of plots). The variable concentration model is formulated to fix the overdispersion of BetaBinomial model for very low proportions which are more likely to appear when the number of variants are higher and therefore it makes sense that they perform similarly for cases C and D. Nevertheless, we keep both BetaBinomial (constant and variable) models in the full model package (Tureli 2025d) and suggest running one or both depending on the number of strains. After we have described the full model and applied it to fit titers as well as replicative fitness parameters, we will do further goodness of fit, cross-validation and diagnostic analysis on both of these models to strengthen the conclusion that BetaBinomial models with variable concentration perform well for this type of data.

#### 1.4 The Full Model

The main components of the full model are replicative fitness, neutralisation and concentration (see section Choice of Likelihood for the definition of the concentration parameter which can be thought of as the inverse of noise). Compared to the only replicative fitness + concentration model described in the previous section, the main differences are the parameters associated with neutralisation. In this section lower case **i** denotes variant index, lower case **j** denotes serum index. For each **i** and **j** pair we have a titer parameter $\mathbf{t_{ij}}$ which gives the titer of variant **i** against serum **s**. For each serum **j** we have a single slope parameter $\mathbf{s_j}$. The left and right asymptotes of the neutralisation titer curves are respectively fixed at 1 and 0. Both $\mathbf{t_{ij}}$ and $\mathbf{s_j}$ themselves are actually priors so they are not “flat” parameters in the classical sense but distributions that allow a reasonable range of values for these parameters. They will be described after the likelihood has been defined. As in the previous section, **c** and **c(p)** denote the concentration parameters which are either constant or depend on proportions (functional form of c(p) isde. We will let **r** denote the table sample index. So $\mathbf{C^r_i}$ in this context denotes the observed count of the $\mathbf{i^{th}}$ virus in the $\mathbf{r^{th}}$ sample (see SI Section **The Data Structure Overview**). We let j(r), d(r) denote respectively the serum and dilution associated with sample r (which can also be no serum and undefined dilution, see Table SI. 1). Let $\mathbf{n^r_i} = \mathbf{n}(t_{ij(r)}, s_{j(r)}, d(r))$ denote the neutralisation amount of virus **i** for serum **j(r)** at dilution **d(r)** (equal to 1/(1+exp(-(d-t)*s)) with indices dropped). Note that if $r_0$ and $r_1$ are two repeats of the same serum with the same dilution then, $\mathbf{n^{r0}_i} = \mathbf{n^{r1}_i}$ because $j(r_0) = j(r_1)$ and $d(r_0) = d(r_1)$. So their neutralisation is identical. The repeat variation matching the one in the observed data will therefore be modelled by the variation of the BetaBinomial likelihood and not by employing different titer parameters for each repeated curve. The likelihood is then given by

$$p^r_i = \frac{p^0_i \times f_i \times n^r_i}{\sum_{k=1}^{N_v} p^0_k \times f_k \times n^r_k}$$

$$C^r_i \sim BetaBinomial\left(n = T^r,\ c\left(p^r_i\right),\ p^r_i\right) \tag{1}$$

where as before $\mathbf{T^r = \Sigma_i C^r_i}$ and BetaBinomial is parameterized in terms of concentration and mean as described in the previous section. As also noted in the section before, we need a likelihood to characterize the change in total virus amounts to remedy the scaling invariance of $\mathbf{f}_i$ and $\mathbf{n}^r_i$ . This is given by the following prior + likelihood

$$\Delta ct^r \sim Normal(\sum_{k=1}^{N_v} p^0_k \times f_k \times n^r_k, \sigma_{ct}) \quad (2)$$

where $\sigma_{ct}$= 0.4 and $\Delta ct^r = ct^0 - ct^r$ is a measure of how much the ct value changes with respect to some baseline value $ct^0$ (such as ct value of the input of the same volume as the volume that was used to measure the ct for the sample) after the assay is done. The exact value of the baseline ct value is irrelevant since it will affect all the samples in the same way and will just end up shifting the replicative fitness values.  The standard deviation of the ct distribution does infact seem to increase with lower PFUs, possibly due to stochasticity and the experimental_models module in our package allows us to model the data using this approach. We have not found a significant difference in terms of the important parameters; the titers and replicative fitnesses, or any improvement for cross-validation of count data. It does however improve the fit for Δct observed values.

#### 1.4.1 Replicative Fitness Prior

For specifying the prior distributions for the replicative fitness values we follow a weakly informative approach. We first estimate the replicative fitness non-parametrically using the input proportions, no serum proportions and Δct. This is done by computing $\log_2$(no serum proportions) - $\log_2$(input proportions) + $ct_0$ - $ct_{noserum}$ for each repeat and then taking the average over repeats. Let $\mu_f$ and $\sigma_f$ denote respectively the population mean and standard deviation of these values. Then we define parameters $f_i^{offset}$, $f^\mu$ by

$$log_2(f^{\ offset}) = ZeroSumNormal(0, 1)$$
$$log_2(f^{\mu}) = Normal(\mu_f, 0.5)$$
$$log_2(f^{\mu}) = log_2 f^{\mu} + \sigma_f \times log_2 f_i^{offset} \quad (3)$$

So replicative fitness values are decomposed as a population bias + population standard deviation   x  individual zero sum variance. Note the factor $\sigma_f$ assures that the priors capture the natural scale of the distribution of these parameters and is weakly informed by the nonparametric estimate of the population average (individual replicative fitness nonparametric estimates are not used in defining the priors).

#### 1.4.2 Neutralisation Curve Prior

In order to define $\mathbf{n}^r_i$, we have to define priors for titers $t_{ij}$ and slopes $\mathbf{s}_j$. The slope prior is

$$s_j \sim 0.8 + \gamma^{-1}(0.5, 0.25) \quad (4)$$

Here $\gamma^{-1}(\mu, \sigma)$ is the inverse gamma distribution in mean, standard deviation parametrization. We also impose an upper bound of 5 on the value of the slope. Note that neutralisation curve slopes tend to be realistically around values of 0.5 to 2. Therefore this could be considered somewhere between an informative and generically weakly informative prior.

To formulate the titer priors, we follow an approach similar to the replicative fitness. We first non-parametrically estimate the titer values for each serum and then compute population mean and standard deviation from these. To estimate the titers non-parametrically, we use the Spearman-Kärber method (Miller 1973; Kärber 1931; Spearman 1908, 1909). This method requires that the neutralisation data is monotonic which is not always the case with noisy data, so we follow the iterative method of Ayer-Brunk-Ewing-Reid-Silverman (Ayer et al. 1955) to monotonize the neutralisation data. These are available as a part of the package Thor (Tureli 2025d). Note that these results are only used to get rough estimates for the population mean and standard deviation of titers so that generically weakly informative priors can be defined. Let $\mu_j$ and $\sigma_j$ denote the population mean and standard deviations of titers for serum j. Then

$$t_{ij} \sim Normal\left(\mu_j, \sigma_j\right) \quad (5)$$

If the range of dilutions for the serum is $d_0$ to $d_1$, then we impose the lower and upper limits of $d_0 -1.5$ and $d_1+1.5$, as trying to extend the calculations too much beyond this range where we have no data would be unreasonable. Then the neutralisation is given as

$$n^r_i = 1 - Sig\left(\left(d(r) - t_{ij(r)}\right) \times s_{j(r)}\right) \quad (6)$$

where **j(r)** and **d(r)** are respectively the serum index and dilution of table sample r. Sig(x) is the sigmoid function which is equal to $1/(1+e^{-x})$.

#### 1.4.3 Concentration Prior

Concentration represents the amount of noise in the likelihood. The smaller the concentration the higher the noise. As we have seen in the section "Choice of Likelihood", a constant concentration parameter per table sample might overestimate the amount of noise. So for each table sample **r**, we define a linear function (see formula below) which has a right value $\mathbf{ra^r}$**,** left value $\mathbf{la^r}$ **(**with the constraint $\mathbf{la^r > ra^r}$**)**, and the independent variable **x** of the linear function is $\log_{10}$ proportion. $la^r$ is the value the linear function attains at floor(min($\log_{10}$(proportions))) and ra is the value of the linear function at 0 (i.e log10 of proportion of 1 which is the max attainable value). So the concentration increases as the proportion decreases. However, due to the form of BetaBinomial, the relative noise on proportions also increases as proportion decreases. So these two effects combined together creates an effect where the noise still increases as proportions decrease but in a more controlled manner (see blue lines for variable c in the first row of plots in each panel of Fig. S3). See also SI section “Relation Between Concentration, PFU and Neutralisation” for a more detailed analysis of the relation between concentration and different table samples. The concentration function is then defined as follows:

$$ra^r \sim \gamma^{-1}(7,\ 4)$$
$$la^r \sim \gamma^{-1}(7,\ 4)\ ,\ la^r \geq ra^r$$
$$log\ c(p^r_{\ i}) \sim ra^r + (la^r - ra^r) \times log_{10}(p^r_{\ i})/log_{10}(p_{min}) \quad (7)$$

Therefore each table sample set has a common **$ra^r$, $la^r$** defined which applies to all of the variants and the dependence of the concentration on the variant only comes from the proportion. Here pmin is floor($min_r$ ($1/N^r_{seq}$)) where $N^r_{seq}$ is the sequence number of table sample r. Also the priors are defined such that each collection of repeats (called an experiment, such as two repeats for dilution 1/20 of a given sera, see SI section “Data Structure Overview”) share the same **$ra^r$, $la^r$** values.

Note that the formula above defines the log concentration. Values of **$la^r$** or **$ra^r$** much higher than 15 would describe unreasonably high concentrations. The highest fitted values we have observed for **ra** are around 8-9 but occasional sampling from higher values can cause sampling slow down and divergences. A value of 15 basically represents the null hypothesis that there is no extra noise on the system apart from the noise given by a Binomial distribution. Values less than 0, while possible, are extremely diffuse leading to likelihoods that concentrate on borders away from the expected values. Therefore we bound **$ra^r$** and **$la^r$** between 0 and 15.

#### 1.4.5 Input Calibration

Counts obtained from the input sample are modelled using a Multinomial likelihood. This is because in the case of the input sample, there is no sources of dispersion (like sampling bottle necks) that are observed on the other samples. The prior and likelihood describing the input is given by

$$p^0 \sim\ DirichletMultinomial(a\ =\ 5)$$
$$C^0 \sim Multinomial(p\ =\ p^o,\ N\ =\ T^0) \quad (8)$$

The experimental methods of generating input mixtures differ between the mixtures mix1-ALL and mix2-ALL. In the former one input mixtures for each of the seven positions 145,155,156,158,159,189 and 193 are generated separately (each including root). These individual input mixtures are then combined with weights calibrated inversely to their PFU/ml (A). After this initial mixture is created, it is sequenced, and the mean $log_2$ proportion of the 19 variants at each position is computed. A second mixture is then created, this time using weights inversely to these proportions (Fig SI. 4B). In contrast, the latter mix uses only PFU/ml based calibration (Fig SI. 4C) and each position's mixture does not contain the root, which is generated individually and also mixed with others based on PFU/ml. The rationale for this simplification is that it is possible to have positions where PFU/ml is low but sequencing proportion is high compared to other positions (i.e, possibly high levels of non-infectious particles). A calibration as in the first method can then create a bottleneck for such input mixtures where the PFU/ml is lowered which could result in more subsampling variance. We demonstrate the benefits of this approach in the SI section "Further Diagnosis of Model Fits". Of the 14 individual position mixtures created this way, we have seen three outlier cases of PFU/ml values not matching the proportions in sequencing (position 193 in Fig SI. 4A and positions 145, 155 in Fig SI. 4C). These discrepancies seem to stem from viable to non-viable virus proportions in each mixture. Indeed when the input from mix1-ALL (Fig SI. 4A) is compared to input from mix2-ALL (Fig SI. 4B), we see a bias for positions 145, 155 being higher but when they are grown on a plate this bias disappears (Fig SI. 5), for which the simplest explanation is that the non-viable viruses leading to proportion inflation in sequencing disappear during the assay as they either get washed away without being able to attach or they fail to replicate. This is supplemented further by comparison of PFU/ml and -ct values (Fig SI. 5) which single out 145,155 variants as having much higher -CT/pfu/ml ratio than other viruses.

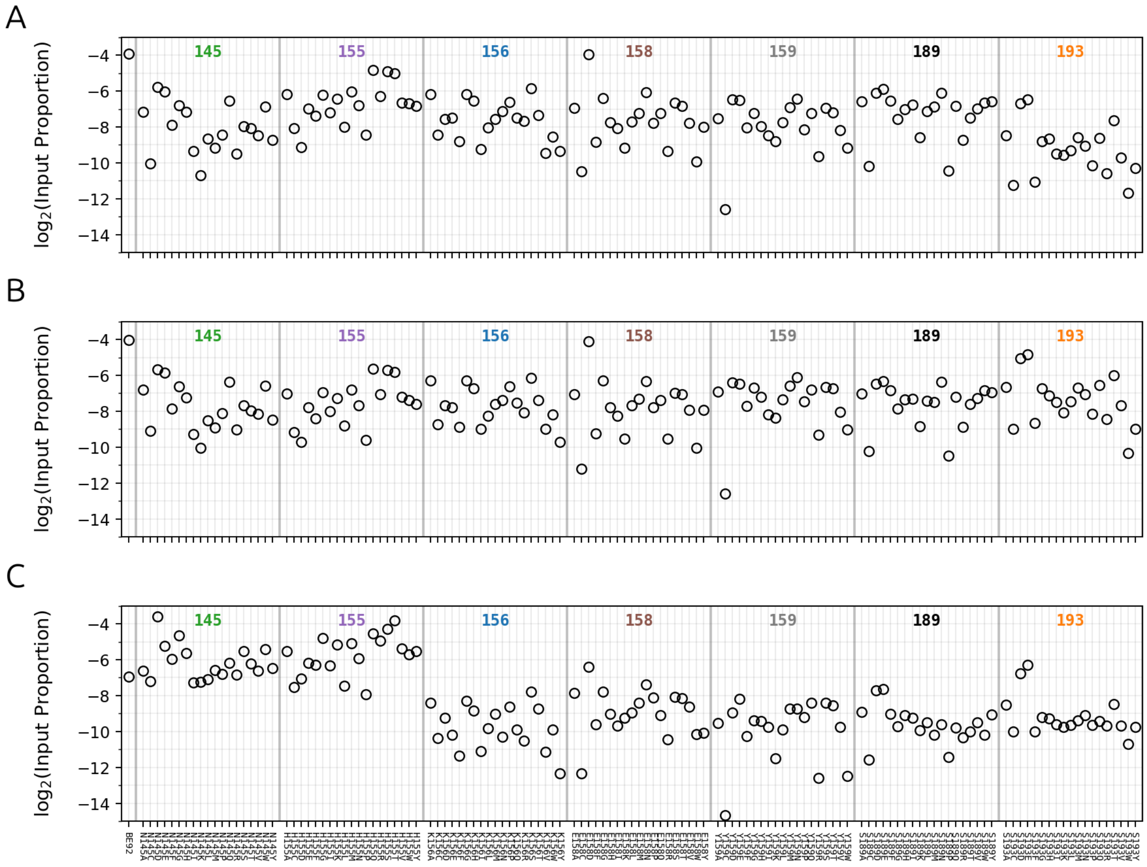

**Fig SI. 4 Comparison of input proportions from different mixtures.**

Each plot shows the input proportions obtained from different methods of mixing seven position mixtures and root. **A.** Mixing is done by PFU/ml, **B.** Mixing is done by sequentially PFU/ml and sequencing proportions, **C.** Mixing is done by PFU/ml. The rescue method for A and C differ (mix1-ALL vs mix2-ALL). The latter case is optimized to reduce effects of replicative fitness on proportions (see “Virus libraries amplification in MDCK cells” in “Experimental Method Details”). Horizontal lines and numbers at the top indicate groups of positions with the first marker being the root variant BE92.

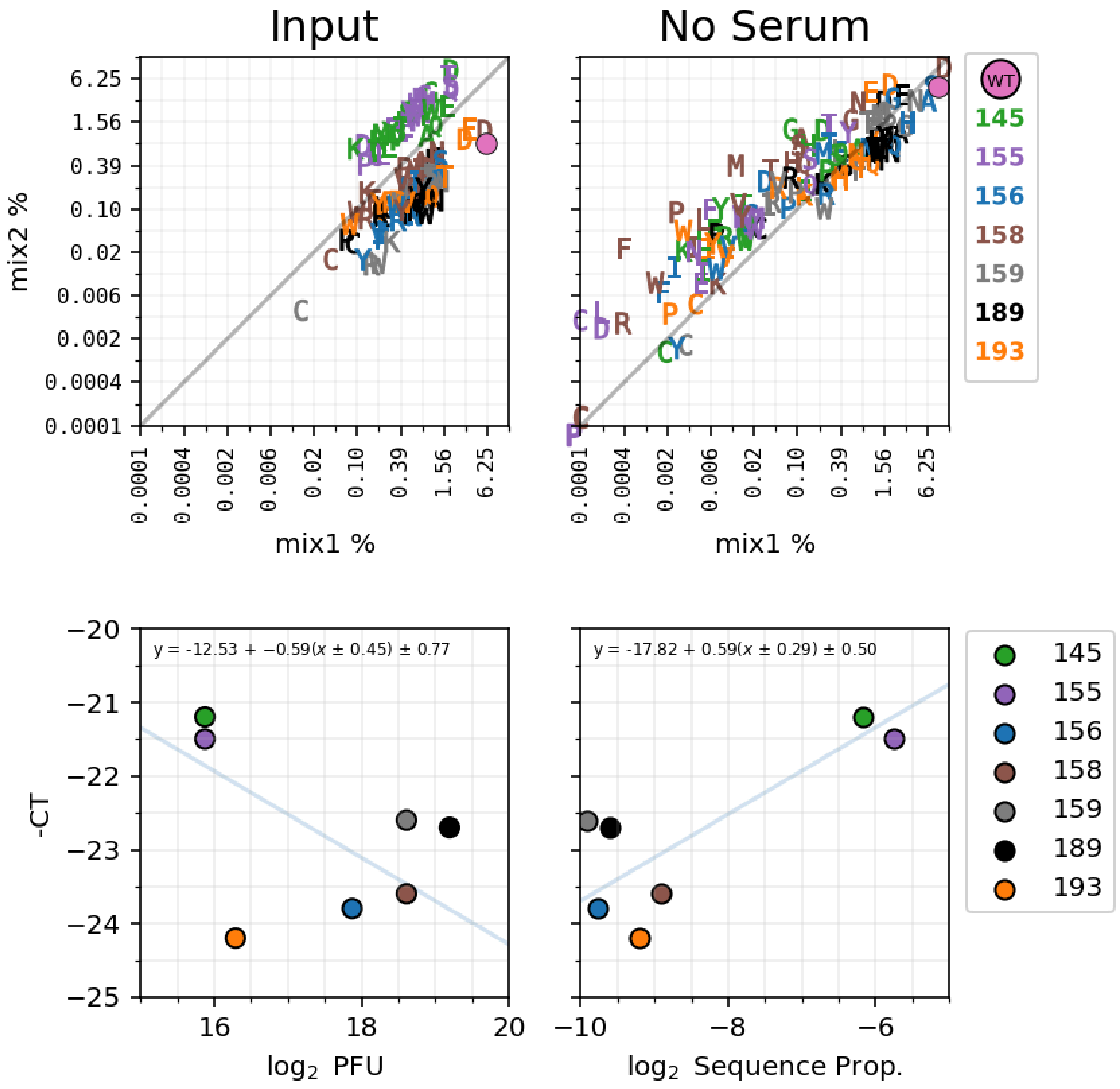


**Fig SI. 5 Fleshing out various discrepancies caused by viable to non-viable virus ratios.**

First two plots show respectively the comparison of variant %s for the input and no serum post assay mixtures for mix1-ALL and mix2-ALL. The last two plots show comparison of -CT vs $log_2$PFU and -CT vs $log_2$ Sequencing

Proportions for each position's mixture which was used to construct the mixture mix2-ALL. Sequencing proportion is the position-wise proportions in Fig SI. 4C. Blue lines and the inset equations at the top show orthogonal regression.

Therefore, in cases where one observed too much discrepancy between PFU/ml and sequencing proportions of constituent mixtures, we allow a further input to the model which calibrates this. This corresponds to renormalizing the estimated mean $\log_2$ proportion of individual positions' mixtures so that they all have the same value. In essence for mix2-ALL this boils down to adding a fix covariate vector (called ppfu_ratios in the model function) so that the likelihood in Equation 8 is changed to

$$C^0 \sim Multinomial(p = p^o \, x \, ppfu, \; N = T^0) \qquad (9)$$

In the case of mix2-ALL this is 9.95 for position 145 variants, 13.20 for position 156 variants and 1 for everything else (these are estimated from the proportions seen in Fig SI. 4C). This has the effect of lowering the estimated $p^0$ values for these positions for the case when we would not use this calibration. This calibration has no effect on titers as they depend only on serum and serum samples; it only changes replicative fitness values (on a position-wise level). In order to test whether this calibration is improving replicative fitness values, we compared three cases: mix1-ALL, mix2-ALL without ppfu ratios and mix2-ALL with ppfu ratios by scatter plotting model estimated means for replicative fitness values against observed plaque sizes (Fig SI. 6). When comparing mix2-all without and with ppfu ratios (second and third panels), one sees that even though the correlation coefficient is similar, the within group standard deviation significantly decreases for each group which is highly suggestive of better grouping. Indeed the culprits causing high standard deviation in mix2-ALL without ppfu (second panel) are mostly green (145) and brown (158) which show a tendency for being lower than expected (as compared against a plaque size) which is indicative of the fact that they might have higher input proportions in sequencing then they would normally have if only viable viruses were considered (higher input proportions lead to lower estimated replicative fitness).

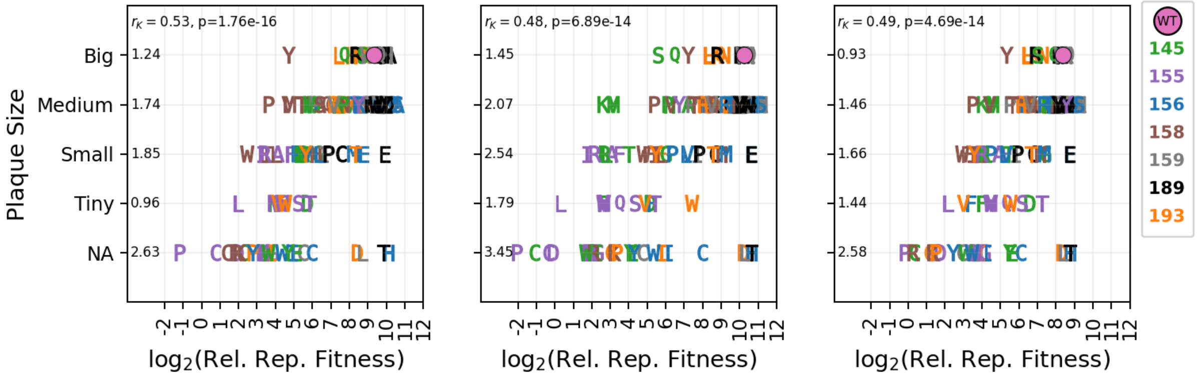


**Fig SI. 6 Comparing replicative fitness measured with various input mixtures to plaque sizes.**

Panels show model mean $\log_2$ replicative fitness values (x-axis) vs plaque sizes (y-axis) for each variant respectively for input mixtures mix1-ALL, mix2-ALL without ppfu ratios, mix2-ALL with ppfu ratios. Legend indicates color of the position and markers indicate variant. NA value on the y-axis indicates that the virus could not be individually rescued or no plaque visible. Inset value at the top shows Kendall $\tau_B$ correlation coefficient and its p-value. The other values along the y-axis show the standard deviation of the x-values within each group.

This analysis demonstrates the benefit of input proportion calibration when there is too much discrepancy between measured PFU/ml and sequencing proportions. It has the caveat that it can only be applied to groups of viruses, we have for instance not attempted to calibrate the root variant's mixture's proportions in the mix2-ALL because that consists of a single virus and no reliable group-wise estimate can be obtained for that virus. This is an aspect of the assay that is in active development and we plan to test various other approaches such as combining position plasmid mixtures (rather than combining them after rescue + passage) and rescuing them together which might uniformize the viable to non-viable virus bias that is observed for each position.

#### 1.4.6 Total Reaction Volume Calibration

In the experiments performed with mix2-ALL, the reaction volume of reagents for the serum incubation was different. The general procedure is to mix 100ul virus with 100ul sera. However, due to lower PFUs/ml of the sample for some of the constituents of this mixture, we mixed 214.2ul virus with 100ul sera. In this setup, the same total amount of PFUs and antibodies are then incubated in a larger reaction volume (314.2ul vs 200ul) which changes the forward reaction rate. This change can be estimated by $\log_2((a/200*b/200)/(a/314*b/314)) \sim 1.3$ (where a and b are total amounts of viruses and antibodies). This reduction in reaction rate mimics a $2^{1.3}$ fold increase in dilution. We allow incorporating this into the model by a xshift parameter (which we set to -1.3 in this case) which gets added to the log transformed dilution covariates. This adjustment is found to increase the consistency of magnitude of titers when comparing variant titers obtained from the only 145 position mixture vs ALL positions mixture (former of which was incubated following the standard volumes in the protocol as it was a single position mixture and did not require larger volumes than 100ul).

### 2 DIAGNOSTICS

#### 2.1 Model Convergence Diagnostics

There are several important diagnostics one can use to check the convergence, goodness of fit and cross-validation for Bayesian models. $\hat{r}$ and Effective Sample Size (ESS) plots can be used for the purposes of convergence checks (Vehtari et al. 2021). It is customary to train Bayesian models on several independent runs (called chains). When there are multiple chains trained, $\hat{r}$ helps identify problems with convergence of densities, whether it be a single chain having difficulty converging or different chains converging to different posteriors. For a given parameter, if $\hat{r}$ < 1.01 or ($log_{10}\hat{r}$ < -2) is not satisfied, then it is indicative of such a problem. ESS plots show the effective number of samples for each variable after all the chains are combined. This is less than the total number of generated samples and intuitively shows the number of samples after the correlation between samples has been accounted for. This can be done in a way which partitions the sample space into quantile regions to assure there are enough samples to characterize different regions of all the posteriors. The accepted threshold is that the effective number of sample sizes should be larger than 400, although a higher number across different quantile regions of the sample space to make sure that the posteriors can produce reliable statistics. We set the threshold at 1000 to make sure tails of the distribution are characterized more efficiently. These diagnostics for the three experiments described in the main manuscript are shown in Fig SI. 7 below.

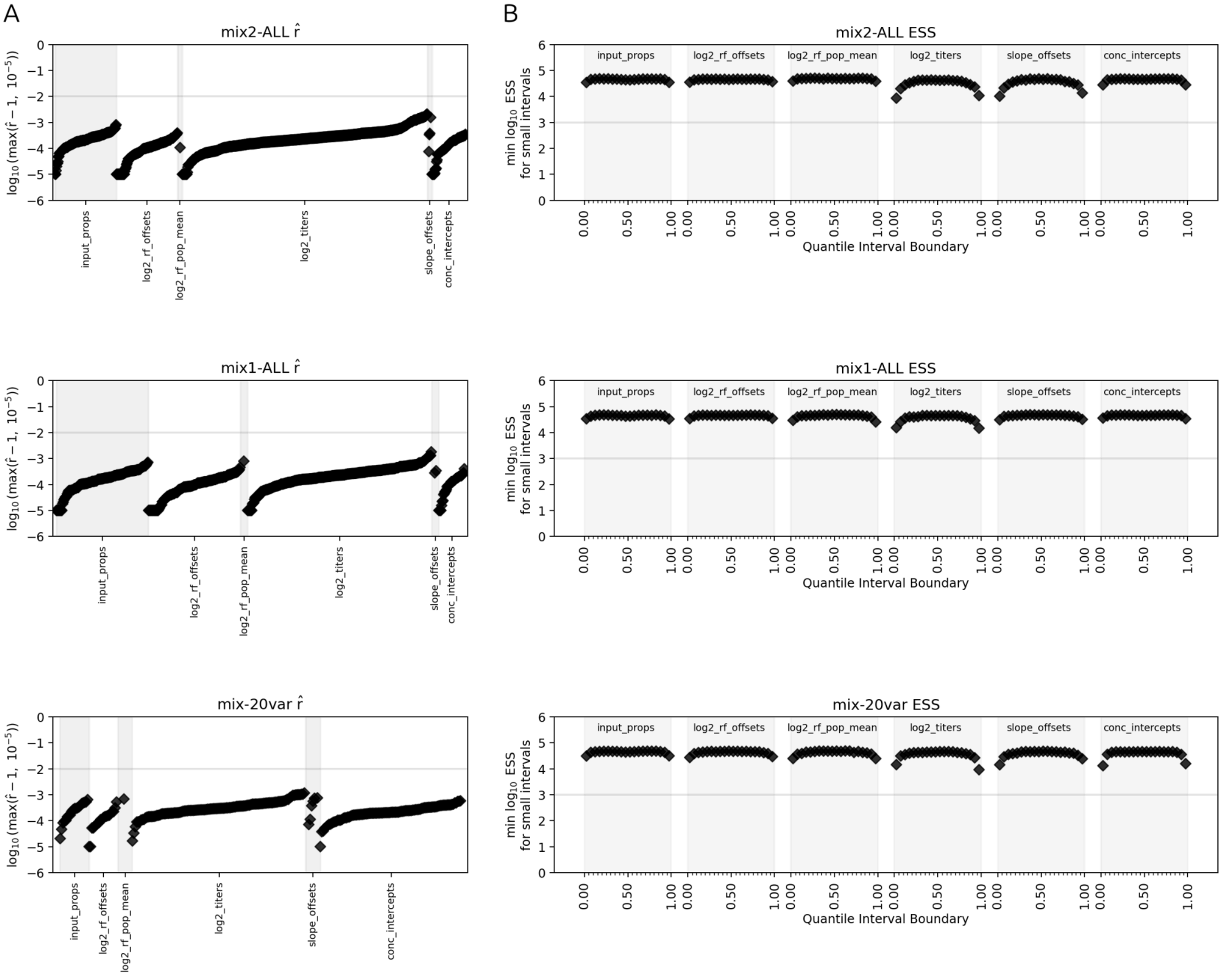


**Fig SI. 7 Model convergence diagnostics for datasets obtained with mix1-ALL, mix2-ALL and mix-20var.**

**(A)** $\hat{r}$ diagnostics for the three datasets coming from mix1-ALL, mix2-ALL, mix-20var. Horizontal grey line indicates the threshold $\log_{10}(0.01)$ beyond which posteriors likely display multimodality or not sufficient sampling. **(B)** ESS diagnostics for the same datasets. Horizontal greyline indicates the minimum required number of essential samples below which statistics (especially those involving tails) may be biased. Both the $\hat{r}$ and ESS calculations are done with ArviZ (Kumar et al. 2019) where for the latter we use method="local" and a quantile range of 0.05 to 0.95 with 20 quantile points placed uniformly in this interval including boundaries. After calculating interval ESS values for every parameter in the model, the minimum at each quantile interval is plotted. $\hat{r}$ and ESS values are shown for every variable in the model grouped according to parameter name. We run these tests on 10 chains each with length 5000.

#### 2.2 Fit and Cross-validation Diagnostics

We then carry on with model predictive accuracy tests which can be categorized into goodness of fit and cross-validation using proportions of variants as the test observable (see section Model Overview for descriptions of these).

For posterior predictive checks we show the plots in log scale for both x and y axis which facilitates the visual evaluation of the fit. This means that we cannot include zero counts in the plot so we write the proportion of zeros for the posterior samples and observed data in the title of the plots. For comparison of residuals, we compute the difference between observed log proportions and sampled log proportions and plot this as a histogram. We compare it to the histogram of differences between observed log proportion repeats. For cross validation we use LOO-PIT. Vanilla LOO-PIT is only suitable for continuous data. Arviz allows discrete data via randomized tie-braking that occurs for CDF of discrete distributions and therefore we use this approach to compute the LOO-PIT of our count data.

In this section we refer to experiments in the main manuscript carried out with mix1-ALL, mix2-ALL and mix-20var as experiments 1, 2 and 3. The former two have 134 strains and respectively four and two sera whereas the latter have twenty variants and six sera.

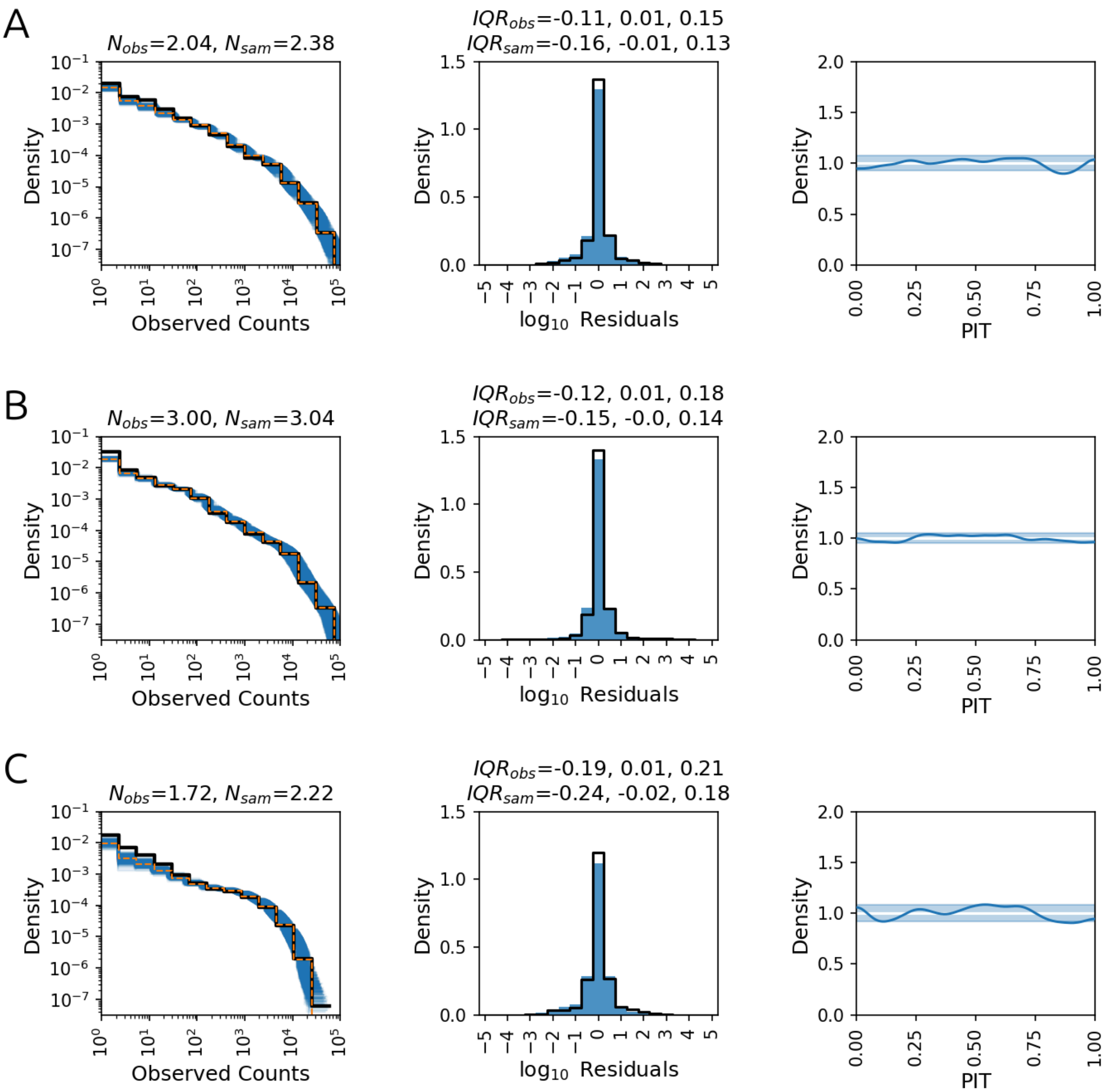


**Fig SI. 8 Goodness of fit and cross validation checks for datasets obtained with mix1-ALL, mix2-ALL and mix-20var.**

Panels A,B,C show PPC (first column), Residuals (second column) and LOO-PIT (third column) for data respectively for mix1-ALL, mix2-ALL and mix-20var. The three comparisons are identical to those in Fig SI. 3.

There are several things to notice in Fig SI. 8. The first one is the slight underestimation seen in the PPC plots (first column) for counts around 10. We deal with sequence datasets with the number of sequences generally higher than 100K. We apply stringent quality checks (see Sequence Processing section in SI). However with so many sequences, there will be inevitable noise signals coming from either to lesser extent sequencing errors and to a higher extent from biochemical sources like replication, transcription and PCR (see section Analysis of Error Rates section in SI). Because this noise is not included in the model, these very low observed count variants are more often sampled as zero (rather than low counts) in the model. This is indeed what we observe from the numbers reported in the titles; the proportion of zeros in observed data is lower than the sampled ones. This may be tackled with a mixture model however mixture models are harder to sample and LOO-PITs reveal that there is no further need for complication. Trying such models did not result in any significant changes to titer estimates which have %94 HDIs lower than three and therefore suggests that these very low counts don't affect titer estimates significantly (not shown here).

The second one is that for the residual vs repeat error plots in the second column, the black distribution, despite corresponding with the residual errors quite well in terms of shape and IQR always seems to be slightly narrower than the blue one. This suggests that the posterior samples might be slightly over-dispersed. Over-dispersed posterior samples generally represent themselves as frown shaped curves in the LOO-PIT diagrams. We do indeed see that in the LOO-PIT diagrams, there is a slight frown-like shape towards the middle. This suggests that the model is slightly over-dispersed but the LOO-PIT plots suggest that this over-dispersion is not far beyond the acceptable limit for and LOO-PIT plots are very sensitive to smallest of deviations from ground truth.

#### 2.3 Comparison of Repeat Errors for mix1-ALL and mix2-ALL

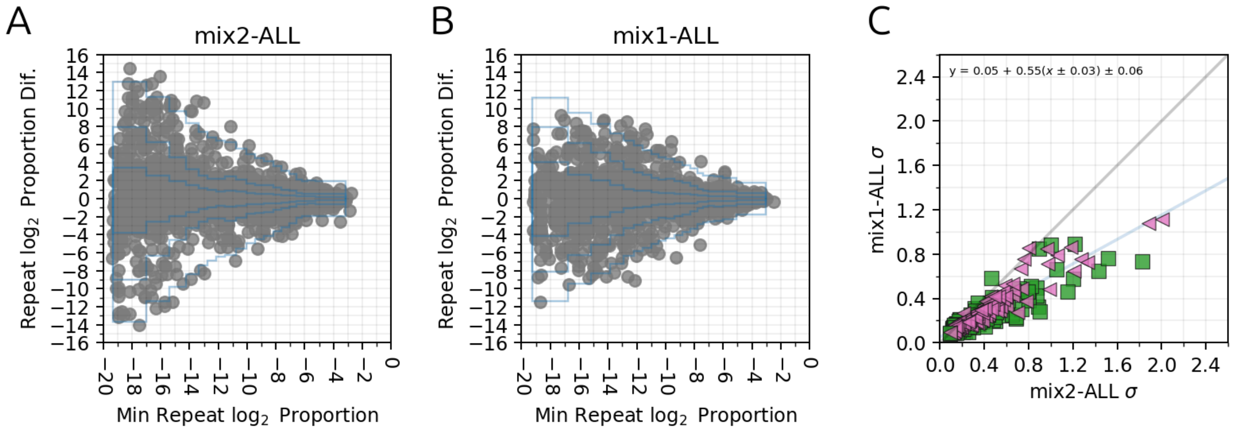


**Fig SI. 9 Comparing repeat error rates and parameter uncertainties for mix1-ALL and mix2-ALL.**

**(A,B)** are similar to the plots seen in the first row of Fig SI. 3. They show the proportion vs repeat error for the No Serum sample of respectively mix2-ALL and mix1-ALL. Construction of the blue quantiles lines is identical to that explained in Fig SI. 3. **(C)** Shows comparison of standard deviation of titer parameters estimates of individual viruses for the two common sera BR/8/96 (BE92, pink) and GD/25/93 (WU95, green).

The experimental changes implemented have positively impacted the results by lowering the amount of repeat noise (see Fig SI. 9) and smaller parameter standard deviations. We can see from the grey circles in the first two plots, as expected, that as x decreases the spread along the y-axis increases. However we see that especially for lower x values the spread in the y-axis is higher for the mix2-ALL experiment suggesting that the effect of stochasticity is more prominent in that experiment where rescue/passage is less optimized compared to mix1-ALL. Similarly, the amount of model fitted noise at lower x values (indicated by blue conditional quantile lines) is lower for the last quantile region in the second experiment. When low replicative fitness variants also have low PFUs they risk higher noise due to stochastic effects of sampling. The experimental changes were designed to make sure that such variants have a relatively uniform number of PFUs compared to other higher fitness variants. This is also reflected in the HDI values of titers, making them smaller for the latter optimized experiment. Note that in the second dataset, dilutions run until 1/10240 whereas in the first one until 1/5120. Given that both of the sera in the last plot of Fig SI. 9 are highly reactive sera, it is not surprising that adding a high dilution decreases the HDI significantly.

#### 2.4 Model Comparison

As promised in section 1.3 “Choice of Likelihood”, we demonstrate here that the variable concentration BetaBinomial model clearly overperforms (for cross-validation) the constant concentration BetaBinomial when tested on the full model. This is a necessary check to ensure that the former model which has more number of variables is not over-fitting. For this we use the 145 and all seven positions mixture from the third dataset in section “Further Optimization of the Rescue and Assay”. We run the assay using the model with constant and variable concentration (see SI Section The Full Model for description of models and SI Section Choice of Likelihood which explains these models in a simpler context).

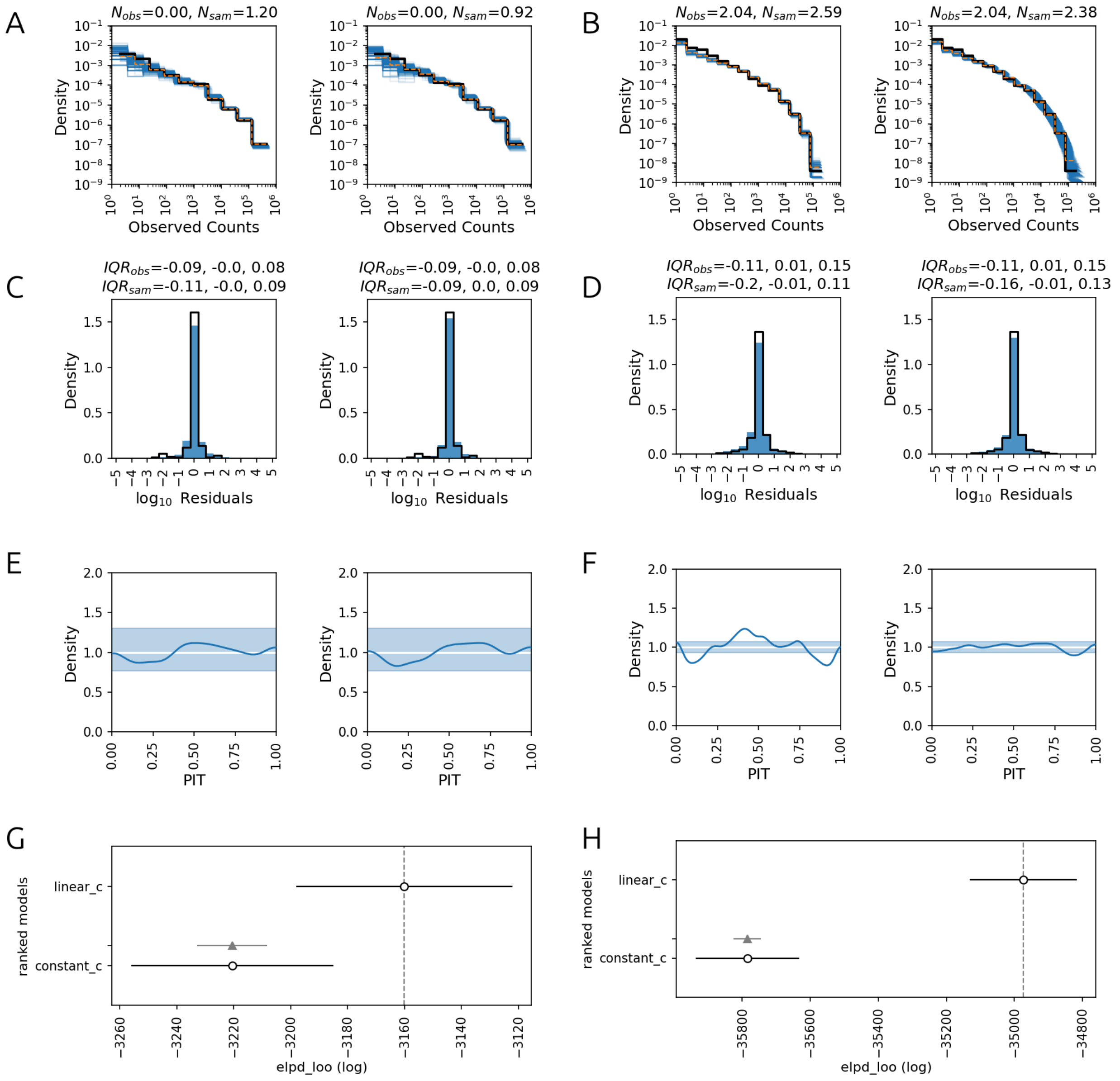


**Fig SI. 10 Comparison of Constant and Variable Concentration Models.**

**A,C,E,G** shows various diagnostics for the constant (first column) and linear (second column) concentration model fitted on mix1-posn145 (N=19) whereas **B,D,F,H** show the same diagnostics for mix1-ALL (N=134). The comparisons made in this figure's first three rows use the identical methods as Fig SI. 8. **(A,B)** shows goodness of fit between observed counts (black) and posterior samples (blue individual, orange combined), **(C,D)** compares log10 repeat errors vs log10 residuals and **(E,F)** checks cross-validation where the ideal case is a straight white line and the blue band shows %99HDI for amount of deviation from straight line with the same number of samples. **(G,H)** are model comparisons done with the ArviZ package compare and plot_compare functions. This method estimates the leave-one-out posterior distributions and compares the model predictions accuracy. The model which performs best is displayed on top and the bar shows the standard error of the expected log pointwise predictive density (ELPD) and the grey bar shows the standard deviation for the paired differences (where pairing is according to which data point is left out) (Vehtari et al. 2015). For a model to be significantly better, one would require the grey bar with triangle to not overlap with the dotted horizontal line.

We see from Fig SI. 10G, that the variable concentration model outperforms constant concentration for both lower number of variants (19) and higher number (134) but the difference is more significant for the latter case. In both cases the constant concentration model is over-dispersed as is evident from Fig SI. 10C,D (more diffuse blue histograms) of both panels and  Fig SI. 10F (a more significant upside down frown pattern in loo-pit). This suggests that the added number of parameters in the variable concentration model is not redundant.

#### 2.5 Prior Informativeness and Sensitivity Tests

In this section we test our claim that the priors we use are weakly informative and we also carry out a prior sensitivity analysis.

Although there is no precise definition of weakly informative prior, one simple perspective is that for a parameter of unit scale, if the prior has standard deviation of 10 then it is weakly informative and if the prior has standard deviation 1 then it is generic weakly informative (GitHub, n.d.). To assess whether a certain model fits any of these criteria, one can look at the standard deviation of posteriors for each parameter and compare it to standard deviations of its prior. If standard deviation shrinks about 1/10, then one can say that the prior choice was weakly informative.

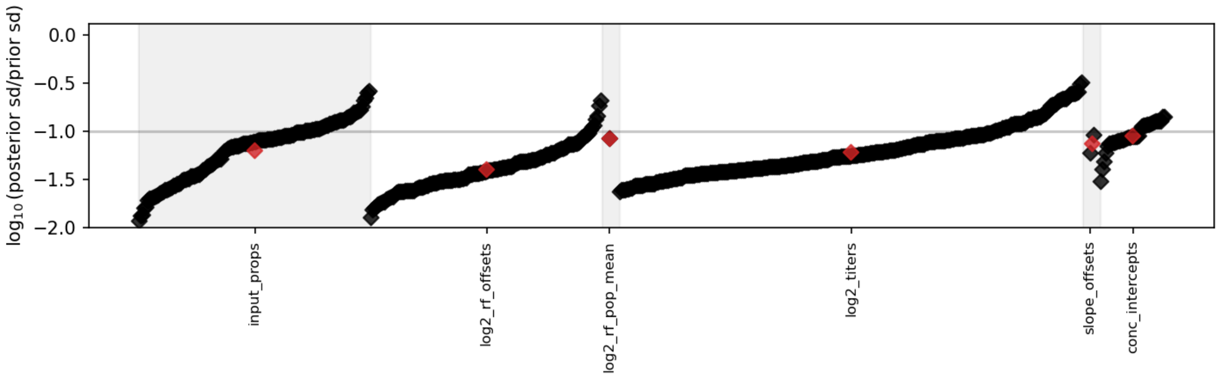


**Fig SI. 11 Prior Informativeness Test**

The plot is separated into groups of parameters that exist in the model and for each parameter the log10 ratio of posterior standard deviation to prior standard deviation is shown sorted from lowest to highest. Data is from mix2-ALL.

As can be seen in figure Fig SI. 11, most of the parameters in the system are below the 0.1 limit (-1 in log10) or close to it. All of the parameters that we are interested in such as titers and replicative fitness show shrinkage more than 3 fold and generally around 10 fold. Those that show less shrinkage are the ones with generally higher HDIs for titers and replicative fitness, i.e low count viruses with higher uncertainty and more noisy data. It is also therefore expected that parameters associated to them show higher posterior standard deviations. Nevertheless, the average value of shrinkage for both of these parameter sets is well below 10 fold.

In sensitivity analysis we modify the priors and prior parameters in various ways and gauge how much the posterior predictive distribution and the titer posteriors change.
To change the priors we follow the following recipe: 1- We change Normal priors with SkewNormals with α=3 and InverseGamma with Gamma or vice versa, 2- We also apply various rescalings to the standard deviation parameter of these priors. 3- Since SkewNormals can't be truncated in pymc, we replace the truncated normal distribution for titers with one where mean has an offset +1 (apart from the sd scaling). For the sd scaling we try scales of 0.25, 1 and 4. When scale=1 then the only change is the form of the priors. For instance, changing the Normal to SkewNormals with α=3 has the effect of adding a slight positive bias to the system and also reducing variation slightly.

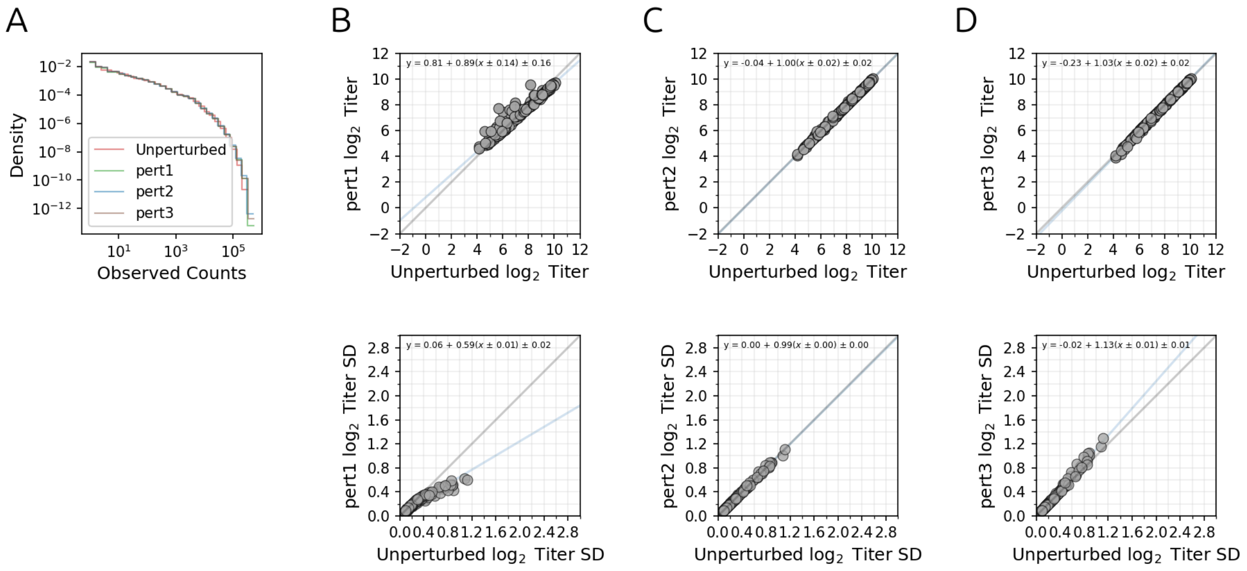


**Fig SI. 12 Sensitivity of Posteriors to Priors**

**(A)** The posterior predictive density histograms of the samples from the models with different priors given by the labels. Original is the model without any modifications, pert1 is where the priors are changed and standard deviation are rescaled by 0.25, pert2 is where the priors are changed but standard deviation is not rescaled, pert3 is where the priors are changed and standard deviations are scaled by 4. Bins and histograms for each of the posterior predictive distributions are determined individually using get_bins and histogram functions inside the stats.density_utils module of ArviZ package (Kumar et al. 2019). **(B-D)** Each column shows respectively plots for comparison of the original model (x-axis) with different perturbations (y-axis) discussed in (A). The first plot in each column shows the mean values for the log titer posteriors (axis ticks labelled in linear). The plots on the second row show the same for the standard deviations of the log titer posterior distributions. The grey line shows the diagonal and the blue line shows the orthogonal regression for the scatter plots. In the last plot seven outliers of comparison are not shown in the plot, see the manuscript SI for a discussion of these.

The sensitivity analysis in Fig SI. 12 reveals that the model results are robust against introducing skew to the model as well as changing standard deviations of the priors. It also however reveals that scaling the standard deviations by 0.25 might bring some of the priors to a too restrictive level given the number of data points and reduce the non-informativeness of the priors. This is evident from the first plot of the second row where the standard deviations for the first perturbation is reduced compared to the original model. On the other hand, most of the SD comparisons in the third perturbation (where standard deviations are scaled by 4) agree with the unperturbed model suggesting that the original model's priors are generally non-informative enough, more vagueness is not necessary. Therefore our choice of priors sit at the sweet scale of not too informative or uninformative for the titer estimates.

### 3. Extra Analysis

#### 3.1 Analysis of Error Rates

In this section we present an estimate of the combined error rates due to processes such as replication, transcription, sequencing. Note that in an ideal situation of no error, one would only expect to encounter variants which are created at the seven positions by directed mutagenesis. Any other variants encountered could be expected to come from biochemical errors. To estimate error rates, we focus on a subset of positions where deviation from root virus' sequence is expected to occur in later stages of the experiment (replication during assay, transcription of RNA to DNA and PCR) rather than earlier stages such as mutagenesis and rescue. We exclude the positions that were subjected to target mutagenesis, and any nucleotide that is within six bases difference to these positions (we have observed higher error rates generally around these regions suggesting they might be affected from the plasmid transformation step). We also exclude about nine nucleotides from the beginning and end of the inserts as these regions are more liable to clipping and alignment errors. For any other position we calculate the rate of errors in the sequences using the data obtained with mix1-ALL and mix2-ALL. The error rates are calculated in the form of nucleotide transitions or transversion for single nucleotide differences. Double nucleotide changes are grouped into categories tt, vv, tv, vv where t is transition and v is transversion. tt for instance means that the change was a double transition. Double nucleotide changes are further categorized into adjacent and non-adjacent. Finally triple nucleotide changes are collected into a single group. Once the error rates are calculated, any substitution whose rate is higher than the 0.95 quantile of its base distribution is used in naming the read variants and therefore not included in mean rate calculations for error rates.

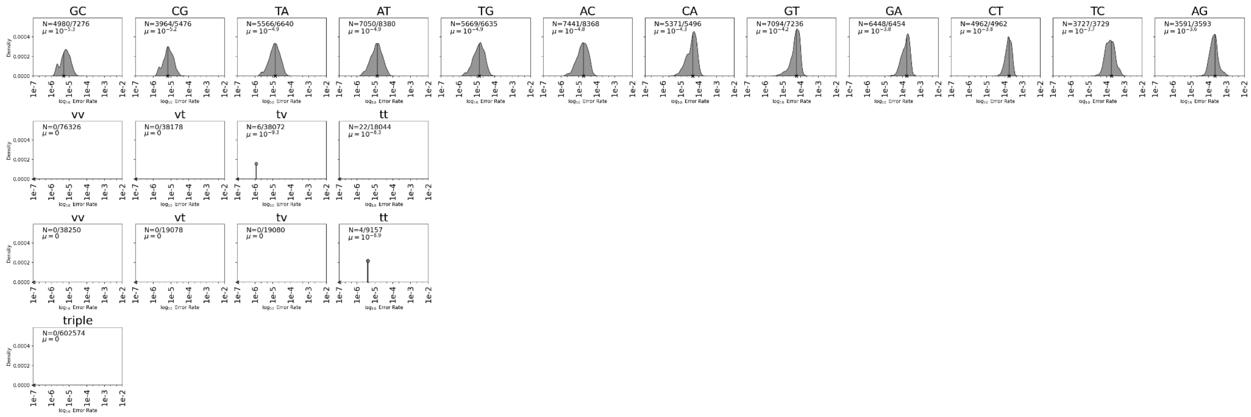


**Fig SI. 13 Log 10 Error Rates**

These figures show the log10 error rates computed using a subset of positions in the reference genome from the mix1-ALL. Text in each panel shows the number of sites across all experiments where non-zero rate was observed and the log10 mean rate (also counting the zero rates). **(A)** Single nucleotide errors, **(B)** adjacent double nucleotide errors, **(C)** non-adjacent double nucleotide errors and **(D)** triple nucleotide errors. Double nucleotide errors are categorized into whether it is a transition (t) or transversion whereas triple nucleotide errors are collected into one single group. If there are less than three unique non-zero rates, rates are shown as a bar with a circle marker at the top for each observation. The height of the bar is the number of times that rate is observed divided by the total number of observations including the zero rates. Otherwise a gaussian kernel is fitted to the densities.The x marker at the bottom shows the log10 of the mean of the rates (which also includes zero rates in the kernel density estimates). The densities are scaled by the ratio of non-zero rates so that different transitions can be compared on an equal footing.

The error rates computed here align with biochemical expectations: transitions generally have higher rates than transversions, adjacent double transitions have higher rates than non-adjacent, triple has the least. Transition single nucleotide error rates are generally around a rate of 1e-4 to 1e-3. In a case where the root variant proportion is around 0.1 of the whole table sample, then this means a transition could affect a variant up to about 1e-5 to 1e-4. If the number of sequences are around the scale of 100K, this equates to roughly about 1 to 10 sequences. This might explain why the model posterior predictive samples have more difficulty producing the correct density around here (see Fig SI. 8). A mixed model as was done in the SI section “Further Diagnosis of Model Fits” can address this. However, as explained in that section, the disadvantages of such a complex model outweigh the advantages and therefore we have only applied it occasionally for testing purposes.

We also compare the rates from mix1-ALL to mix2-ALL as well as rates from other sources and methods.

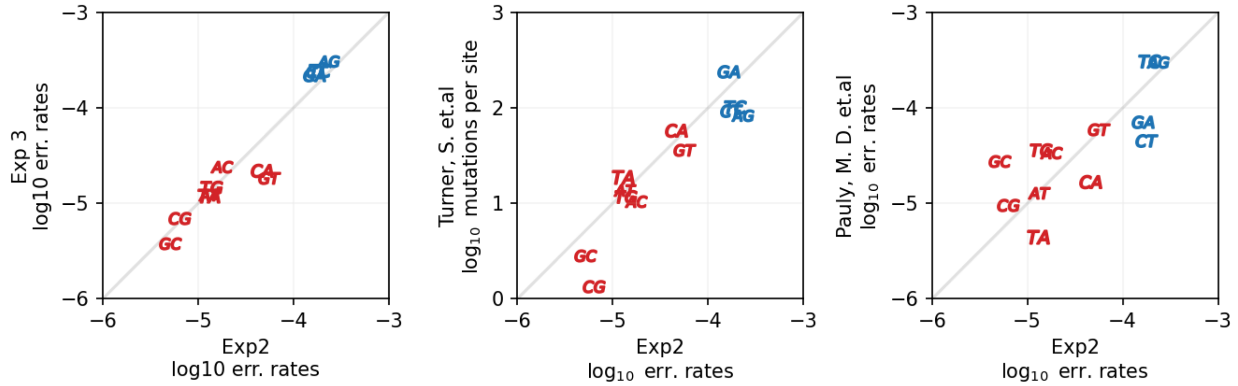


**Fig SI. 14 Comparison of Single Nucleotide Error Rates Computed Using Different Datasets or Methods**

**(A)** The single substitution error rates obtained with the method described in this section obtained using mix1-ALL and mix2-ALL. **(B)** mix2-ALL single substitution error rates compared to error rates from (Turner et al. 2026) which is a tree based mutation counting method that spans H3N2 influenza strains from XYZ to XYZ. **(C)** mix2-ALL single substitution error rates compared to (Pauly et al. 2017) which calculates the replication error rates on the root A/Hong Kong/4801/2014 H3N2.

In order to tease out the contribution of sequencing errors on the data, we compare the error rates when the quality threshold is 50 vs 10 (using the dataset from mix1-ALL). Note that this is the threshold for merged sequences whose qualities are computed from qualities of paired reads according to a Bayesian rule (Edgar and Flyvbjerg 2015). One can roughly think of it as addition of qualities for bases from the paired reads when bases agree and difference (max from min) when they don't. Our prior theoretical calculation of expected error rates (based on distribution of merged quality scores) suggests sequencing errors should be around 1e-5 - 1e-4 when Q=10 and less than 1e-6 when Q=50 (calculation not shown). The results in Figure Fig SI. 15 confirm this as when the quality threshold is dropped, only error rates which are around or less than 1e-5 - 1e-4 are affected and they generally increase to a mean rate of around 5e-5, whereas those which are around 1e-4 - 1e-3 don't change appreciably.

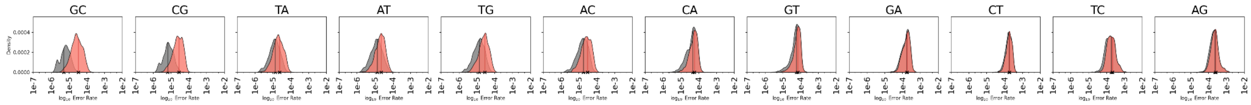


**Fig SI. 15 Comparison of Log10 Error Rates with Higher and Lower Quality Thresholds**

Error rates calculated as in Fig SI. 13 for the cases of when the filtering quality threshold is 50 (grey) and 10 (red). When the filtering threshold is 50, after the sequences are merged any base which has a lower quality than this threshold is not taken into account for error rate calculations.

### 3.2 Relative Titer and Replicative Fitness Line Plots with HDI Intervals

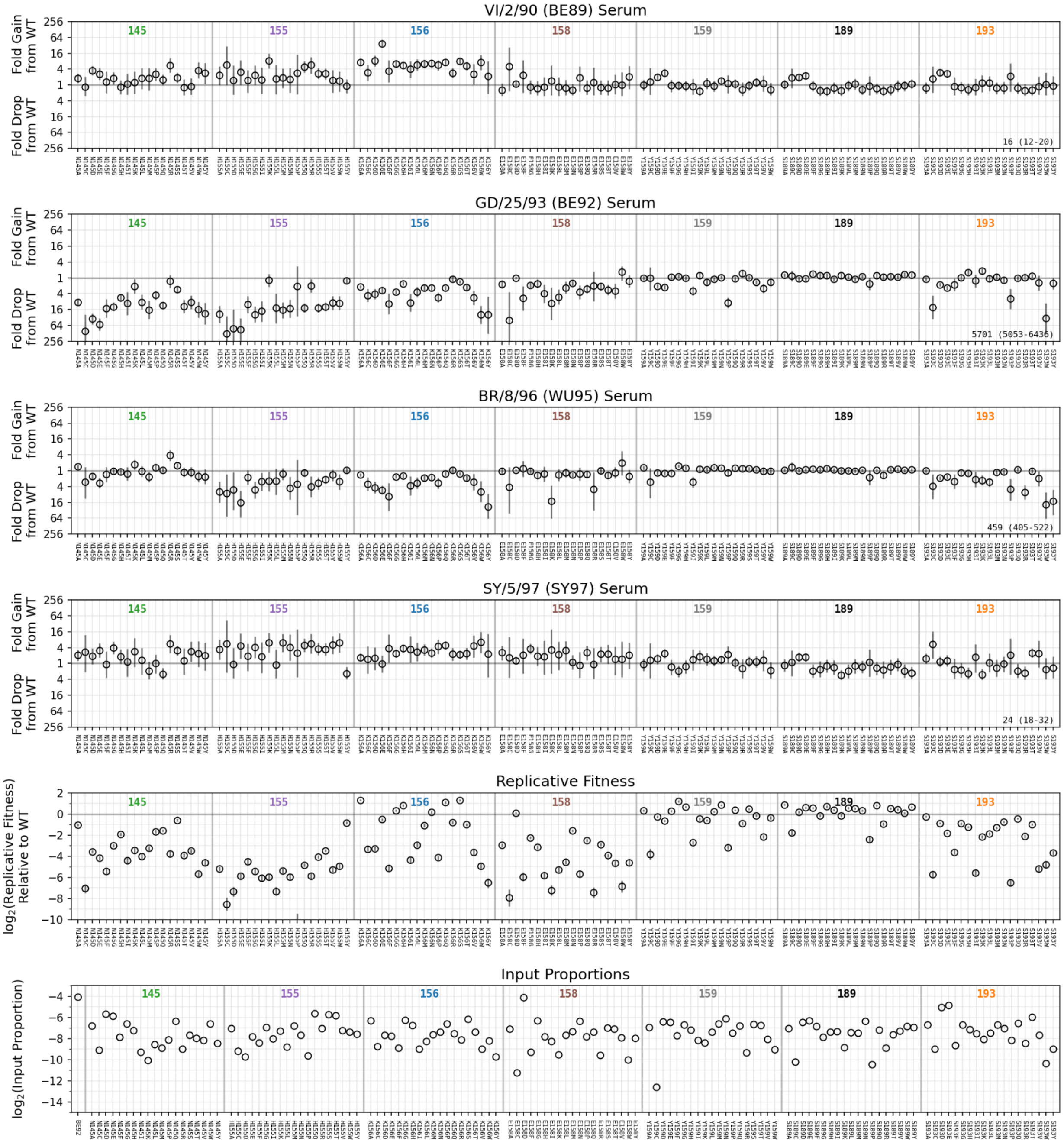


**Fig SI. 16 Various parameters of interest and observables obtained from mix2-ALL**

The first four plots show the mean titers relative to the root variant and the fifth one shows mean $\log_2$ replicative fitness relative to the root variant. The last plot shows observed $\log_2$ input proportions for m mix2-ALL. Errorbars show the %95 HDI intervals for the difference of observables. Horizontal lines show position divisions for the seven mutated positions: 145, 155, 156, 158, 159, 189, 193.

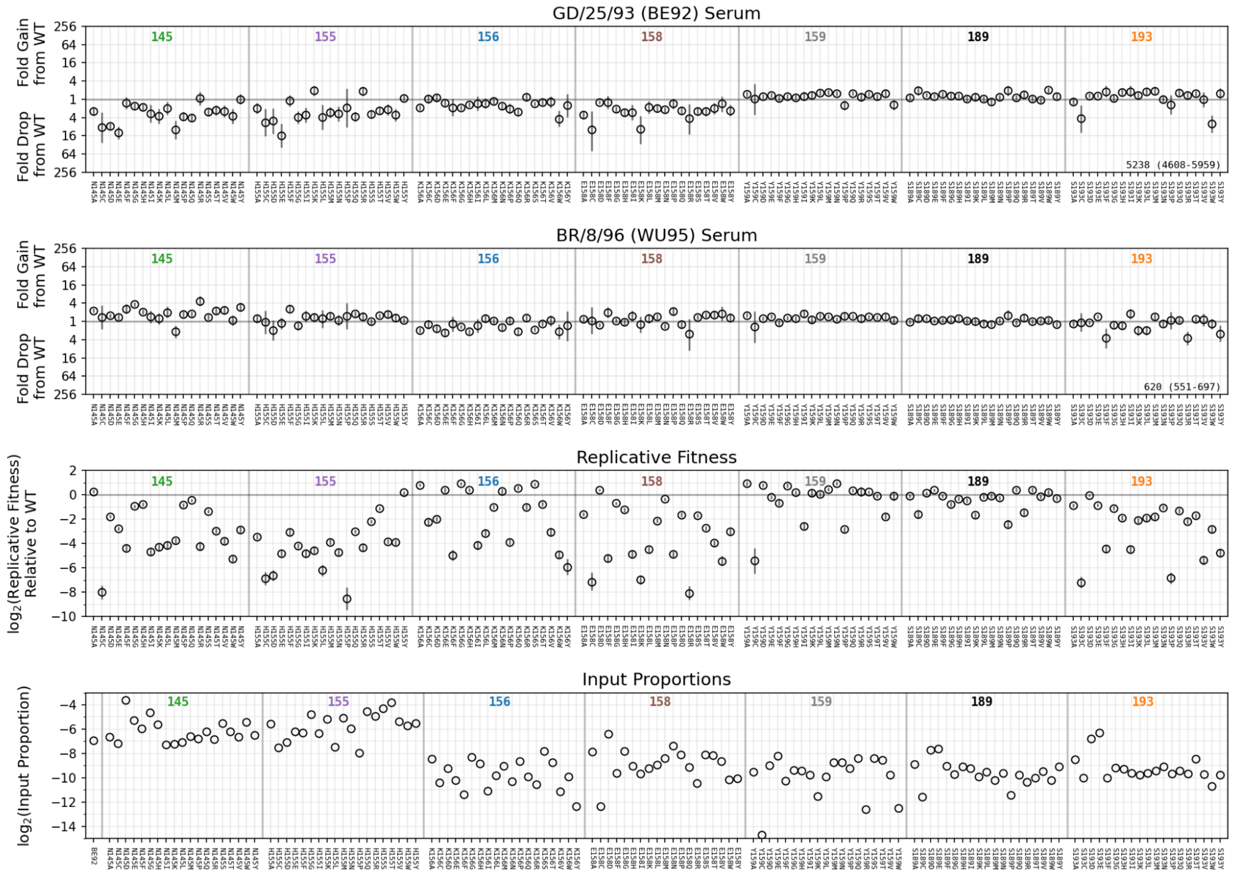


**Fig SI. 17 Various parameters of interest and observables obtained from mix1-ALL**

The figure is identical to Fig SI. 16 except that it shows data from mix2-ALL.

### 3.3 Fitted Curve Examples

In this section we show examples of fitted curves for a subset of strains using the data from mix1-ALL. Note that the models used in this paper fit to sequencing counts and not neutralisation data. Therefore we first show log10 sequencing proportion curves (numbers of sequences for each table sample are constant therefore can be disregarded when comparing fits). We use log10 so that strains of different proportions can be easily  compared visually.

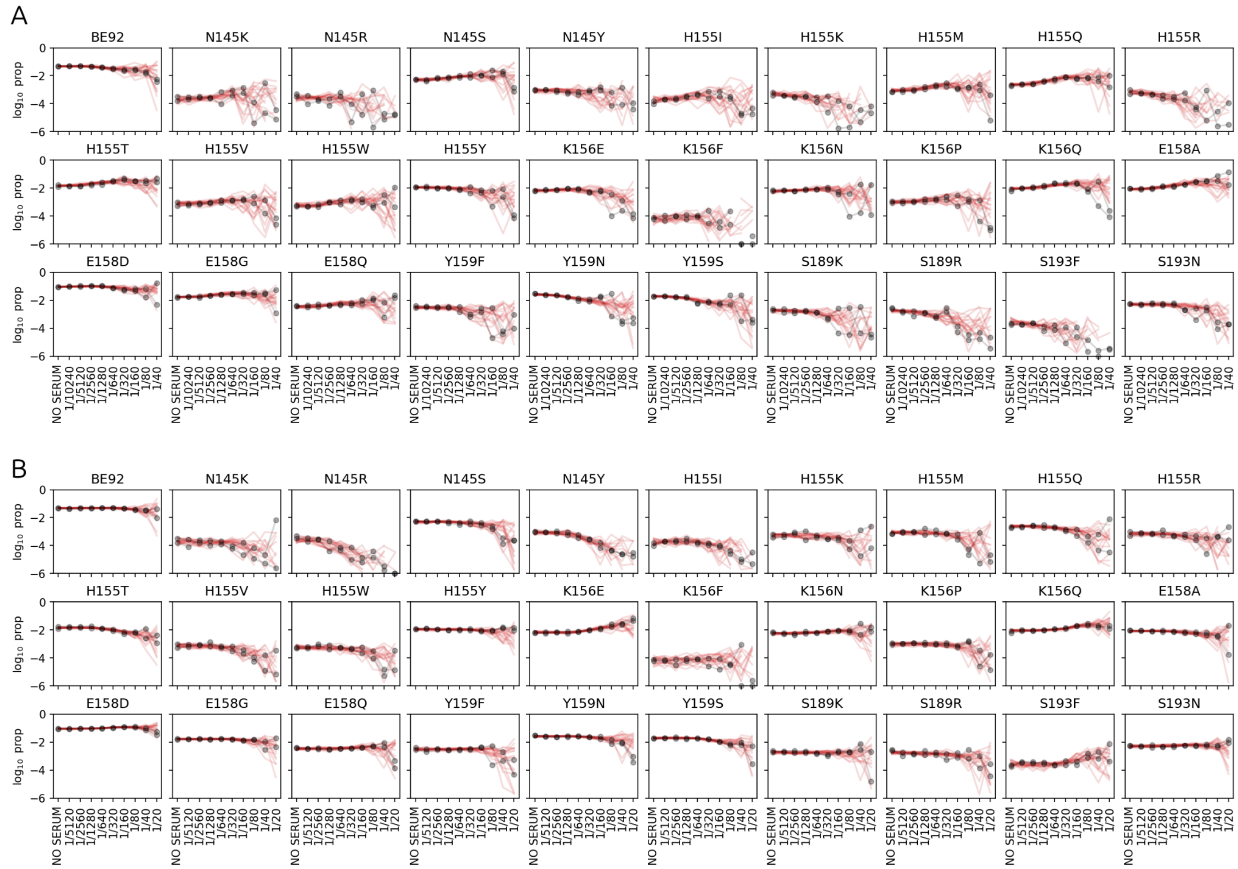


**Fig SI. 18 Sampled and observed count proportions**

**(A,B)** Sampled and observed proportion data respectively for WU95 and BE92. The grey markers show observed count proportion repeats (N=2) for either no serum or the given dilution in the x-axis.The red lines are twenty sampled posterior predictive count proportions. Zero proportions are not shown in the plots.

Fig SI. 18 demonstrates the utility of the Bayesian mindset. Instead of succumbing to the practice of fitting a curve to each of the repeats, one constructs a simulator for the experiment that can produce a whole range of behaviours which encompass the observed repeats. This simulator can then be used to analyse different scenarios such as different numbers of strains, lower or higher number of sequences, lower or higher reactivity etc (see SI Section “Relation Between Noise, PFU and Average Neutralisation”). This in turn can inform decisions about further pushing the capacity of the experiment pitting it against the precision required.

Even though the model does not fit neutralisation curves they appear as intermediate transformed latent variables (see equation 6) and therefore can be sampled. Moreover one can take the sequencing counts and back calculate latent fraction remaining for each virus at each dilution. The details for how this is done can be found in the Thor package samplers module function BB_sample_neut. If the model is working as it should, one could expect the latent 1 - neutralisation curve to agree with the back calculated fraction remaining.

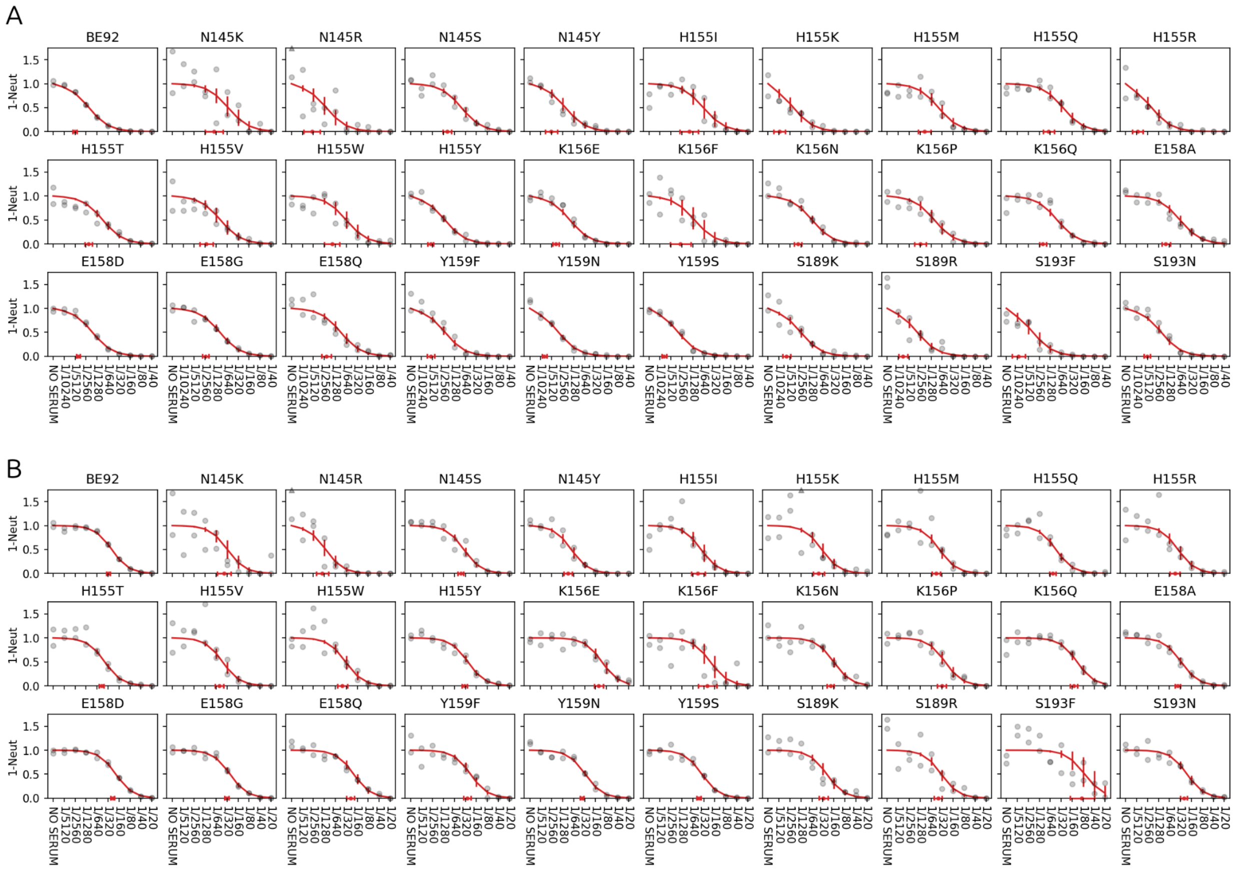

**Fig SI. 19 Backcalculated latent neutralisation data and fraction remaining**

The figure is similar to Fig SI. 18 except that it shows neutralisation curves instead of sequencing proportions. The error bars show %95 HDI for the neutralisation curves. Note that the full extent of the noise in the observable is captured not only by the variation in the neutralisation curves but also by the noise in the BetaBinomial likelihood that appears in the model. Moreover the grey dots are no longer strictly observed quantities (since the only observed quantities of the experiment are reads and ct values). Instead they are constructed by combining observed read proportions with some of the parameters sampled from the system (such as replicative fitness and quantities representing the amount of viruses in the system relative to no serum). They represent latent fractions of remaining viruses at each dilution. Since these will themselves be distributions, their mean value is plotted as the grey dots.

### 3.4 Relation Between Noise, PFU and Average neutralisation

In order to build a simulator which can study the effects of changing parameters such as PFUs, number of strains and neutralisation we need to understand how the concentration parameters depend on these. Note that concentration parameters in the model are structured independently, i.e they do not have an explicit dependence on any of the mentioned values. If the hypothesis that lower concentration (i.e higher noise) is mostly caused by the sampling bottleneck after incubation then there should be a relationship between the concentration parameter and

$$S^{r} = \frac{1}{N_{v}} \times \frac{PFU}{N_{v}} \sum_{n=1}^{N_{v}} n_{i}^{r}$$

where **r** stands for the table sample index, $\mathbf{N_v}$ is the total number of variants, $\mathbf{n^r_i}$ is the value of neutralisation curve for virus **i** at table sample **r**. Note that the second factor in the right hand side represents the expected number of PFUs left in the table sample after neutralisation and therefore dividing it by $N_v$ gives the expected number of PFUs per variant left after neutralisation. We analyse whether if this is related to the parameter ra (see section "Concentration Prior" in SI, equation 7) and the slope of the concentration line, which is (la-ra)/$|p_{min}|$ where $p_{min}$ = floor(min(log10( proportions))). Note that the higher the S, the more PFUs per variant and therefore one would expect less noise.

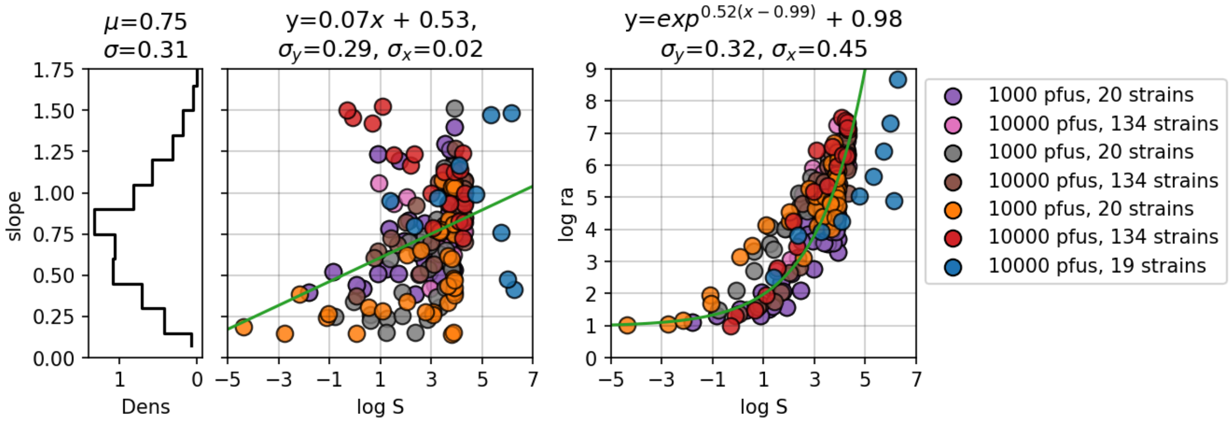


**Fig SI. 20 Relation Between logS, ra and slope**

The first two plots show respectively the marginal density of slope and scatter plot of logS vs slope. The title on the first plot shows mean and standard deviation and the titles on the second plot show the slope and intercept parameters of the three green quantile regression lines for q=[0.05, 0.5, 0.95]. The last plot shows the scatter plot of logS vs log ra (right asymptote for concentration, see SI section Concentration Prior). The green line in that plot shows the exponential regression with the parameters and residual standard deviation shown in the title. The dots in the scatter plots are colored according to table samples whose PFUs and number of strains are shown in the legend.

The analysis in figure Fig SI. 20 reveals a strong relationship between log ra and log S regardless of dataset. On the other hand, the relation between slope and log S is not so strong (even when the x axis is in log scale and y in linear), the general trend looks random though some datasets (i.e red) show more of a linear relation. In simulating an NGS-RFA/FRA system, one will be inputting titers, replicative fitness and PFUs to the simulator. Titer curve slopes and pfu scales for titers can be generated using the posteriors in one of the experimental datasets (one would not expect them to depend on other amounts of neutralisation though they could depend on the experimental setup). Finally, neutralisation values (to be computed from titers and slopes) and PFU can be used to generate la and slope by

$$log\, ra^{r} = exp(0.53(logS^{r} - 1.09)) + 1.02 + \epsilon_{1}$$

$$s^{r} = 0.07 log S^{r} + 0.54 + \epsilon_{2}$$

where $\epsilon_1$ and $\epsilon_2$ are normally distributed with mean 0 and standard deviations 0.89, 0.27. The slope for the concentration curve can also be determined by s ~ Normal(0.75, 0.31) given the weakness of the relation. Once a number of sequences per table sample is determined, then the likelihood of the full model can be used to simulate repeats for novel conditions. The simulated repeats can be used as observations for fitting the model to see how much changing the parameters affect the results. A simulator module is available in the Thor package to carry out such a simulation for the interested reader.

We have carried out one such simulation in which we try different numbers of variants and PFUs. In order to simulate log2 titers and log2 rf values, we fit kernel density estimates (KDE) to the log2 titer and log2 rf values fitted to the dataset presented in the section “All Single Substitution Variants at the Koel Seven Positions”. After building the KDEs we sample from there for the given number of variants. After these values are sampled, the simulator module in the Thor library is used to simulate data and then a model is fitted to the simulated data. The residual errors for the fits and HDI widths are compared in Fig SI. 21.

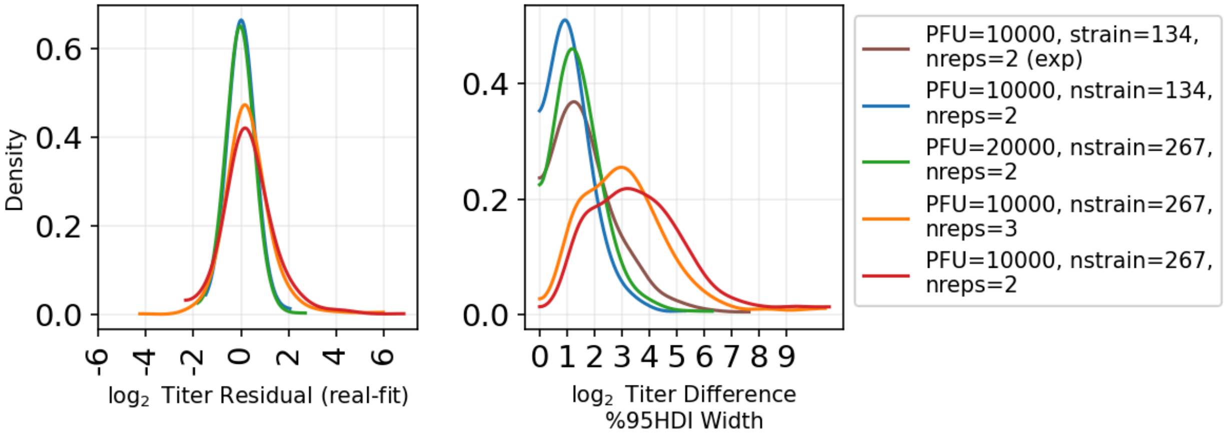


**Fig SI. 21 Residuals and HDIs Fitted for Simulated Datasets**

FIrst plot shows a kernel density estimate of the distribution of residuals of log2 titers for each of the simulated datasets where real titers are known. The second plot shows the kernel density estimate of the distribution of %95 HDI widths of log2 titer differences from wild-type both for experimental (brown) and simulated datasets (x-axis plotted in log scale). The number of PFUs, variants and repeats are given in the legend. Kernel density estimates are plotted via the plot_kde function of the ArviZ library with bw=0.5.

From the analysis of simulated data, we see that a combination of 10000 PFUs with about 134 strains is likely near the sweet spot. Indeed the blue and brown HDI curves seem to peak around an HDI width of around 1 suggesting that with a %95HDI, differences more than 2-fold can be significantly identified under this setting, which is generally a sufficient amount of power to determine important antigenic changes on a single substitution level. On the other hand lower PFU per variant curves peak around values of HDI around 3 suggesting only differences higher than 8-fold can be reliably identified when one reduces the number of PFUs per variant to such low levels. An 8-fold level of power can only generally identify antigenic differences between representative variants from different clusters that differ in multiple important antigenic sites.

#### 3.5 Fitting the Model to Data with Single Repeats

The analysis presented in the last section allows us to fit the model to experimental data with single repeats. We call this a parametric concentration model. In the previous cases the concentration parameters were estimated from the dispersion observed between repeats. In the parametric case we use the exponential relation revealed in the analysis done in the last section. In this model the priors for ra and la (see section “Concentration prior”) are given as

$$ra^r \sim N(0.65 + e^{0.39\, logS^r}, 0.9)$$
$$la^r \sim N(ra^r + 0.78|p_{min}|, 0.9)$$

where $|p_{min}|$ is the absolute value of the floor of the log2 observed proportions, superscript **r** indicates table sample index and $\mathbf{S}^r$ is defined in the previous section. We train this model on the dataset of the section “Further Optimization of the Rescue and Assay” and compare the estimated titers and HDI values to the model trained on two repeats.

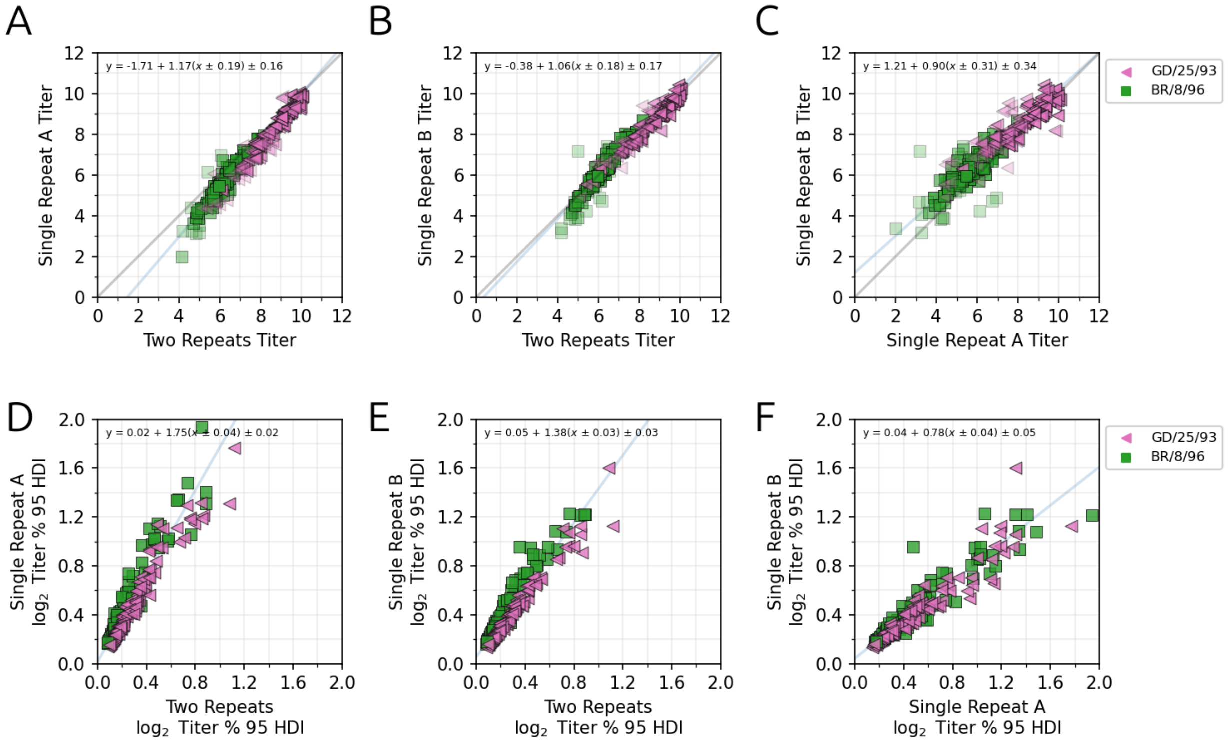

**Fig SI. 22 Comparison of Titers and HDI Widths Obtained from Data With Single and Double Repeats**

**(A)** Comparison of titers obtained by fitting the model on single repeat A vs two repeats A,B. **(B)** Comparison of titers obtained by fitting the model on single repeat B vs two repeats A,B. **(C)** Comparison of titers obtained by fitting the model on two single repeats A,B individually. **(D-F)** Same as data as in **(A-C)** except comparing %95HDI intervals for the estimates

We see from Fig SI. 22 that a model can, when needed, be trained on a single repeat dataset where the parameter estimates for the mean will be similar to that obtained from two repeats. The HDI widths of single parameter models will be roughly two times that of double repeat models, resulting in more HDI widths being larger than 1 (in log2 units).

#### 3.5 Pilot NGS-FRA/RFA with Two Time Points

An initial test was performed using a single serum A/Guangdong/25/93 (GD/25/93 in the BE92 antigenic cluster) and the twenty variant mixture previously described. Titrations were performed for dilutions ranging from 1/20 to 1/2560 at two growth time points (T=30h and T=40h) in duplicate. Comparison of T=30h and T=40h NGS-FRA titers confirmed assay stability across this window: orthogonal regression slope 0.89, standard deviation 0.27 (Fig SI. 23B). Comparison of T=30h NGS-FRA titers to HI titers gave slope 0.76, standard deviation 0.76 (Fig SI. 23D). One variant, H155R, showed systematically higher titers in the NGS-FRA than in the HI assay across all experiments. To determine whether this reflects a genuine assay-type difference rather than a pipeline artefact, we titrated the same virus using a classical plaque reduction assay and observed the same elevated titer relative to HI (Fig SI. 23C). This confirms that the discrepancy is a property of HI vs neutralisation-type assays for this variant, not a feature of the NGS-based method.

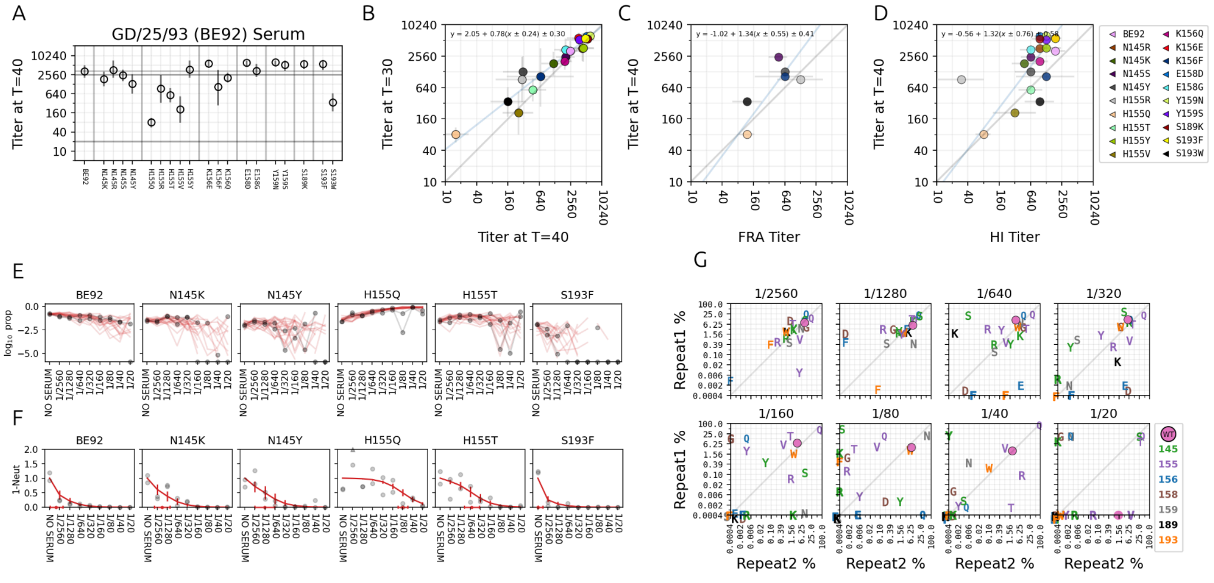


**Fig SI. 23 Testing the NGS-FRA/RFA Pipeline with a Single Serum (GD/25/93)**

**A)** The line plot for titers measured at T=40h as well as their %95HDI bars. The variants are grouped according to position using vertical separators. Horizontal lines show experimental dilution limits. The model titer range is +-1.5 $\log_2$ units of respectively upper limit and lower limit of detections for the experiment. **(B)** Comparison of NGS-FRA titers measured from supernatants collected at T=30h and T=40h. The blue diagonal line shows the orthogonal regression line and its equation is displayed as inset. The lines on each marker indicate the %95 HDI for the titer estimate for both cases.**(C)** Comparison of FRA titers vs NGS-FRA titers. **(D)** Comparison of HI titers vs NGS-FRA titers. **(E)** Comparison of observed repeat variant %s (grey markers) vs twenty posterior predictions sampled from the Bayesian model (red lines) for six exemplary viruses with variable behaviour. Zero observed proportions are plotted on the x-axis whereas zero sampled proportions are plotted as broken lines. x-axis shows dilutions on $\log_2$ scale, and NO SERUM at one minus the maximum dilution. **(F)** Comparison of latent samples for remaining fraction of six viruses (grey markers) to neutralisation curves (red) constructed from mean estimates for titer curve parameters. Markers outside the bound of the plot limits are shown with triangle markers at the top. x-axis is the same as the x-axis of panel E. Red markers and bars on the x-axis indicate mean estimate and %95HDI for the titers. In case $\log_2$(titer/10) < -1, then it is plotted at the NO SERUM x-level. **(G)** Comparison of repeat variant %s across different dilutions (given in the title). Marker color indicates position and letter indicates variant as given by the legend. Round marker is the root variant.

#### 3.6 Antigenic Cartography

To find a representative set of sera giving a similar triangulation as in, we identified a subset of ten first-exposure ferret sera such that when the full map is subsetted to these ten sera and antigens from clusters BE89, BE92, WU95, SY97 and remade by using the relax function in Racmacs ("Racmacs," n.d.), the topology of the map remains similar to the full map. triangulate the nearby clusters well, producing a smaller map, similar in topology to the full map (compare Fig.6A vs Fig.6B).

Map in Fig.9B and the map for reproducing the antigenic distances in Fig.9C were made using Racmacs with parameters minimum column basis of=320, dimensions=2, number_of_optimizations=20000 and default parameters for others. In Fig.9B the serum SY97 was not included in the map because there is no antigen that could be considered homologous-like for this particular serum and its titers are overall very low and non-discriminatory which would result in low angular precision.

#### 3.7 Counting GISAID Sequences

To produce the natural frequency data used in Fig.8B GISAID database sequences between 1979-01-01 and 1997-12-31 were considered. Among those sequences with more than %10 unidentified amino acids (i.e X) were not included in the analysis. The resulting sequences were aligned with maffts (Katoh et al. 2019) using default parameters. If after aligning more than %99 of the sequences have a gap at a position, sequences without gaps at that position are removed and remaining sequences are realigned. After alignment sequences are named based on their amino acids at the Koel Seven positions: 145, 155, 156, 158, 159, 189, 193 and then frequency of each named variant is computed.

### 4. DESCRIPTION OF THE IN-HOUSE CODE PACKAGES

For this pipeline we have developed three in-house packages. The first two packages Mer|ign (**Mer**ge **|** Al**ign**) (written in C) and Mitril (**M**er|ign **It**e**r**at**i**on **L**ibrary) (written in Python) are used to merge paired end reads and align merged reads against a reference. The third package Thor (**T**iter from **H**igh-throughput **O**utput **R**eads) (written in Python) is used to analyse the variants counts using the Bayesian models described in the SI section 1.4 “The Full Model”. In this section we give a brief overview of these packages.

**Mer|ign (Merge and Align)**

Mer|ign is a C library that consists of a pipeline with two components **mer** (for merge) and **ign** (for align) (Tureli 2025a). The output of the mer component can be directly piped into ign (hence the name Mer|ign) for a streamlined analysis without producing intermediary outputs (although using them as isolated components with file inputs/outputs is also possible). mer first aligns paired reads using Wavefront Aligner 2 (WFA2) (Marco-Sola et al. 2021, 2023) and merges paired reads according to the Bayesian update algorithm described in (Edgar and Flyvbjerg 2015). When merging paired reads, the quality of agreeing bases increases whereas those of which disagree decrease. We use a quality threshold of 40 which means that when the merged base has quality less than 40, it is recorded as a small letter to be handled separately in downstream analysis. When merging, regions of any of the pairs which are out of the domain of the overlapping alignment are discarded. This effectively trims the adapters because our libraries are optimised such that the insert, MID, and primer together is less than 300 base pairs which is the read length, so each pair comes with adapters on 3’ or 5’ ends that stick out of the pair alignment.

The second step, ign, aligns the merged sequence to a set of given references (using WFA2) and if MIDs are present identifies them. MIDs are aligned using Edlib (Šošić and Šikić 2017). Alignment of MIDs are done after merging and a merged read is only accepted as valid if MIDs at both the 3’ and 5’ are identified up to 1 substitution or indel and are identical at both ends (different MIDs at 3 ’ and 5’ ends are indicative of cross-over artifacts during index PCR). Any read whose alignment has more than dif_threshold (default: 50) number of mismatches is recorded as OTHER.

For more details on how the package works and its input parameters, see the package repository (Tureli 2025a). A bash script is provided in the Mer|ign package that builds this pipeline. The default parameters in the bash script are used throughout the paper except the merge quality threshold is set to 40, end_free and begin_free parameters are set to 100.

**Mitril (Merlign Iteration Library)**

This is a python package which contains functions to iterate through the outputs produced by Mer|ign to call variants (Tureli 2025b). A setting file must be supplied as input. We used the example setting file provided in the repository. Mitril carries out quality filtering, calculating error rates and calling variants based on a combination of presupplied positions and substitutions which are significantly higher than their base error rates (which we collectively call significant substitutions). In terms of naming variants, the Koel seven positions 145, 155, 156, 158, 159,

189, 193 are supplied as imporant_nt_positions in the Mitril settings file which means that any substitution that occurs in these sites will be used in naming the reads. The error rates (compound error rates of the whole process such as PCR, replication etc and not just sequencing) are calculated at a set of positions outside these regions (see the package repo for details) to get mean estimates for base error rates such as G→C, C→G etc and then a substitution at a position is used for naming if it occurs significantly higher than a given quantile (default: 0.95). This is a middle-ground approach to using every substitution that occurs in the alignments for naming (which could lead to significant reduction in data) and using only the important_nt_positions in naming (which risk incorporating context effects from substitutions which may be biologically significant).

In terms of quality filtering, any merged read which has a total error rate of >0.5 is filtered out. Moreover, any read which has a merged quality lower than 40 in one of the important_nt_positions is not used in counting. Finally any read which has more than 3nt insertions or 3nt deletions is also discarded. For those not discarded, then indels which are not multiples of three are recorded as X and those which occur in a codon window are either recorded as for instance Ins134C for an insertion or D134Δ for deletions. Note that variants with indels are not used in the analysis presented in this paper.

**Thor (Titers from High-Output Reads)**

This is a python library (Tureli 2025d) that contains the models described in the SI Section 1.4 “The Full Model”. The application of these models to the sequencing data obtained from these experiments is presented in this manuscript’s code repo (Tureli 2025c). Besides the models, it also contains functions for computing non-parametric estimates of replicative fitness and titer values, where the latter is computed via a combination of the Spearman-Kärber method and the iterative normalization method of Ayer-Brunk-Ewing Reid-Silverman (Ayer et al. 1955). See SI Section 1.4 “The Full Model” for more details.

**Main Code Repo**

The analysis presented in this paper is collected in a github repo (Tureli 2025c) .

### REFERENCES


Ayer, Miriam, H. D. Brunk, G. M. Ewing, W. T. Reid, and Edward Silverman. 1955. “An Empirical Distribution Function for Sampling with Incomplete Information.” *The Annals of Mathematical Statistics* 26 (4): 641–647.

“Bioicons - High Quality Science Illustrations.” n.d. Accessed July 14, 2025. https://bioicons.com/.

Bloom, Jesse D., and Richard A. Neher. 2023. “Fitness Effects of Mutations to SARS-CoV-2 Proteins.” *Virus Evolution* 9 (2): vead055.

Boni, Maciej F. 2008. “Vaccination and Antigenic Drift in Influenza.” *Vaccine* 26 Suppl 3 (Suppl 3): C8–14.

Czado, Claudia, Tilmann Gneiting, and Leonhard Held. 2009. “Predictive Model Assessment for Count Data.” *Biometrics* 65 (4): 1254–1261.

Dadonaite, Bernadeta, Jenny J. Ahn, Jordan T. Ort, et al. 2024. “Deep Mutational Scanning of H5 Hemagglutinin to Inform Influenza Virus Surveillance.” *PLoS Biology* 22 (11): e3002916.

Developers, Inkscape Website. 2025. “Inkscape - Draw Freely.” May 12. https://inkscape.org.

Edgar, Robert C., and Henrik Flyvbjerg. 2015. “Error Filtering, Pair Assembly and Error Correction for next-Generation Sequencing Reads.” *Bioinformatics (Oxford, England)* 31 (21): 3476–3482.

Elbe, Stefan, and Gemma Buckland-Merrett. 2017. “Data, Disease and Diplomacy: GISAID’s Innovative Contribution to Global Health.” *Global Challenges (Hoboken, NJ)* 1 (1): 33–46.

Fowler, D. M., and S. Fields. 2014. “Deep Mutational Scanning: A New Style of Protein Science.” *Nature Methods* 11 (8). https://doi.org/10.1038/nmeth.3027.

Frank, Filipp, Meredith M. Keen, Anuradha Rao, et al. 2022. “Deep Mutational Scanning Identifies SARS-CoV-2 Nucleocapsid Escape Mutations of Currently Available Rapid Antigen Tests.” *Cell* 185 (19): 3603.

GitHub. n.d. “Prior Choice Recommendations.” Accessed July 11, 2025. https://github.com/stan-dev/stan/wiki/Prior-Choice-Recommendations.

Han, Alvin X., Simon P. J. de Jong, and Colin A. Russell. 2023. “Co-Evolution of Immunity and Seasonal Influenza Viruses.” *Nature Reviews. Microbiology* 21 (12): 805–817.

Hay, James A., Huachen Zhu, Chao Qiang Jiang, et al. 2024. “Reconstructed Influenza A/H3N2 Infection Histories Reveal Variation in Incidence and Antibody Dynamics over the Life Course.” *PLoS Biology* 22 (11): e3002864.

Hom, Nancy, Lauren Gentles, Jesse D. Bloom, and Kelly K. Lee. 2019. “Deep Mutational Scan of the Highly Conserved Influenza A Virus M1 Matrix Protein Reveals Substantial Intrinsic Mutational Tolerance.” *Journal of Virology* 93 (13). https://doi.org/10.1128/JVI.00161-19.

Iuliano, A. Danielle, Katherine M. Roguski, Howard H. Chang, et al. 2018. “Estimates of Global Seasonal Influenza-Associated Respiratory Mortality: A Modelling Study.” *Lancet (London, England)* 391 (10127): 1285–1300.

Jian, Fanchong, Jing Wang, Ayijiang Yisimayi, et al. 2024. “Evolving Antibody Response to SARS-CoV-2 Antigenic Shift from XBB to JN.1.” *Nature* 637 (8047): 921.

Kärber, G. 1931. “Beitrag zur kollektiven Behandlung pharmakologischer Reihenversuche.” *Naunyn-Schmiedeberg’s archives of pharmacology* 162 (4): 480–483.

Katoh, Kazutaka, John Rozewicki, and Kazunori D. Yamada. 2019. “MAFFT Online Service: Multiple Sequence Alignment, Interactive Sequence Choice and Visualization.” *Briefings in Bioinformatics* 20 (4): 1160–1166.

Khare, Shruti, Céline Gurry, Lucas Freitas, et al. 2021. “GISAID’s Role in Pandemic Response.” *China CDC Weekly* 3 (49): 1049–1051.

Koel, Björn F., David F. Burke, Theo M. Bestebroer, et al. 2013. "Substitutions near the Receptor Binding Site Determine Major Antigenic Change during Influenza Virus Evolution." *Science (New York, N.Y.)* 342 (6161): 976–979.

Krammer, Florian, Gavin J. D. Smith, Ron A. M. Fouchier, et al. 2018. "Influenza." *Nature Reviews. Disease Primers* 4 (1): 3.

Kumar, Ravin, Colin Carroll, Ari Hartikainen, and Osvaldo Martin. 2019. "ArviZ a Unified Library for Exploratory Analysis of Bayesian Models in Python." *Journal of Open Source Software* 4 (33): 1143.

Lei, Ruipeng, Andrea Hernandez Garcia, Timothy J. C. Tan, et al. 2023. "Mutational Fitness Landscape of Human Influenza H3N2 Neuraminidase." *Cell Reports* 42 (1): 111951.

Li, Chengjun, Masato Hatta, David F. Burke, et al. 2016. "Selection of Antigenically Advanced Variants of Seasonal Influenza Viruses." *Nature Microbiology* 1 (6): 16058.

Loes, Andrea N., Rosario Araceli L. Tarabi, John Huddleston, et al. 2024. "High-Throughput Sequencing-Based Neutralization Assay Reveals How Repeated Vaccinations Impact Titers to Recent Human H1N1 Influenza Strains." *Journal of Virology* 98 (10): e0068924.

Marco-Sola, Santiago, Jordan M. Eizenga, Andrea Guarracino, Benedict Paten, Erik Garrison, and Miquel Moreto. 2023. "Optimal Gap-Affine Alignment in O(s) Space." *Bioinformatics (Oxford, England)* 39 (2). https://doi.org/10.1093/bioinformatics/btad074.

Marco-Sola, Santiago, Juan Carlos Moure, Miquel Moreto, and Antonio Espinosa. 2021. "Fast Gap-Affine Pairwise Alignment Using the Wavefront Algorithm." *Bioinformatics (Oxford, England)* 37 (4): 456–463.

Miller, Rupert G. 1973. "Nonparametric Estimators of the Mean Tolerance in Bioassay." *Biometrika* 60 (3): 535.

Mögling, Ramona. 2016. "Evolution of Human Seasonal Influenza Viruses : Intrinsic and Extrinsic Fitness / Ramona Mögling." 2016.

Neumann, G., T. Watanabe, H. Ito, et al. 1999. "Generation of Influenza A Viruses Entirely from Cloned cDNAs." *Proceedings of the National Academy of Sciences of the United States of America* 96 (16): 9345–9350.

Nguyen, Alan B. H., Marco Bonici, Glen McGee, and Will J. Percival. 2025. "LOO-PIT: A Sensitive Posterior Test." *Journal of Cosmology and Astroparticle Physics* 2025 (01): 008.

Nunes-Correia, I., J. Ramalho-Santos, S. Nir, and M. C. Pedroso de Lima. 1999. "Interactions of Influenza Virus with Cultured Cells: Detailed Kinetic Modeling of Binding and Endocytosis." *Biochemistry* 38 (3): 1095–1101.

"Open Science Art." n.d. Accessed July 14, 2025. https://opensciencart.com/.

Pauly, Matthew D., Megan C. Procario, and Adam S. Lauring. 2017. *A Novel Twelve Class Fluctuation Test Reveals Higher than Expected Mutation Rates for Influenza A Viruses*. June 9. https://doi.org/10.7554/eLife.26437.

"Racmacs." n.d. Accessed December 1, 2025. https://acorg.github.io/Racmacs/index.html.

Smith, Derek J., Alan S. Lapedes, Jan C. de Jong, et al. 2004. “Mapping the Antigenic and Genetic Evolution of Influenza Virus.” *Science* 305 (5682): 371–376.

Šošić, Martin, and Mile Šikić. 2017. “Edlib: A C/C ++ Library for Fast, Exact Sequence Alignment Using Edit Distance.” *Bioinformatics* 33 (9): 1394–1395.

Spearman, C. 1908. “The Method of Right and Wrong Cases (Constant Stimuli) without Gauss’s Formulae.” *British Journal of Psychology* 2 (3): 227.

Spearman, C. 1909. “Review of the Method of ’right and Wrong Cases' ('constant Stimuli') without Gauss's Formula.” *Psychological Bulletin* 6 (1): 27–28.

Tenforde, Mark W., Rebecca J. Garten Kondor, Jessie R. Chung, et al. 2021. “Effect of Antigenic Drift on Influenza Vaccine Effectiveness in the United States-2019-2020.” *Clinical Infectious Diseases : An Official Publication of the Infectious Diseases Society of America* 73 (11): e4244–e4250.

Tureli, Sina. 2025a. “Merlign.” GitHub. https://github.com/iAvicenna/Merlign.

Tureli, Sina. 2025b. “Mitril.” GitHub. https://github.com/iAvicenna/Mitril.

Tureli, Sina. 2025c. “NGS-RFA/FRA Paper Repository.” GitHub. https://github.com/iAvicenna/ngs_rfa_fra_paper_repo.

Tureli, Sina. 2025d. “Thor.” GitHub. https://github.com/iAvicenna/Thor.

Turner, Samuel A., David J. Pattinson, Ron A. M. Fouchier, and Derek J. Smith. 2026. “Predicting the Antigenic Evolution of Seasonal Influenza Viruses Using Phylogenetic Convergence.” In *bioRxiv*. April 10. https://doi.org/10.64898/2026.04.10.717627.

Vehtari, Aki, Andrew Gelman, and Jonah Gabry. 2015. *Practical Bayesian Model Evaluation Using Leave-One-out Cross-Validation and WAIC*. July 16. https://doi.org/10.1007/s11222-016-9696-4.

Vehtari, Aki, Andrew Gelman, Daniel Simpson, Bob Carpenter, and Paul-Christian Bürkner. 2021. “Rank-Normalization, Folding, and Localization: An Improved Rˆ for Assessing Convergence of MCMC (with Discussion).” *Bayesian Analysis* 16 (2). https://doi.org/10.1214/20-ba1221.

Welsh, Frances C., Rachel T. Eguia, Juhye M. Lee, et al. 2024. “Age-Dependent Heterogeneity in the Antigenic Effects of Mutations to Influenza Hemagglutinin.” *Cell Host & Microbe* 32 (8): 1397–1411.e11.

Wilks, Samuel H., Barbara Mühlemann, Xiaoying Shen, et al. 2022. “Mapping SARS-CoV-2 Antigenic Relationships and Serological Responses.” In *bioRxiv*. January 28. https://doi.org/10.1101/2022.01.28.477987.

Yu, Yingpu, Maximilian A. Kass, Mengyin Zhang, et al. 2024. “Deep Mutational Scanning of Hepatitis B Virus Reveals a Mechanism for Cis-Preferential Reverse Transcription.” *Cell* 187 (11): 2735–2745.e12.
